# Fly Cell Atlas: a single-cell transcriptomic atlas of the adult fruit fly

**DOI:** 10.1101/2021.07.04.451050

**Authors:** Hongjie Li, Jasper Janssens, Maxime De Waegeneer, Sai Saroja Kolluru, Kristofer Davie, Vincent Gardeux, Wouter Saelens, Fabrice David, Maria Brbić, Jure Leskovec, Colleen N. McLaughlin, Qijing Xie, Robert C. Jones, Katja Brueckner, Jiwon Shim, Sudhir Gopal Tattikota, Frank Schnorrer, Katja Rust, Todd G. Nystul, Zita Carvalho-Santos, Carlos Ribeiro, Soumitra Pal, Teresa M. Przytycka, Aaron M. Allen, Stephen F. Goodwin, Cameron W. Berry, Margaret T. Fuller, Helen White-Cooper, Erika L. Matunis, Stephen DiNardo, Anthony Galenza, Lucy Erin O’Brien, Julian A. T. Dow, FCA Consortium, Heinrich Jasper, Brian Oliver, Norbert Perrimon, Bart Deplancke, Stephen R. Quake, Liqun Luo, Stein Aerts

**Affiliations:** Department of Biology, Howard Hughes Medical Institute, Stanford University, Stanford, CA 94305, USA; Huffington Center on Aging and Department of Molecular and Human Genetics, Baylor College of Medicine, Houston, TX 77030, USA; VIB-KU Leuven Center for Brain & Disease Research, KU Leuven, Leuven 3000, Belgium; Laboratory of Computational Biology, Department of Human Genetics, KU Leuven, Leuven 3000, Belgium; Departments of Bioengineering and Applied Physics, Stanford University, Stanford CA USA, and Chan Zuckerberg Biohub, San Francisco CA USA; Laboratory of Systems Biology and Genetics, Institute of Bioengineering, School of Life Sciences, Ecole Polytechnique Fédérale de Lausanne (EPFL) and Swiss Institute of Bioinformatics, CH-1015 Lausanne, Switzerland; Department of Computer Science, Stanford University, Stanford, CA 94305, USA; Department of Cell and Tissue Biology, University of California, San Francisco, CA 94143, USA; Department of Life Science, College of Natural Science, Hanyang University, Seoul, Republic of Korea 04763; Department of Genetics, Blavatnik Institute, Harvard Medical School, Harvard University, Boston, MA 02115; Howard Hughes Medical Institute, Boston, MA, USA; Aix-Marseille University, CNRS, IBDM (UMR 7288), Turing Centre for Living systems, 13009 Marseille, France; Institute of Physiology and Pathophysiology, Department of Molecular Cell Physiology, Philipps-University, Marburg, Germany; Department of Anatomy, University of California, San Francisco, CA 94143; Behavior and Metabolism Laboratory, Champalimaud Research, Champalimaud Centre for the Unknown, Lisbon, Portugal; National Center of Biotechnology Information, National Library of Medicine, NIH, Bethesda, MD 20894, USA; Centre for Neural Circuits & Behaviour, University of Oxford, Tinsley Building, Mansfield road, Oxford, OX1 3SR; Department of Developmental Biology, Stanford University School of Medicine, Stanford, CA 94305, USA; Department of Genetics, Stanford University School of Medicine, Stanford CA 94305, USA; Molecular Biosciences Division, Cardiff University, Cardiff, CF10 3AX UK; Department of Cell Biology, Johns Hopkins University School of Medicine, Baltimore, MD 21205 USA; Perelman School of Medicine, The University of Pennsylvania, and The Penn Institute for Regenerative Medicine Philadelphia, PA 19104 USA; Department of Molecular and Cellular Physiology, Stanford University School of Medicine, Stanford CA 94305 USA; Institute of Molecular, Cell & Systems Biology, College of Medical, Veterinary and Life Sciences, University of Glasgow, Glasgow G12 8QQ, UK; Immunology Discovery, Genentech, Inc., 1 DNA Way, South San Francisco, CA 94080, USA; Laboratory of Cellular and Developmental Biology, National Institute of Diabetes and Kidney and Digestive Diseases, National Institutes of Health, Bethesda, MD 20892, USA; see details of FCA Consortium below, and affiliations listed in the end of the document

**Author notes:** corresponding authors (N.P.), (B.D.), (S.R.Q.), (L.L.), (S.A.). equal contribution (Hongjie Li, Jasper Janssens). **FCA Consortium** **<u>Overall coordination</u>** Hongjie Li, Jasper Janssens, Norbert Perrimon, Bart Deplancke, Stephen R. Quake, Liqun Luo, Stein Aerts **<u>Logistical coordination</u>** Hongjie Li, Jasper Janssens, Maxime De Waegeneer, Sai Saroja Kolluru, Robert C. Jones, Norbert Perrimon, Bart Deplancke, Stephen R. Quake, Liqun Luo, Stein Aerts **Biohub**: Aaron McGeever, Angela Oliveira Pisco, Jim Karkanias, Sheela Crasta **Flybase**: Gil dos Santos, Clare Pilgrim, Alex McLachlan **EBI**: David Osumi-Sutherland, Irene Papatheodorou, Nancy George, Jonathan Manning **<u>Tissue dissection</u>** **Liqun Luo lab (head, body, antenna, haltere)**: Liqun Luo, Hongjie Li, Jiefu Li, David Vacek, Anthony Xie **Lucy O’Brien lab (gut)**: Lucy Erin O’Brien, Yu-Han Su, Erin Nicole Sanders, Samantha Gumbin, Paola Moreno-Roman, Aparna Sherlekar, Andrew Thomas Labott, Sang Ngo **Norbert Perrimon lab (Malpighian tubule)**: Norbert Perrimon, Ruei-Jiun Hung, Jun Xu **Yuh-Nung Jan lab (body wall)**: Yuh Nung Jan, Jacob S. Jaszczak, Ruijun Zhu, Ke Li, Yanmeng Guo, Kai Li, Liying Li, Tun Li, Han-Hsuan Liu, Caitlin E. O’Brien, Wanpeng Wang, Maja Petkovic **Rolf Bodmer lab (heart)**: Rolf Bodmer, Georg Vogler, Marco Tamayo, James Kezos, Katja Birker, Tanja Nielsen **Mariana Wolfner lab (male reproductive glands)**: Mariana F. Wolfner, Norene A. Buehner **Kristin Scott lab (leg, wing, proboscis & max palp)**, Kristin Scott, Amy Chang, Stefanie Engert, Amanda Gonzalez, Meghan Laturney, Sarah Leinwand, Carolina Reisenman, Philip Shiu, Gabriella Sterne, Zepeng Yao **Heinrich Jasper lab (fat body, oenocyte)**: Heinrich Jasper, Xiaoyu Tracy Cai, Nadja Sandra Katheder **Tom Kornberg lab (trachea)**: Thomas B Kornberg, Wanpeng Wang **Margaret Fuller lab (testis)**: Margaret T. Fuller, Neuza Reis Matias, Cameron W. Berry, Susanna E. Brantley, Catherine C. Baker, Devon E. Harris, Yiu-Cheung E. Wong, Benjamin Bolival **Todd Nystul lab (ovary)**: Todd G. Nystul, Katja Rust **Katja Brueckner lab (hemocyte)**: Katja Brueckner, Jordan Augsburger, Anjeli Mase **Seung Kim lab (insulin-producing cell, corpora cardiaca cell)**: Seung K. Kim, Lutz Kockel **<u>Library preparation and sequencing</u>** **Luo and Quake labs**: Hongjie Li, Sai Saroja Kolluru, Colleen N. McLaughlin, Qijing Xie, Robert C. Jones, Felix Horns **Biohub**: Angela M. Detweiler, Jia Yan, Michelle Tan, Norma Neff, Rene V. Sit **NIH group**: Harold E. Smith, Brian Oliver **<u>Data analysis</u>** Jasper Janssens, Maxime De Waegeneer, Kristofer Davie, Swann Floc’hlay, Stein Aerts, Vincent Gardeux, Wouter Saelens, Fabrice David, Maria Litovchenko, Bart Deplancke, Maria Brbić, Kumar Ayush, Hongjie Li, Jure Leskovec **<u>Case study analysis/writing</u>** **Common cell - hemocyte**: Katja Brueckner, Jiwon Shim, Sudhir Tattikota, Jasper Janssens, Hongjie Li **Common cell - muscle**: Frank Schnorrer, Jasper Janssens **Gut data integration**: Wouter Saelens **Metabolic pathway**: Carlos Ribeiro, Zita Carvalho-Santos, Darshan B. Dhakan, Rita Cardoso-Figueiredo, **Ovary data integration**: Katja Rust, Todd G. Nystul **Sex differences**: Brian Oliver, Soumitra Pal, Teresa Przytycka, Aaron M. Allen, Devika Agarwal, Stephen F Goodwin, Julian A. T. Dow **Testis annotation & trajectory**: Margaret T. Fuller, Cameron W. Berry, Erika L. Matunis, Stephen DiNardo, Helen White-Cooper, Brian Oliver, Sharvani Mahadevaraju, Julie A. Brill, Henry M. Krause, Wouter Saelens, Bart Deplancke **<u>Cell type annotation</u>** **SCope team**: Stein Aerts (PI), Jasper Janssens, Maxime De Waegeneer, Kristofer Davie **ASAP team**: Bart Deplancke (PI), Vincent Gardeux, Wouter Saelens, Fabrice David **Gut**: Lucy Erin O’Brien, Anthony Galenza, Aparna Sherlekar, Erin Nicole Sanders, Yu-Han Su, Anna A. Kim, Kazuki Yoda, Norbert Perrimon, Joshua Shing Shun Li **Malpighian tubule**: Norbert Perrimon, Julian A. T. Dow, Jun Xu, Jasper Janssens, Yifang Liu **Body wall**: Yuh Nung Jan, Jacob S. Jaszczak, Ruijun Zhu, Ke Li, Yanmeng Guo, Liying Li, Hongjie Li **Heart**: Rolf Bodmer, Georg Vogler, Hongjie Li, Jasper Janssens **Muscle**: Frank Schnorrer, Jasper Janssens **Male reproductive glands**: Mariana F. Wolfner, Nora C. Brown, Yasir Ahmed-Braimah, Helen White-Cooper, Mikaela Matera-Vatnick, Timothy L. Karr **Leg, wing, prob. and max palp**: Kristin Scott, Zepeng Yao, Carlos Ribeiro, Ibrahim Tastekin **Antenna and haltere**, Liqun Luo, Hongjie Li, Colleen N. McLaughlin, Andrew K. Groves, Shinya Yamamoto, Daniel Sutton, Rachel I. Wilson, Stephen L. Holtz **Fatbody and oenocyte**: Heinrich Jasper, Xiaoyu Tracy Cai, Nadja Sandra Katheder, Sudhir Gopal Tattikota, Carlos Ribeiro, Zita Carvalho-Santos, Rita Cardoso-Figueiredo, Hongjie Li, Jasper Janssens Trachea: Thomas B Kornberg, Wanpeng Wang, Hongjie Li **Testis (see case study)** **Ovary**: Todd G. Nystul, Katja Rust, Ruth Lehman, Maija Slaidina, Torsten Banisch, Zita Carvalho-Santos, Mariana F. Wolfner, Wanpeng Wang, Brian Oliver, Sharvani Mahadevaraju **Hemocyte**, Katja Brueckner, Jiwon Shim, Sudhir Gopal Tattikota, Jasper Janssens, Hongjie Li **Insulin-producing cell (IPC) and corpora cardiaca cell (CC)**: Seung K. Kim, Lutz Kockel, Maria Brbić **Head and body**: Aaron M. Allen, Bruno Hudry, Caitlin E. McDonough-Goldstein, Christoph Treiber, Clare Pilgrim, Claude Desplan, David Sims, Devika Agarwal, Erika Donà, Fernando Casares, Gregory S.X.E. Jefferis, Majd M. Ariss, Megan Neville, Michelle Arbeitman, Mehmet Neset Özel, Nikolaos Konstantinides, Scott Waddell, Stephen F Goodwin, Thomas R. Clandinin, Yael Heifetz, Jasper Janssens, Hongjie Li, Stein Aerts **<u>Writing group</u>** Hongjie Li, Jasper Janssens, Norbert Perrimon, Bart Deplancke, Stephen R. Quake, Liqun Luo, Stein Aerts **<u>Principal investigators (A-Z):</u>** Stein Aerts, Yasir Ahmed-Braimah, Rolf Bodmer, Julie A. Brill, Katja Brueckner, Fernando Casares, Thomas R. Clandinin, Bart Deplancke, Claude Desplan, Stephen DiNardo, Julian A. T. Dow, Margaret T. Fuller, Stephen F Goodwin, Andrew K. Groves, Yael Heifetz, Bruno Hudry, Yuh Nung Jan, Heinrich Jasper, Gregory S.X.E. Jefferis, Timothy L. Karr, Seung K. Kim, Nikolaos Konstantinides, Thomas B Kornberg, Henry M. Krause, Jure Leskovec, Hongjie Li, Liqun Luo, Erika L. Matunis, Todd G. Nystul, Lucy Erin O’Brien, Brian Oliver, Norbert Perrimon, Teresa M Przytycka, Stephen R. Quake, Carlos Ribeiro, Katja Rust, Frank Schnorrer, Kristin Scott, Jiwon Shim, Helen White-Cooper, Rachel I. Wilson, Mariana F. Wolfner, Shinya Yamamoto.

## Abstract

The ability to obtain single cell transcriptomes for stable cell types and dynamic cell states is ushering in a new era for biology. We created the Tabula *Drosophilae*, a single cell atlas of the adult fruit fly which includes 580k cells from 15 individually dissected sexed tissues as well as the entire head and body. Over 100 researchers from the fly community contributed annotations to >250 distinct cell types across all tissues. We provide an in-depth analysis of cell type-related gene signatures and transcription factor markers, as well as sexual dimorphism, across the whole animal. Analysis of common cell types that are shared between tissues, such as blood and muscle cells, allowed the discovery of rare cell types and tissue-specific subtypes. This atlas provides a valuable resource for the entire *Drosophila* community and serves as a comprehensive reference to study genetic perturbations and disease models at single cell resolution.

## Main Text

The fruit fly *Drosophila melanogaster* has a long and fruitful history in biological research, dating back to the first experiments of Thomas Hunt Morgan in the early 20th century (Morgan, 1910). *Drosophila* has been at the basis of many key discoveries in genetics, development, neurobiology, circadian rhythms, stem cell biology, and aging (Bellen et al., 2010). The highly collaborative nature of the *Drosophila* community contributed to many of these successes, and led to the development of essential research resources, including a high-quality genome (Adams et al., 2000), a large collection of mutant lines and a large variety of genetic and molecular tools, as well as important databases such as Flybase (Larkin et al., 2021), FlyMine (Lyne et al., 2007), FlyLight (Jenett et al., 2012), VirtualFlyBrain (Milyaev et al., 2012) and ModERN (Kudron et al., 2018). The fruit fly genome contains about 17,000 genes, including an estimated 13,968 protein-coding genes of which ∼63% have human orthologues. Studies such as ModENCODE (modENCODE Consortium et al., 2010) and FlyAtlas (Chintapalli et al., 2007) have explored their expression patterns in embryonic, larval, pupal, and adult tissues, but have lacked cell type resolution. Recent advances in single cell technologies have enabled the transcriptomic profiling of thousands of cells at once, leading to greater insight in tissue composition and differential gene expression. Several studies have already applied single cell RNA sequencing (scRNA-seq) on *Drosophila* tissues, including embryo, imaginal discs, brain, ventral nerve cord, gut, blood, antenna, abdominal cuticle, ovary and testis (Li, 2020; Mohr et al., 2021). However, these data have been generated by different laboratories on different genetic backgrounds, with different dissociation protocols and sequencing platforms, hindering systematic comparison of gene expression across cells and tissues.

Here, we present a single cell transcriptomic atlas of the entire adult *Drosophila,* separately analyzing male vs female samples, using a uniform genotype and a unified single-nucleus RNA-seq (snRNA-seq) platform (McLaughlin et al., 2021) with two sequencing strategies: droplet-based 10x Chromium and plate-based Smart-seq2. The resulting Tabula *Drosophilae*, the first comprehensive dataset within the Fly Cell Atlas consortium, contains over 580k cells, resulting in more than 250 distinct cell types that were annotated by over 100 experts from 40 international *Drosophila* research laboratories. This atlas reports cellular signatures for each tissue, providing a valuable resource for the entire *Drosophila* community as a reference for studies that probe the effects of particular genetic perturbations and disease models at single cell resolution. All data and annotations can be accessed through multiple visualization and analysis portals from https://flycellatlas.org.

### Sampling single cells across the entire adult fly

To achieve a comprehensive sampling of cell types in the adult fruit fly, we used two complementary strategies. In the first strategy, we dissected 12 individual tissues from both males and females, and 3 additional sex-specific tissues (see Methods). Tissues were frozen, followed by nuclei extraction, staining, and fluorescence-activated cell sorting (FACS). Next, sorted nuclei were collected into one tube or 384-well plates to perform snRNA-seq using 10x Genomics (Zheng et al., 2017) and Smart-seq2 (Picelli et al., 2013), respectively (**Fig. 1A**). For three tissues that are localized across the body and cannot be directly dissected, we used specific GAL4 lines driving nuclear-GFP to label these nuclei and collected them using FACS based on GFP fluorescence signal: *Cg-GAL4* for fat body, *PromE800-GAL4* for oenocytes, and *Btl*-GAL4 for trachea (see Methods). In addition, we sequenced two more rare cell types using Smart-seq2: insulin-producing cells (IPCs, *Dilp-GAL4*) and corpora cardiaca cells (CCs, *Akh-GAL4*). In the second strategy, we sorted nuclei from the entire head and body of sexed wild-type flies and profiled them on 24 10x Chromium lanes, aiming to detect cell types that may not be covered by the selected tissues. The libraries were sequenced using Illumina NovaSeq 6000 in 4 batches, for a total of 38 billion reads. In total, we obtained 580k high-quality cells: 570k from 10x Genomics of 63 runs and 10k from Smart-seq2 of 35 384-well plates (**Fig. 1A)**.

**Figure 1.**
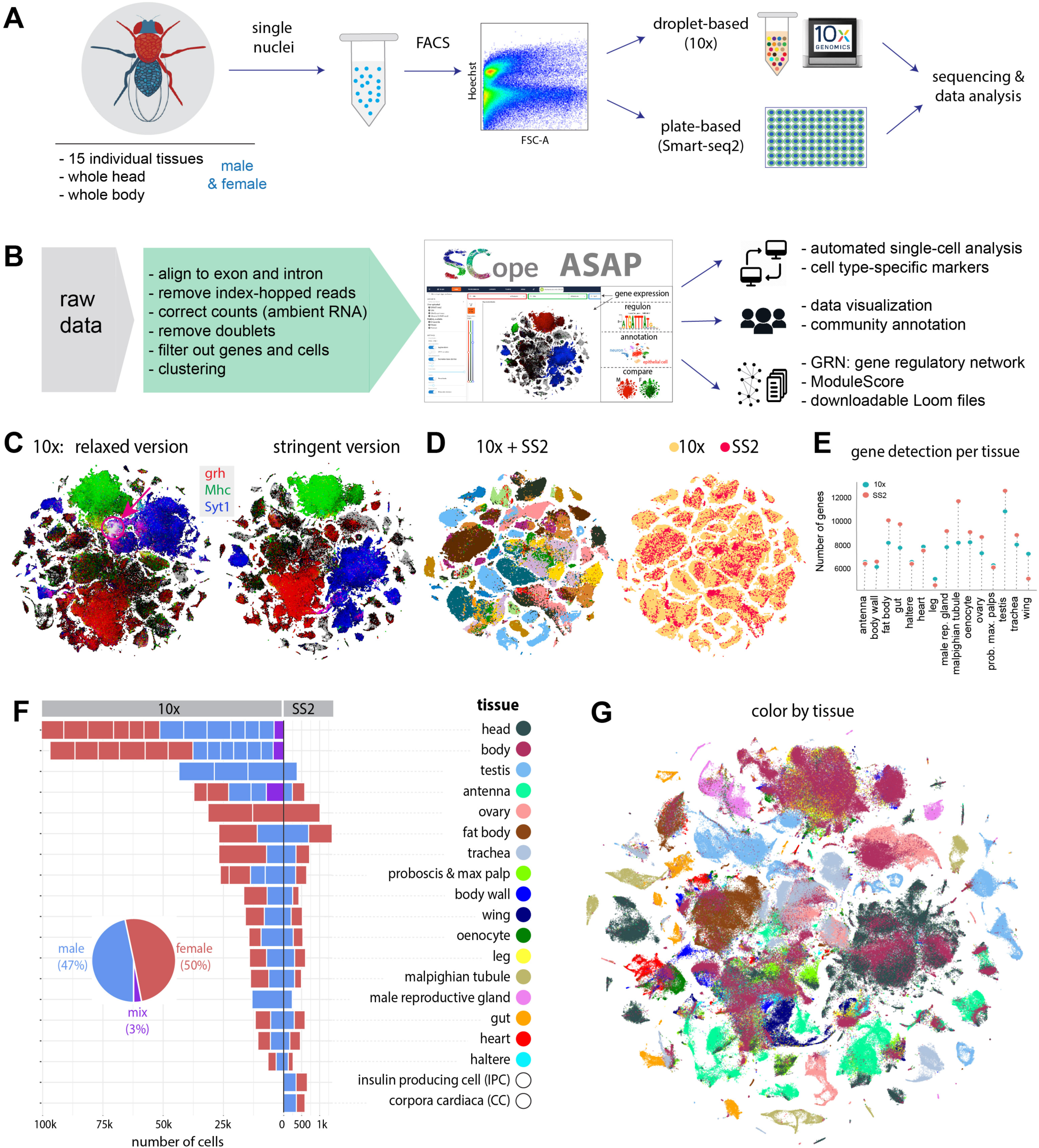
**Overview of the Fly Cell Atlas** **(A)** Experimental platform of single-nucleus RNA-seq (snRNA-seq) using 10x Genomics and Smart-seq2. **(B)** Data analysis pipeline and data visualization using SCope <u>(Davie et al., 2018)</u> and ASAP <u>(David et al., 2020)</u>. **(C)** Two versions of 10x datasets have been generated: a *Relaxed* and a *Stringent* version. tSNE colors based on gene expression: *grh* (epithelia, red), *Mhc* (muscle, green) and *Syt1* (neuron, blue). Red arrow highlights artefactual cluster with co-expression of all three markers in the *Relaxed* dataset. **(D)** tSNE visualization of cells from the *Stringent* 10x dataset and Smart-seq2 (SS2) cells. 10x cells are from individual tissues. Integrated data is colored by tissue (left) and technology (right). **(E)** Tissue-level comparison of the number of detected genes between 10x and Smart-seq2 technologies (see Methods). **(F)** Number of cells for each tissue by 10x and Smart-seq2. Male and female cells are indicated. Mixed cells are from pilot experiments where flies were not sexed. Different batches are separated by vertical white lines **(G)** All 10x cells from the *Stringent* dataset clustered together; cells are colored by tissue type. Tissue names and colors indexed in F.

To analyze the 10x Genomics data in a reproducible manner, we used an automated computational pipeline called VSN (https://doi.org/10.5281/zenodo.3703108; see Methods), which takes the raw sequencing data as input, performs filtering, normalization, doublet removal, batch effect correction, dimensionality reduction, and clustering, and produces *LoomX* formatted files with expression data, embeddings and clusterings (**Fig. 1B** and **fig. S1**). As we observed a presumed artefactual cluster of cells showing expression of nearly all genes, we added an additional preprocessing and filtering step that models and then subtracts ambient RNA from the gene expression values (Yang et al., 2020), resulting in computational removal of this cluster. However, since adjusting the gene expression values per cell can also introduce other biases (e.g., overcorrection, removal of non-doublet cells), we retained both a *Relaxed* dataset of 570k cells, and a *Stringent* dataset of 510k cells (see Methods and **Fig. 1C**). In the analyses below, we focus on the *Stringent* dataset, unless otherwise mentioned (e.g., Fig. 2C).

**Figure 2:**
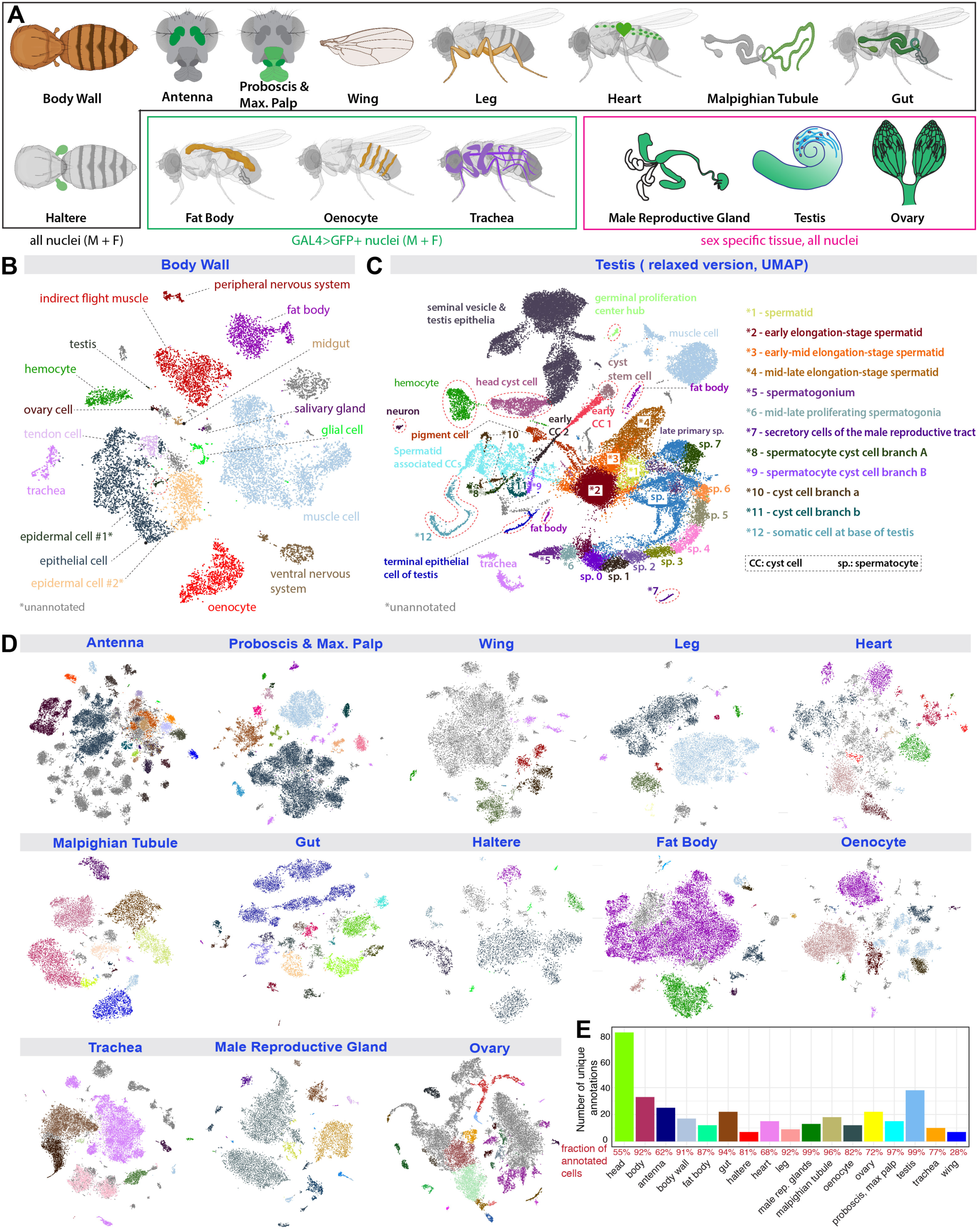
**Cell type annotation for dissected tissues** **(A)** Illustration of 15 individual tissues. 12 sequenced separatly from males and females, 3 sex specific. Fat body, oenocyte and tracheal nuclei were labeled using a tissue-specific GAL4 driving UAS-nuclearGFP. **(B)** tSNE plot with annotations for body wall from the *Stringent* 10x dataset. *1, epidermal cells of the abdominal posterior compartment. *2, epidermal cells specialized in antimicrobial response. **(C)** UMAP plot with annotations for the testis from the *Relaxed* 10x dataset. See detailed description in the Supplemental text. **(D)** tSNE plots of the other 13 tissues from the *Stringent* 10x dataset. Detailed annotations are in fig. S3– S15. **(E)** Number of unique annotations for each tissue. Fractions of annotated cells over all analyzed cells from the relaxed dataset are indicated in red.

Cells from 10x Genomics and Smart-seq2 were well aligned after batch correction using Harmony (Korsunsky et al., 2019) (**Fig. 1D**). As expected, smart-seq2 yielded a higher number of detected genes for most tissues (**Fig. 1E**), because Smart-seq2 cells were sequenced to a higher depth (∼500k reads per cell for Smart-seq2 versus ∼50k reads per cell for 10x). We analyzed each tissue separately, combining the male and female runs, which yielded between 6.5k (haltere) and 100k (head) cells per tissue with a median of 16.5k cells per tissue for 10x Genomics and between 263 (male reproductive gland) and 1,349 (fat body) cells per tissue with a median of 534 cells for Smart-seq2 (**Fig. 1F**). We obtained a similar number of male and female cells for non-sex-specific tissues. We retrieved on average 1895 UMIs and 828 genes per cell with a variation between tissues (**fig. S2**), suggesting that different tissues have different numbers of active genes, similar to other atlas data, such as Tabula Muris (Tabula Muris Consortium et al., 2018). Next, all cells were combined together in a meta-analysis, showing tissue-specific clusters like the germline cells of the testis and ovary, and shared clusters of common cell types, such as neurons, glial cells, fat cells, epithelial cells, hemocytes, and muscle cells from different tissues (**Fig. 1G,** also see fig. S21, 22).

### Crowd-based cell type annotation by tissue experts

Next, we focused on cell type annotation for 15 individual tissues, including 12 tissues for both males and females: antenna, body wall, fat body, haltere, heart, gut, leg, Malpighian tubule, oenocyte, proboscis with maxillary palp, trachea, and wing; and 3 additional sex-specific tissues: male reproductive gland, testis, and ovary (**Fig. 2A**). To allow experts from different international laboratories to participate in the annotation, we developed a consensus-voting strategy within the SCope web application (https://flycellatlas.org/scope) (Davie et al., 2018), allowing curators and users to access and annotate the snRNA-seq data simultaneously. We hosted a series of virtual ‘jamborees’, where tissue experts worked with the SCope team to annotate all clusters at multiple resolutions, with additional analysis performed in ASAP (https://flycellatlas.org/asap) (David et al., 2020). If the same cluster was annotated with two terms, the voting system allows users to vote for their preferred annotation. To ensure that cell type annotations are consistent with previous literature and databases and to allow *a posteriori* computational analyses at different anatomical resolutions, we linked the controlled vocabulary to the Flybase anatomy ontology (Costa et al., 2013).

Since some cell types are present at low clustering resolution, and others appear only at higher resolution, we used the VSN pipeline to calculate different Leiden resolutions ranging from 0.8 to 8 (**fig. S3A**). Annotations were then collapsed across resolutions and the annotation with the highest number of up-votes was selected per cell. All initial annotations were performed on the *Relaxed* dataset, and were then exported to the *Stringent* dataset, where field experts verified the accuracy of the annotation transfer (**Fig. 2 and fig. S3-S15**). Overall, such efforts led to the annotation of 251 cell types in the *Stringent* dataset (264 cell types if combining *Relaxed* and *Stringent datasets*), with a median of 15 cell types per tissue.

While single cell datasets exist for a number of fly tissues, our data yield two further advances. First, our dataset provides the first single cell transcriptomic profiling for several adult tissues, including the haltere, heart, leg, Malpighian tubule, proboscis, maxillary palp, trachea, and wing (**fig. S3–S15**). In these tissues, all major expected cell types were identified. For example, in the proboscis and maxillary palp (**fig. S4A, B**), we could annotate gustatory and olfactory receptor neurons, mechanosensory neurons and several glial clusters, and all 7 olfactory receptors expressed in the maxillary palp could be detected. In the wing (**fig. S5**), we could identify four different neuronal types – gustatory receptor neurons, pheromone-sensing neurons, nociceptive neurons and mechanosensory neurons, as well as three glial clusters. In the leg (**fig. S6**), we could distinguish gustatory receptor neurons from two clusters of mechanosensory neurons. In the heart (**fig. S7**), we found a large proportion of resident hemocytes and muscle cells, with the cardial cells marked by the genes *Hand* and *tinman* constituting a small proportion. In the Malpighian tubule (**fig. S8**), we were able to identify 15 cell types, including the different principal cells of the stellate and main segments. In the haltere (**fig. S10**), we identified two clusters of neurons, three clusters of glial cells and a large population of epithelial cells. In some tissues, one cell type formed a big cluster instead of being split into distinct populations. In these cases, we tried to identify genes or pathways that showed gradient or compartmentalized expression. For example, in the fat body (**fig. S16**), the main fat body cells formed one big cluster, but our metabolic pathway enrichment analysis performed through ASAP (David et al., 2020) revealed that fatty acid biosynthesis and degradation are in fact compartmentalized, highlighting possible fat body cell heterogeneity in terms of metabolic capacities.

Second, our crowd annotations with tissue experts revealed many cell types not profiled previously, such as multinucleated muscle cells (**Fig. 2B**), and uncovered some surprising new features. For example, in the testis (**Fig. 2C**), one of the best characterized but also most complex tissues in the fly, our FCA data allowed identification of 25 unique cell types or subtypes, covering all expected cell types, including very rare cells, such as germinal proliferation center hub cells (79 nuclei in the Relaxed version, out of 44,621 total testis nuclei). New features that emerged from the testis data include the stratification of later stage spermatocytes into three streams, all of which express diagnostic spermatocyte-specific markers, but vary in terms of overall expression levels for second chromosome or third chromosome-linked genes. In addition to the annotation of differentiated cell types, the snRNA-seq data captured the dynamic sequence of two stem cell lineages in the testis, the germ line lineage and somatic cyst cell lineage, identifying many dynamic genes with altered expression at specific steps during stem cell differentiation (see trajectory analyses in **Fig. 6**, and a more detailed discussion in the Supplementary text). Clustering at higher resolution (Leiden 6) was especially useful in these stem cell lineages as it highlighted as an emergent property how clusters of nuclei closely related in terms of expressed genes nevertheless turned on or off key genes as the developmental sequence advanced. Another surprising finding was the existence of two distinct types of nuclei among main cells in the male accessory gland (**fig. S14**), a cell type previously thought to be uniform.

Next, we compared the distribution of cells between 10x and Smart-seq2 for all tissues profiled by both sequencing technologies. In almost all cases, cells from two technologies were well matched from co-clustering analysis (**fig. S17 and S18**). Since Smart-seq2 cells only account for a small fraction, our previous annotations focused on 10x cells. The cell-matched co-clustering analysis allowed us to directly transfer annotations from 10x to Smart-seq2 datasets (**fig. S17E**), validating the transfer using cluster-specific markers (**fig. S17F**). We also identified a number of genes specifically detected using Smart-seq2, presumably because the deeper sequencing allowed a higher gene detection rate (**fig. S17G** and **Fig. 1E**). In summary, the 10x datasets, containing a large number of cells, provide an important resource for identifying cell types while the Smart-seq2 datasets facilitate the detection of low-expressed genes and enable future exploration of cell-specific isoform information.

### Correspondence between dissected tissues and whole head and body

To generate a complete atlas of the fly, we performed snRNA-seq experiments on whole-head and whole-body samples, inspired by earlier work on whole-body single cell experiments of less complex animals (Cao et al., 2017; Levy et al., 2021). Full head and body sequencing can provide a practical means to assess the impact of mutations or to track disease mechanisms, without having to focus on, or dissect, specific tissues. In addition, such a strategy could yield cell types that are not covered by any of the targeted tissue dissections.

First, we analyzed the head and body separately. In the head, we annotated 81 unique cell types with the majority of cells being neurons, as expected (**Fig. 3A and S19**). We identified photoreceptors and cone cells based on *gl* and *lz* expression respectively, olfactory receptor neurons with *orco* expression, glial cells based on *repo* expression, head hemocytes based on *Hml* expression, and different types of muscle cells and epithelial cells. In the body, we annotated the top 33 most abundant cell types, including epithelial and muscle groups, and ventral nerve cord and peripheral neurons, followed by fat body cells, oenocytes, germ line cells, glial cells, and tracheal cells (**Fig. 3B and S20**). Many of these broad cell types can be further divided into subtypes for further annotation, similar to that from tissue sequencing (see **Fig. 2** and **fig. S3-S15**).

**Figure 3:**
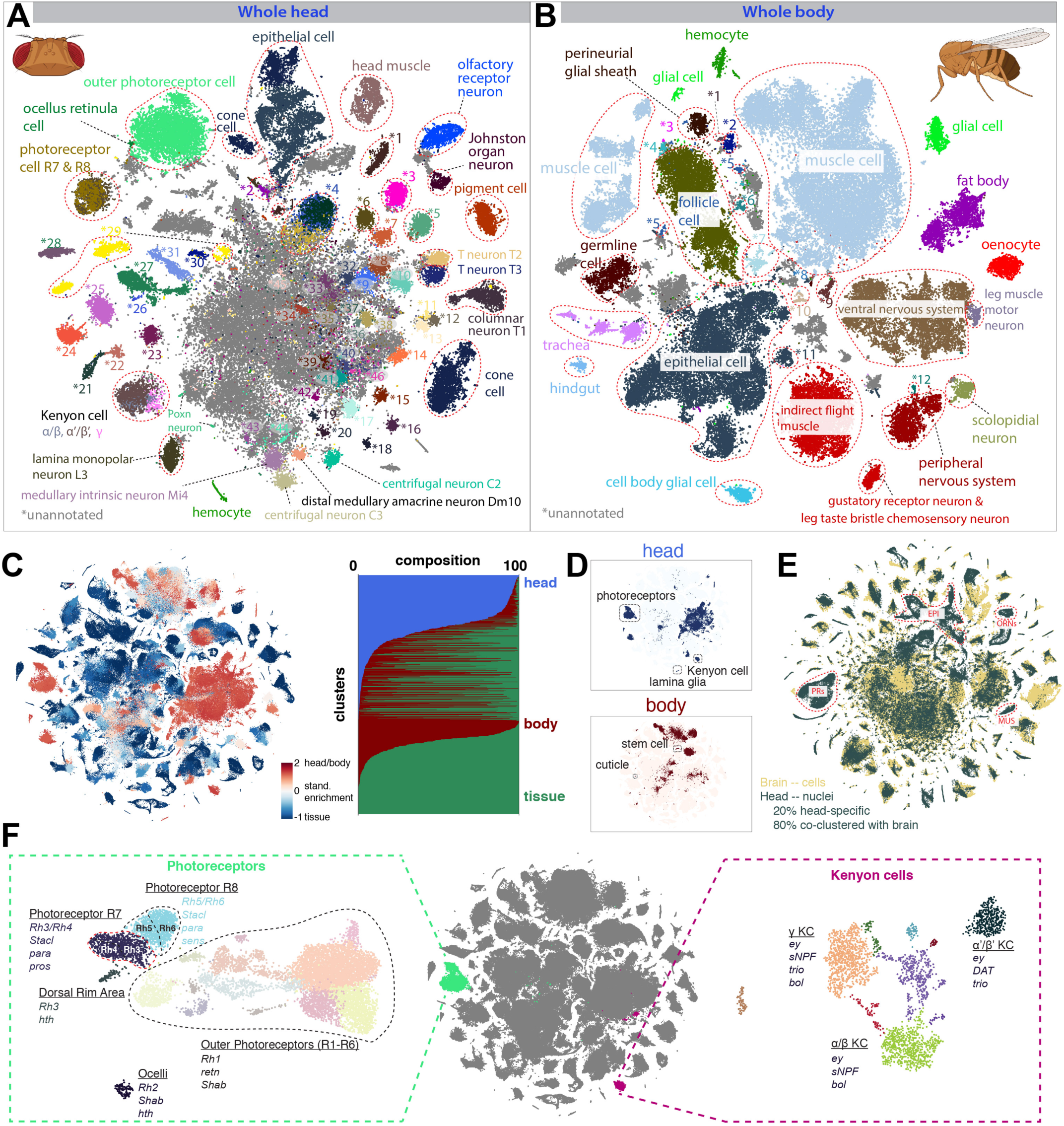
**Whole head and whole body sequencing leads to full coverage of the entire fly** **(A)** tSNE of the whole head sample with 81 annotated clusters. See fig. S18 for full cell types. A large number of cells in the middle (gray) are unannotated cells, most of which are neurons from the central brain. **(B)** tSNE of the whole body sample with 33 annotated clusters, many of which can be further divided into sub-clusters. Cells in gray are unannotated cells. See fig. S19 for full cell types. **(C)** (left) tSNE of the entire dataset colored by standardized tissue enrichment, leading to the identification of head- and body-specific clusters. (right) Barplots showing tissue composition for different clusters at Leiden resolution 50. **(D)** Examples of head- and body-specific clusters. **(E)** Integration of a brain scRNA-seq dataset with the head snRNA-seq for label transfer. Outlined are example clusters revealed by the head snRNA-seq dataset but not by the brain scRNA-seq datasets, including epithelial cells (EPI), photoreceptors (PRs), olfactory receptor neurons (ORNs), and muscle cells (MUS). **(F)** Subclustering analysis reveals subtypes of photoreceptors, including inner and outer photoreceptors, with the inner photoreceptors further splitting into R7 and R8 subtypes, and mushroom body Kenyon cells, forming three distinct KC subtypes: α/β, α’/β’ and γ.

Next, we examined how well the body and head samples covered the cell types from the dissected tissues. Analyzing the head and body samples together with the tissue samples revealed that most of the selected tissues cluster together with the body sample, as expected. We also detected head and body enriched clusters (**Fig. 3C**) and even clusters without cells from the dissected tissues (**Fig. 3D**). One had high expression of cuticular genes (p=1.35E-35), indicating a role in connective tissue. Others were relatively rare cell types in their respective tissues, such as adult stem cells. Conversely, most tissue clusters contained body cells, with only a small number being completely specific to dissected tissues. As tissue-specific clusters were mostly observed in tissues with high cell coverage, such as the testis and Malpighian tubule, we anticipate that these clusters would also be identified in the body upon sampling a larger number of cells.

For head, antenna and proboscis with maxillary palp were dissected for tissue sequencing. Cell types from those two tissues largely overlapped with head sequencing data. Many other cell types, such as central brain cells, including Kenyon cells (*ey*, *prt*) and lamina glia (*repo*, *Optix*), were only detected in the head sample.

Next, we integrated our head snRNA-seq dataset with previously published single cell RNA-seq data from the brain (Davie et al., 2018; Kurmangaliyev et al., 2020; Li et al., 2017; Özel et al., 2021) to automatically annotate our head cells (**Fig. 3E**). Head-specific clusters made up 20% of the cells, including the antennae, photoreceptors, muscle, cone cells and cuticular cell types, while the other 80% were present in mixed clusters containing both head- and brain-derived cells covering the neuronal and glial cell types of the brain. This co-clustering across genotypes and protocols underscores the usefulness of our snRNA-seq data, as well as its quality compared to more conventional scRNA-seq data. Next, we used a neural network for optic lobe cell types and trained a support-vector machine on a whole brain sample to predict annotations per cluster that were manually checked in an additional Jamboree. Given the high number of neuronal cell types, additional automatic subclustering was performed on each cluster, identifying different groups of peptidergic neurons based on *dimm* and *Pdf* expression and olfactory projection neurons based on *oaz*, *c15* and *kn*. Finally, we identified a large number of optic lobe cell types, including cell types of the lamina (e.g., L1–L5), medulla (e.g., Mi1, Mi15), lobula (e.g., LC) and lobula plate (e.g., LPLC). Using *acj6* and *SoxN*, we identified the T4/T5 neurons of the optic lobe that split in T4/T5a-b and T4/T5c-d subtypes by subclustering. Lastly, we identified the major subtypes of glia, leading to a total of 81 cell types in the head. It is worth noting that a large number of neuronal cells cluster together into a big clump that we were unable to annotate (**Fig. 3A**), indicating that our dataset does not contain a sufficient number of cells to resolve the complexity of the central brain, which may contain hundreds or thousands of neuronal types.

Subclustering analysis performed on cells within larger groups was especially useful in identifying subtypes in the combined dataset (**Fig. 3F**). This separated photoreceptors into inner and outer, with the inner photoreceptors further splitting into R7 and R8 subtypes, each with pale and yellow subtypes based on *rhodopsin* expression. Additionally, subclustering identified dorsal rim area and ocellar photoreceptors and further split Kenyon cells into the three different KC subtypes: α/β, α’/β’ and γ based on known marker genes (Davie et al., 2018). These case studies highlight the resolution in our dataset and the potential of using subclustering to discover rare cell types.

### Cross-tissue analyses allow comparison of cell types by location

Using the whole body and head sequencing data, we created a compilation of the entire fly, assigning cells to major broad cell types (e.g. epithelial cells, neurons, muscle cells, hemocytes) using a hierarchy of ontology terms and marker gene expression (**Fig. 4A-C**). Epithelial cells and muscle cells were the most abundant cells in the body, while neurons were the most abundant in the head. These assignments allowed us to pool common cell types across tissues for more in depth study (**fig. S21, S22**).

**Figure 4:**
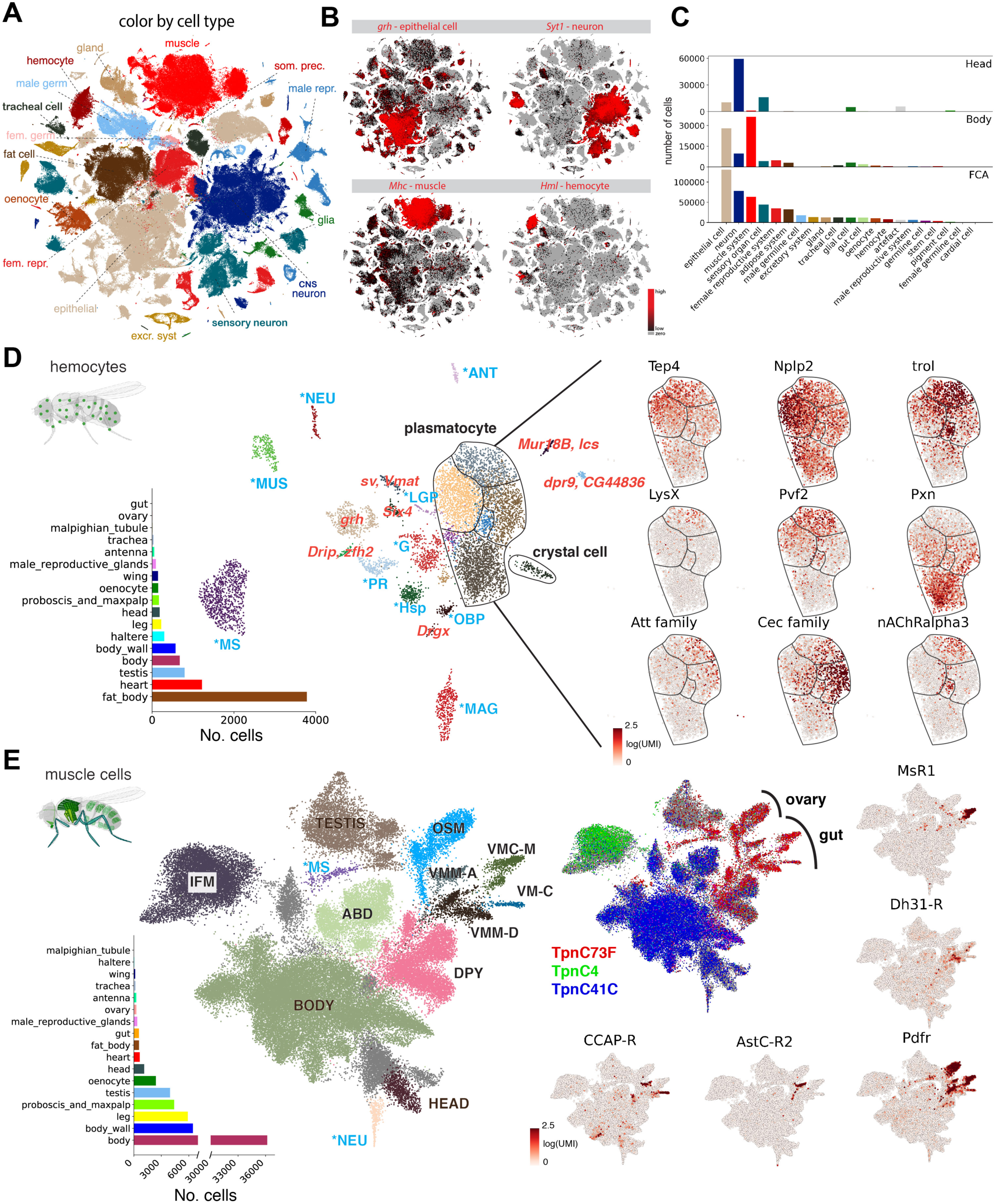
**Cross-tissue analyses of common cell types** **(A)** Overview of the different main cell classes identified throughout the fly cell atlas. Som. pre. for somatic precursor cells; male repr. for male reproductive system; fem. repr. for female reproductive system; male germ. for male germline cells; fem. germ. for female germline cells. **(B)** tSNE plots showing the expression of markers of four common cell types: *grh* for epithelial cells, *Syt1* for neuronal cells, *Mhc* for muscle cells and *Hml* for hemocytes. **(C)** Composition of whole head and body samples, showing a shift from neurons to epithelial and muscle cells. Composition of the entire fly cell atlas shows enrichment for rarer cell classes compared to the whole body sample. The barplot shows the number of cells. **(D)** Cross-tissue analysis of hemocytes reveals different cell states of plasmatocytes. Annotations marked as blue are hemocytes containing markers of different cell types, including lymph gland posterior signaling center (LGP), muscle markers (MUS), antennal markers (ANT), neuronal markers (NEU), photoreceptor markers (PR), male accessory glands makers (MAG), glial markers (G), male testis and spermatocyte markers (MS), olfactory-binding proteins (OBP), heat-shock proteins (Hsp). Other abbreviations show top marker gene(s) in red. Plasmatocytes and crystal cells are indicated. On the right are genes showing compartmentalized expression patterns within the plasmatocyte cluster. **(E)** Cross-tissue analysis of muscle cells reveals subdivision of the visceral muscle cells based on neuropeptide receptors. Annotations marked as blue are muscle cells containing markers of different cell types, including neuronal markers (NEU) and male testis and spermatocyte markers (MS). Muscle cells from three body parts are indicated: head muscle (HEAD), body muscle (BODY), and testis muscle (TESTIS). Other annotated muscle types include indirect flight muscle (IFM), ovarian sheath muscle (OSM), abdominal visceral muscle (ABD), *dpy* expressing muscle (DPY), visceral muscle of the midgut *AstC-R2* (VMM-A), visceral muscle of the crop *MsR1* (VMC-M), visceral muscle of the midgut *Dh31-R* (VMM-D), and visceral muscle *CCAP-R* (VM-C). *Pdfr* expressed in all visceral muscle cells, including the ovarian sheath muscle; other four receptor genes (*AstC-R2, MsR1, Dh31-R, CCAP-R*) expressed in different subtypes of gut visceral muscles.

First, we compared blood cells, termed hemocytes in the fly, across tissues by selecting all *Hml*-positive cells, a known marker for hemocytes (**Fig. 4D**). Hemocytes are generated in the embryo and larva and become resident cells in many tissues. Combining hemocytes from across tissues revealed a major group of plasmatocytes, the most common hemocyte type (∼56%), crystal cells (1.5%, PPO1, PPO2), and several unknown subtypes (**fig. S23A, B**). Looking deeper into the plasmatocytes, we uncovered different gradients of gene expression that were not seen in the plasmatocyte clusters of individual tissues. This gradient is based on the expression of *Pxn*, *LysX*, *Tep4*, *trol* and *Nplp2* and can be linked to maturation of hemocytes and their plasticity with *Pxn* positive cells showing the highest *Hml* expression, while *Tep4*, *trol* and *Nplp2* are prohemocyte markers (Cho et al., 2020; Grigorian et al., 2013; Makki et al., 2010). Furthermore, different antimicrobial peptide (AMP) families such as the *Attacins* and *Cecropins* were expressed in different subgroups of plasmatocytes, indicating further specialisation. Finally, expression of acetylcholine receptors was specific for a subset of hemocytes, which could be related to the cholinergic anti-inflammatory pathway as described in humans and mice (Gallowitsch-Puerta and Tracey, 2005; Pavlov and Tracey, 2005). Lamellocytes were not observed in adults as previously suggested (Sanchez Bosch et al., 2019). On the other hand, one of the unknown hemocyte subtypes expressed *Antp* and *kn* (43 cells, 0.5%) reminiscent of the posterior signaling center in the lymph gland, an organization center previously thought to be absent in the adult (Krzemień et al., 2007; Mandal et al., 2007) (**Fig. S23B**). These surprising findings highlight the value of performing a whole organism-level single cell analysis and constitute a great foundation to investigate the fly immune system in greater detail in the near future.

Second, we compared the muscle cell types of the different tissues (**Fig. 4E, S23C**). Muscle cells are syncytia, single cells containing many nuclei, and to our knowledge have not been profiled by single cell sequencing prior to our study. With snRNA-seq, we recovered all known muscle cell types, with specific enrichment in the body, body wall, and leg. The integrated analysis of all muscle cells provided a comprehensive view of the fly muscular system, highlighting separation of visceral, skeletal, and indirect flight muscle based on the expression of different *troponins*. Interestingly, we discovered previously unreported gradients of *dysf* and *fln* in the indirect flight muscle, which may indicate regional gradients of transcriptional activity in these very large cells containing more than 1000 nuclei per cell (**fig. S23D**). In visceral muscles, we noted a separation of subtypes based on neuropeptide receptors being expressed. We identified 4 subtypes of visceral muscle in the gut based on expression of the *AstC, Ms, Dh31* and *CCAP* neuropeptide receptors, indicating potential modulators for muscle contraction (Siviter et al., 2000). Receptors for Ms and Dh31 have been described to be expressed in spatially restricted domains in the gut, suggesting that the other two neuropeptides might also exhibit spatial domains (Hadjieconomou et al., 2020; LaJeunesse et al., 2010). Furthermore, the corresponding neuropeptides are released by neuroendocrine cells in similar spatially bound sections (Nässel, 2018). Interestingly, all visceral muscle cells, including the ovarian sheath muscle, are enriched for the receptor of *Pdf*, a neuropeptide involved in circadian rhythms in the brain, but that has also been found to regulate ureter contractions, pointing towards a dual role of circadian rhythm and muscle contraction (Talsma et al., 2012).

### Transcription factors and cell type specificity

Our data allow comparison of gene expression across the entire fly for 251 annotated cell types. Clustering cell types showed the germline cells as the most distinct group, followed by neurons (**fig. S24-29**). Marker genes were calculated for every cell type using the whole FCA data as background, with 14,240 genes found as a marker for at least one cell type and a median of 638 markers per cell type (min: visceral muscle (94), max: spermatocyte (7736)). Interestingly, markers that are specific for cell types in a tissue were not always specific when using the whole body as background, highlighting the importance of choosing the background depending on the goal **(fig. S30)**.

To begin to understand what underlies the specific transcriptomes that we observed per cell type, we analysed the expression of all predicted transcription factors (TFs) (Larkin et al., 2021) encoded by the fly genome, and investigated whether certain TFs are specifically expressed in a single cell type (**Fig. 5A**). To assess this, we used the *tau* score of tissue specificity (Yanai et al., 2005) and identified 500 TFs with a score > 0.85, indicating a high specificity for one or very few cell types (**Supplemental Table S1**). Interestingly, 127 of these TFs are “CG” numbers (computed genes without a symbol), indicating that their function is poorly studied. The cell type-specific expression of these TFs strongly suggests that these uncharacterized genes play functional roles, and these findings may therefore lead to new studies to characterize mutant phenotypes of these CG genes in the linked cell types. Interestingly, we found that the male germline stands out in showing expression of a great number of cell type-specific TFs.

**Figure 5:**
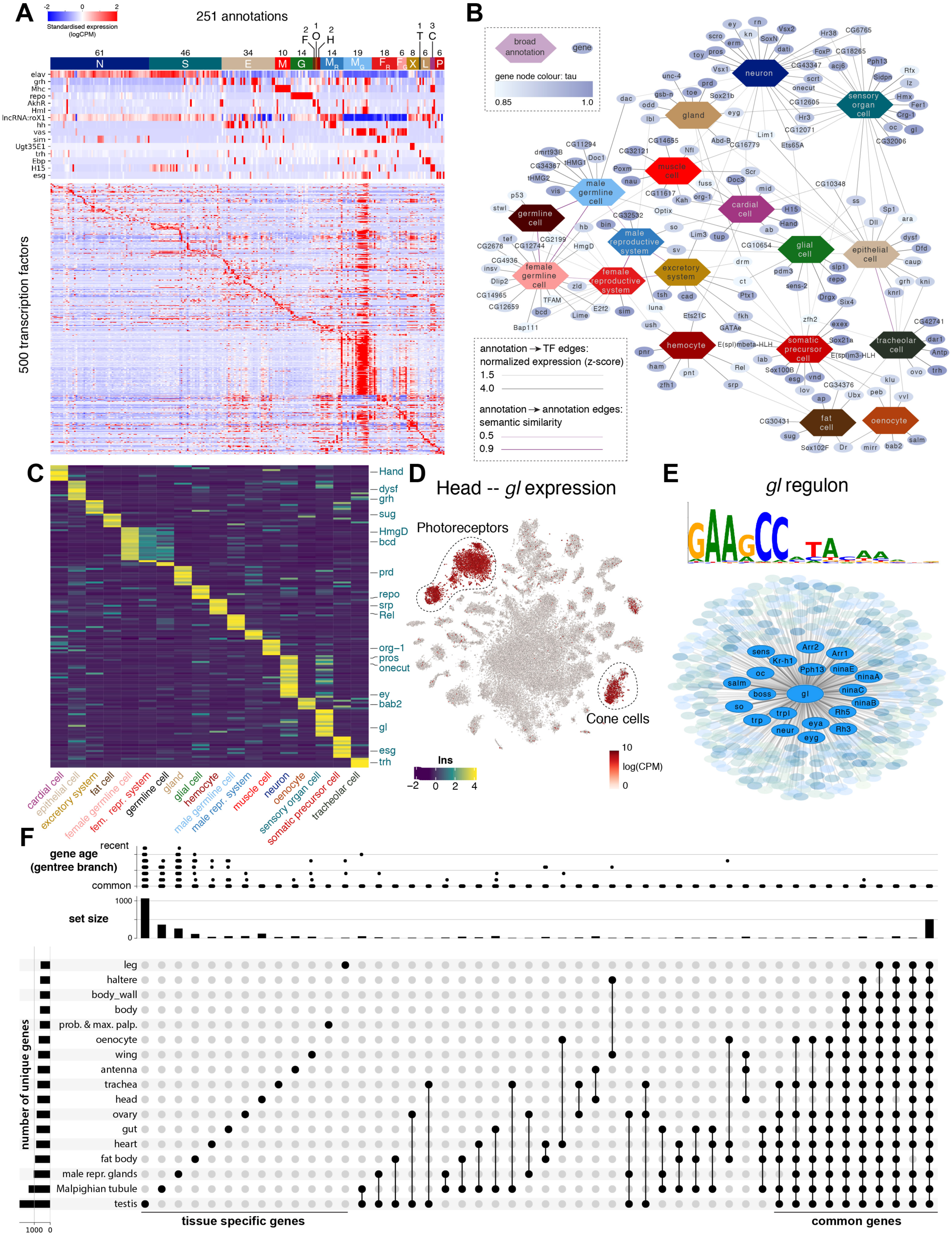
**Transcription factor (TF) pleiotropy versus cell type specificity** **(A)** Heatmap showing the expression of key marker genes and unique TF profiles for each of the annotated cell types. TFs are selected based on tau factor. Cell types are grouped based on hierarchical terms: CNS neurons (N), sensory organ cells (S), epithelial cells (E), muscle cells (M), glia (G), fat cells (F), oenocytes (O), hemocytes (H), (fe)male reproductive system and germline (MR, MG, FR, FG), excretory system (X), tracheal cell (T), gland (L), cardial cell (C), somatic precursor cell (P). **(B)** A network analysis of TFs and cell classes based on similarity of ontology terms, reveals unique and shared TFs across the individual tissues. **(C)** Heatmap showing the expression of unique TFs per categorical term. Factors from the literature are highlighted. **(D)** *Glass* is uniquely expressed in photoreceptors and cone cells in the head. **(E)** Overview of the *Glass* regulon of 444 target genes, highlighting known photoreceptor marker genes. **(F)** Comparison of genes expressed per tissue (mean logCPM>1) shows highly unique gene expression in the testis, Malpighian tubule and male reproductive glands, while also highlighting a common module of conserved, ubiquitously expressed genes. Only sets with more than 10 genes are shown. The left bar graph shows the number of unique genes for each tissue. The top bar graph shows the gene age in branches, ranging from the common ancestor to *Drosophila melanogaster*-specific genes (http://gentree.ioz.ac.cn).

Similar Similar analysis across broad cell types **(Fig 5B, C)** identified 156 TFs with high *tau* scores, for example the previously known regulators *grh* for epithelial cells and *repo* for glial cells; as well as 24 uncharacterised genes. These factors can be visualized in a network, also taking semantic similarity into account for linking similar terms (see Methods). This led to the grouping of CNS neurons and sensory organ cells, including many sensory neurons, with shared pan-neuronal factors such as *onecut* and *scrt* but each cluster having a unique set of TFs, such as *ey*, *scro* and *dati* for CNS neurons and *lz* and *gl* for sensory neurons. Further, we found *esg*, *Sox21a* and *Sox100B* expression was mostly restricted to somatic precursor cells.

In addition to the specificity of TF expression, we predicted gene regulatory networks based on co-expression and motif enrichment using the SCENIC method (Aibar et al., 2017). Because of the stochasticity of this network inference method, we ran SCENIC 100 times, enabling the ranking of predicted target genes by their recurrence. This approach selected 6112 “regulons” for 583 unique TFs across all the tissues, whereby each regulon consists of the TF, its enriched motif, and the set of predicted target genes. In fat cells, our analysis predicted a regulon for *sugarbabe* (*sug*), a sugar-sensitive TF necessary for the induction of lipogenesis (Mattila and Hietakangas, 2017). In photoreceptors, the analysis identified a *glass* (*gl*) regulon, with key photoreceptor markers such as *Arr1*, *eya* and multiple rhodopsins as predicted target genes **(Fig. 5D, E)**. This is in line with previous studies where *gl* was found as the key photoreceptor regulator (Bernardo-Garcia et al., 2016; Moses et al., 1989). The SCENIC predictions for all cell types can be easily investigated via the SCope browser (https://flycellatlas.org/scope).

Comparative analysis of genes across all tissues (**Fig. 5F**) allowed identification of genes expressed tissue-specifically versus commonly expressed genes, such as a shared set of 555 housekeeping genes that are expressed in all tissues. There was a wide spectrum between tissues, with the testis having the highest number of uniquely expressed genes, followed by the Malpighian tubule and male reproductive glands. Interestingly, these tissue-specific genes seemed to be evolutionarily “younger” based on GenTree age compared to the set of commonly expressed genes that are all present in the common ancestor. This suggests that evolutionary pressure works on the tissue specialization level, with the highest pressure on testis, male reproductive tract, and Malgpighian tubules (Shao et al., 2019). In addition, this analysis allowed the estimation of transcriptomic similarity or difference measured by the number of shared unique genes. For example, the two flight appendages, the haltere and wing, share a set of 16 unique genes including *vestigial*, a key factor for wing and haltere development (Williams et al., 1991), presumably reflecting the evolutionary origin of halteres as a modified wing (Lewis, 1978).

In conclusion, TF expression analysis reveals many known regulators, with specific expression across the adult, as well as dozens of previously uncharacterised genes with specific expression that constitute promising candidates for downstream functional characterization.

### Analysis of sex-biased expression and sex-specialized tissues

Much of the genome showed sex-biased expression (Parisi et al., 2004), presumably to support the function of highly sex-specialized gonads and reproductive tract tissues. Sequencing male and female samples separately allowed us to study sex-related differences in tissues common to both sexes. Comparing male-versus female-derived nuclei for all common tissues (**fig. S31**), we found the top male- or female-specific genes were known male or female markers (*roX1/2*, *Yp1/2/3*). Interestingly, a large fraction of genes with male-enriched expression were CGs (Andrews et al., 2000), suggesting that these previously uncharacterized genes may have sex-related functions. The primary sex determination pathway is activated by a two X chromosome karyotype followed by an alternative splicing cascade beginning with *Sex-lethal* (*Sxl*) autoregulation, female-specific splicing of *transformer* (*tra*), and finally sex-specific splicing of *doublesex* (*dsx*) to encode female- or male-specific TFs (Salz and Erickson, 2010) (**Fig. 6A**). Consistent with this, we found *dsx* expression in a largely non-sex-specific pattern, while many other genes showed sex-biased expression (**Fig. 6B**).

**Figure 6.**
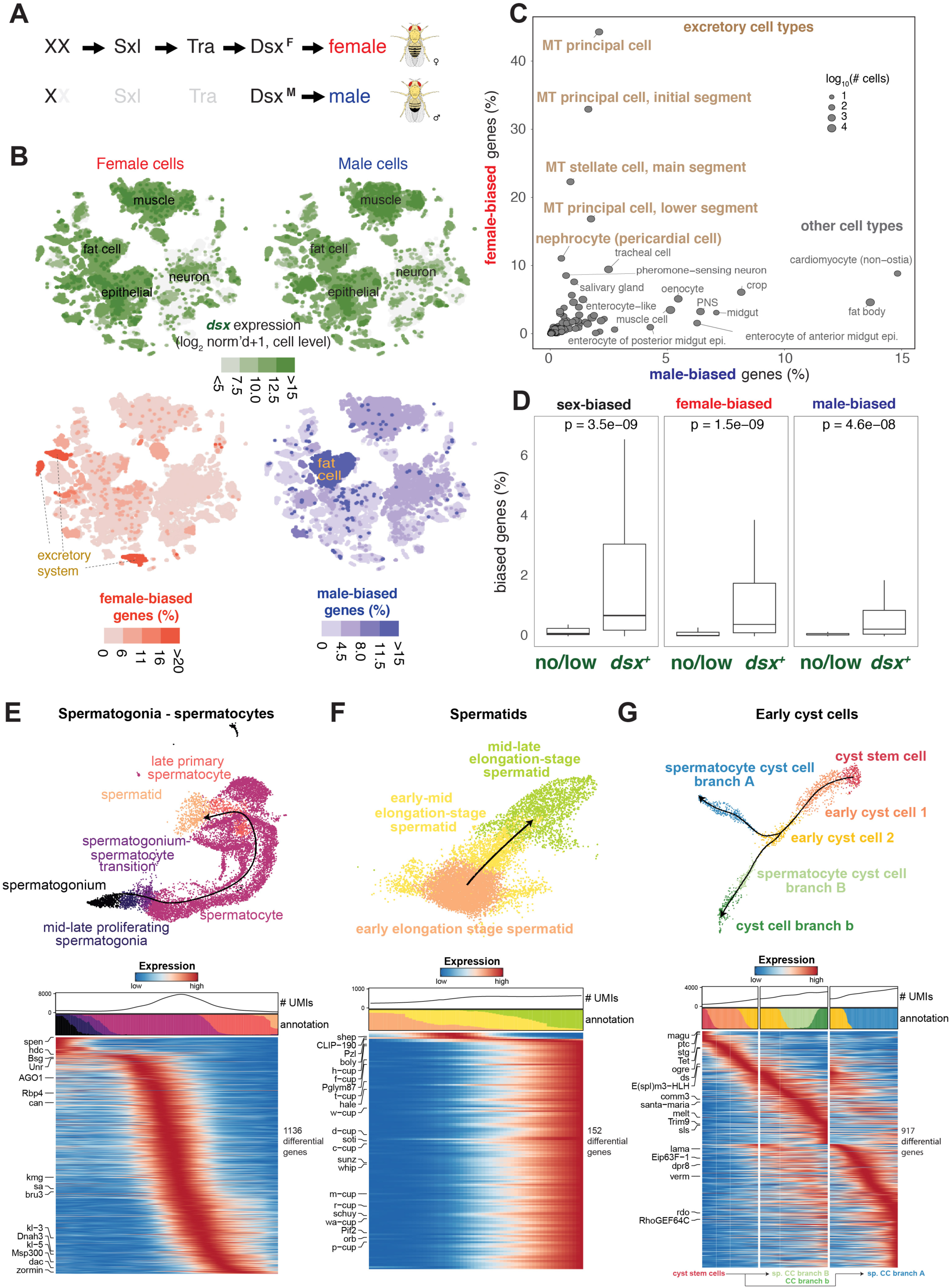
**Sex-biased expression and trajectory analysis of testis cell lineages** **(A)** Simplified sex determination pathway. Sex chromosome karyotype (XX) activates Sex-lethal (Sxl) which regulates transformer (Tra), resulting in a female Dsx isoform (Dsx^F^). In XY (or X0) flies, Sxl and Tra are inactive (light gray) and the male-specific Dsx^M^ is produced. **(B)** Dsx expression and female- and male-biased expression projected on tSNE plots of all female (left column) and male (right column) cells except reproductive tissue cells (see Supplementary Table S3). The top two plots showing *Dsx* expression. The bottom two plots showing female- and male-biased expression, measured as the percentage of genes in the cluster showing biased expression in favor of the respective sex (see Supplementary Table S4). These percentage values were computed for each annotated cluster and those cluster-level values were projected on the individual cells in the corresponding clusters. For all four tSNE plots, values outside the scale in the heatmap key, are represented by the closest extreme color (denoted by > and < signs in the scale). **(C)** Scatter plot of female- and male-bias values across non-reproductive cell clusters defined as % sex-biased genes (at least 2-fold change with FDR < 0.05 on Wilcoxon test and BH correction) in the cluster (see Supplementary Table S4). Data point size shows the number of cells per cluster (key). Selected clusters are labeled, with those from excretory cells highlighted (brown). MT, Malpighian tubule. **(D)** Box plots showing the relationship between *dsx* gene expression and sex-biased expression (see Supplementary Table S4). Clusters (B) were partitioned into the set of clusters with Dsx expression (*dsx*^+^) or not (no/low) using *dsx* expression in germ cells as an expression cut-off. Sex-, female-, and male-biased expression in the two groups were plotted. Each box shows hinges at first and third quartiles and median in the middle. The upper whisker extends from the upper hinge to the largest value no further than 1.5 * IQR from the hinge (where IQR is the inter-quartile range, or distance between the first and third quartiles). The lower whisker extends from the hinge to the smallest value at most 1.5 * IQR of the hinge. Outliers are not shown. Wilcoxon test p values shown. **(E–G)** Trajectory of testis subsets. We used slingshot to infer a possibly branching trajectory for spermatogonia-spermatocytes (E), spermatids (F), and early cyst cells (G). Shown are the trajectories on a UMAP (top) and the expression patterns of the strongest differentially expressed genes, together with the smoothed proportions of annotated cells and average number of unique molecular identifiers (UMIs) along the trajectory (bottom).

As many cell types common to both sexes showed sex-biased expression, we performed differential expression between sexes for all cell types. Notably, cell types tended to show either high female- or male-bias, not both (**Fig. 6B-C**). We found striking female-bias in the excretory system, including the principal and stellate cells of the Malpighian tubule (MT) and in the pericardial nephrocytes (**Fig. 6C**). Examples of genes expressed with female-bias include *eEF1A1*, *eEF2*, *irk3*, and *Rack1*, which are also differentially expressed when Cl^−^ channels are disrupted in a *Drosophila* cystic fibrosis model (Kim et al., 2020). Other female-biased genes (i.e., *Ics* and *whe*) were differentially expressed under high salt conditions (Kim et al., 2017), suggesting sex-bias in nephric ion transport. Across cell types, sex-biased expression strongly correlated with *dsx* expression (**Fig. 6D**) (Arbeitman et al., 2016; Clough et al., 2014), consistent with the role of Dsx as a key regulator.

For analysis of sex-specialized tissues, we focused on the testis (plus seminal vesicle) as a case study, as this dissected sample had one of the most complex cell type compositions, the highest number of ’unique’ marker genes **(Fig. 5F)**, and two dynamic differentiation trajectories. Among all tissues in the adult fly, those best characterized with ongoing cellular differentiation are the gut, ovaries, and testis. Trajectory analysis has been performed on gut and ovary stem cell lineages in previous studies (Hung et al., 2020; Jevitt et al., 2020; Rust et al., 2020), and our FCA data on gut and ovary accurately co-clustered with these published datasets (**fig. S32, S33**).

The testis has two populations of stem cells, namely the somatic cyst stem cells (CySCs) that produce cell types with supporting roles that are essential to spermatogenesis, and the germline stem cells (GSCs) that produce haploid sperm (**Fig. 2C**). Trajectory inference on germ cells from the testis (**Fig. 6E-F**) revealed transitions from GSCs to proliferating spermatogonia, then early spermatocytes (e.g., marker genes include *kmg* and *Rbp4;* **Fig. 2C**), followed by progressively maturing spermatocytes, early spermatids, and finally mid-to-late elongation stage spermatids. As expected, the spermatocyte stage featured a robust increase in the number of genes being transcribed (high UMI) (**Fig. 6E**), leading to a substantial rise in RNA complexity. Many of the genes that are strongly upregulated in spermatocytes (*kmg, Rbp4, fzo, can, sa*, and, for later spermatocytes, Y-linked fertility factors *kl-3* and *kl-5*) are not substantially expressed in most other cell types. A surprise in the FCA data was that late spermatocytes showed expression of many genes that were normally thought of as markers of somatic cells (*Upd1* and *eya*, for example). Notable otherwise cell type-specific examples were marker genes from **Fig. 4B**, *grh* (epithelial cells), *Syt1* (neurons), *Mhc* (muscle), and *Hml* (**Fig. 5A**), although the level of expression of these genes in spermatocytes was considerably lower than in their cognizant marked tissue. What role, if any, these genes may have in the biology of germ cells remains to be investigated. Early spermatids, as expected from their transcriptional quiescence, had a very low number of nuclear transcripts (**Fig. 6F**, low UMI). Later, elongating spermatids turn on a coordinated burst of new transcription as previously identified (Barreau et al., 2008). Our snRNA-seq study identified a number of new such genes (**Fig. 6F**). Again, the comprehensive Tabula *Drosophilae* snRNA-seq effort identified many new genes to be investigated for roles in differentiation of specific cell types.

The early stages of the somatic cyst cell lineage also emerged as a clear trajectory, starting from CySCs expressing the cell cycle marker *string*, transitioning into post-mitotic (no *string* expression) early cyst cells, then branching into two types of cyst cells likely associated with spermatocytes (**Fig. 6G)**. Together, the FCA data from the progression of germ cell types and early somatic cyst cell types identify many additional genes differentially up or down regulated at specific steps in the respective stem cell lineages, presenting a unique opportunity for functional tests to identify new players in the transitions from one cell state to the next.

## Discussion

With the rapid development of single cell sequencing technologies in the past decade, single cell transcriptomic atlases of the whole organism have been achieved for *C. elegans* (Cao et al., 2017) and partially for mouse and human (Cao et al., 2020; Han et al., 2018, 2020; Tabula Muris Consortium, 2020; Tabula Muris Consortium et al., 2018). Here, we provide the first single cell transcriptomic map of the entire adult *Drosophila*, a premier model organism for studies of fundamental and evolutionarily conserved biological mechanisms ranging from development to diseases. The Fly Cell Atlas provides a valuable resource for the *Drosophila* community as a reference for studies of gene function and disease models at single cell resolution. Indeed, combining single cell omics read-outs with the powerful genetic tools available in *Drosophila* to test function *in vivo* holds great potential for new discoveries.

One of the key challenges in large scale atlassing projects is the definition of cell types. Here we solved this by using a consensus-based voting system across multiple resolutions. An FCA cell type is thus defined as a cluster detected at any clustering resolution (Leiden, ranging from 0.4 to 10 (50 in brain)) that could be separated by the expression of known marker genes from other clusters. Further, every annotation was manually curated by a tissue expert, leading to a high confidence dataset with over 250 annotated cell types. Still, we would like to add two notes. First, there are differences in annotation depth with some cell types only linked to broad cell types (e.g. epithelial cell), in contrast to much more detailed cell types (e.g., different types of olfactory receptor neurons, ORNs). For example, in the gut dataset five clusters were identified as crop cells, each with their own transcriptome profile, but without literature knowledge on how to further divide them. These shallow annotations provide information about the cell’s function, and present ample opportunities for follow-up studies to further identify them based on novel marker genes and to highlight new cell types. Secondly, while many marker genes are very useful in identifying cell types, we have also detected cases where marker gene expression was not congruent with cluster expression. This can be caused by discrepancies between mRNA expression and protein, or by mistakes made in literature. These examples highlight the need and the opportunities presented by the FCA to serve as the basis for future validation.

Comparative analysis of different scRNA-seq datasets poses common challenges, including variation in genetic background, sequencing technologies, and sample preparation. To minimize these effects, we used the same wild-type *w^1118^* flies from one laboratory for most tissues except for tissues of which simple dissection is impossible, such as tracheal cells. Second, we used a unified snRNA-seq platform for all samples, because for many adult *Drosophila* cuticular tissues, such as the antenna, wing, leg and haltere, it is extremely difficult to isolate intact cells. In addition, isolation and sequencing of nuclei solves the issue associated with large multinucleated cells (e.g., muscle), and collection of (frozen) tissues from different laboratories. In a recent study (McLaughlin et al., 2021), we directly compared snRNA-seq and scRNA-seq of the same fly cell types and found that about 70–90% of transcriptomic information was preserved from snRNA-seq compared to scRNA-seq. In addition, especially important for dynamic cell types differentiating in adult stem cell lineages, snRNA-seq offers a more immediate picture of changes in transcript expression as cells progress, separable from mRNAs purduring from later stages or stored for later usage.

To further validate the snRNA-seq dataset, we compared single-nucleus FCA data with previously published scRNA-seq datasets from three tissues: brain (**Fig. 3E**), gut (**fig. S32**) and ovary (**fig. S33**). In all three cases, our data not only included all previously identified cell types, but also revealed additional cell types not detected in previous studies. For example, in the ovary, our data captured late-stage germ cells while three published datasets did not. In addition, the Tabula *Drosophilae* data set has increased our knowledge of late-stage follicle cells and allowed identification of new marker genes that were validated using *in vivo* genetic reporters (**fig. S33**). One drawback of snRNA-seq is a higher chance of capturing ambient RNAs, especially for very abundant RNAs that are released from other cells during lysis. For example, a recent study showed that unexpected transcripts from genes encoding odorant binding proteins (Obp) were detected in olfactory receptor neurons using snRNA-seq (McLaughlin et al., 2021).

Sex is a major, and often overlooked, biological variable influencing physiology, behavior, and disease (Arnegard et al., 2020). *Drosophila* has both sex-specific and non-sex-specific cell types. Because we profiled male and female samples separately, our data allowed identification of sex-biased genes expressed at different levels in males vs females in the same cell type. Through detailed analyses, we did not observe significant bias at the cell level for most cell types, meaning cell numbers are similar between males and females for the same cell type, but we did identify a long list of sex-biased genes **(Fig. 6A–D)**. As we are just beginning to fully appreciate the many levels of sex-biased gene expression within the *Drosophila* genome and how these differences meet the developmental and physiological needs of females and males, the sexed single cell data will generate many new testable biological hypotheses.

Our FCA data provide lists of cell type-specific marker genes, in particular TFs, providing a blueprint for generating cell type-specific reporters and expression drivers for studying new aspects of cellular function. However, it is important to realize that some genes that are cell type-specific markers in one tissue might be expressed in more cell types in the whole body. Because marker genes were selected by their higher expression levels in the group of interest compared to the rest of the cell population, when selecting marker genes for such applications, some markers identified in the tissue-specific versus full body analyses may not overlap (see three examples in **fig. S30**). In this regard, the FCA presents the first opportunity to distinguish marker genes on different levels of specificity. In addition, our FCA data together with other atlas data (e.g. *C. elegans*, mouse, human) will allow cross-species studies of common cell types, such as neurons, muscle and blood cells for the generation of evolutionary lineages of cell types in the tree of life.

Finally, our analysis also presents several technical novelties, including the use of Nextflow pipelines improving reproducibility (VSN, https://github.com/vib-singlecell-nf), the availability of raw and processed datasets for users to explore (https://www.flycellatlas.org), and the development of a crowd-annotation platform with voting, comments and references (SCope, https://flycellatlas.org/scope). Furthermore, our atlas is fully linked to existing *Drosophila* databases through the use of a common vocabulary, benefitting its use and integration in future projects. We believe that these elements will serve as inspiration for future atlassing projects and will become indispensable for large-scale single cell experiments. All the FCA data are freely available for further analysis via multiple portals, including SCope, ASAP, and cellxgene, and can be downloaded for custom analysis using other single cell tools (all links can be found on https://www.flycellatlas.org).

## Acknowledgments

We would like to thank the whole fly community for the enthusiastic support for this project. Further, we would like to express our appreciation for William (Bill) Burkholder for organizing the FCA meeting at Biohub, for Cathryn Murphy for helping organize the monthly FCA Zoom meeting, and for Kathleen Vogelaers for coordinating all Jamborees.

## Author contributions

See author list for full contributions.

## Data and materials availability

All data are available for user-friendly querying via https://flycellatlas.org/scope and for additional custom analyses at https://flycellatlas.org/asap. For each tissue, a CellxGene portal is also available (links can be found on www.flycellatlas.org). Raw sequencing data and count matrices can be downloaded from ArrayExpress (accession number E-MTAB-10519 for 10x dataset, and E-MTAB-10628 for smart-seq2 dataset). Files with expression data, clustering, embeddings, and annotation can be downloaded for each tissue, or all data combined, in h5ad and loomX formats from www.flycellatlas.org.

## Materials and Methods

### Fly sample information

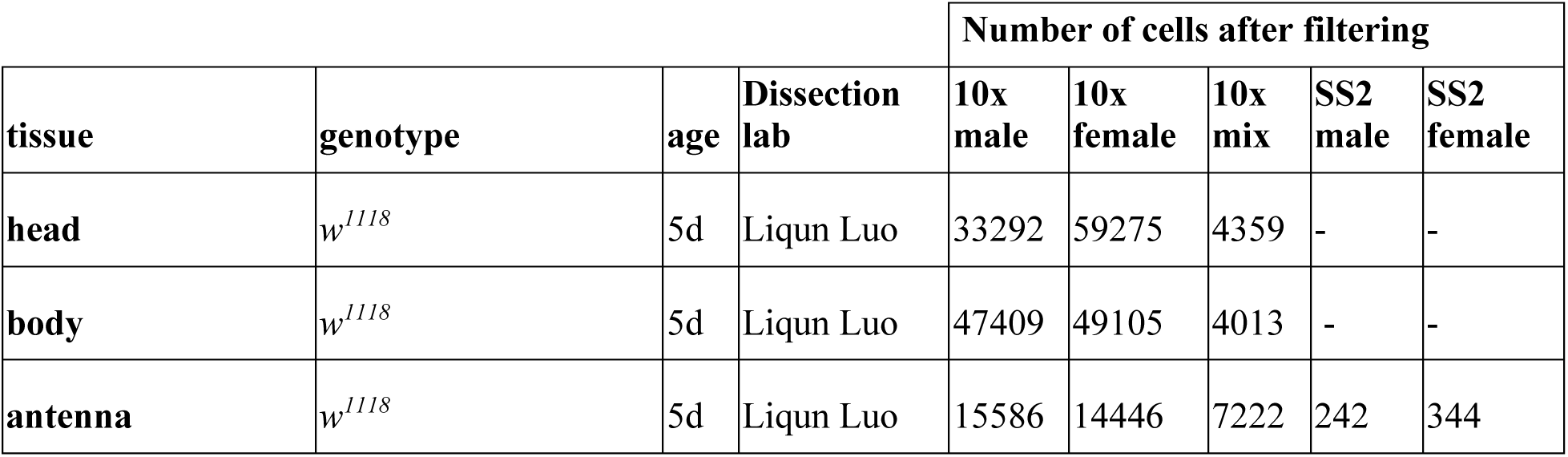
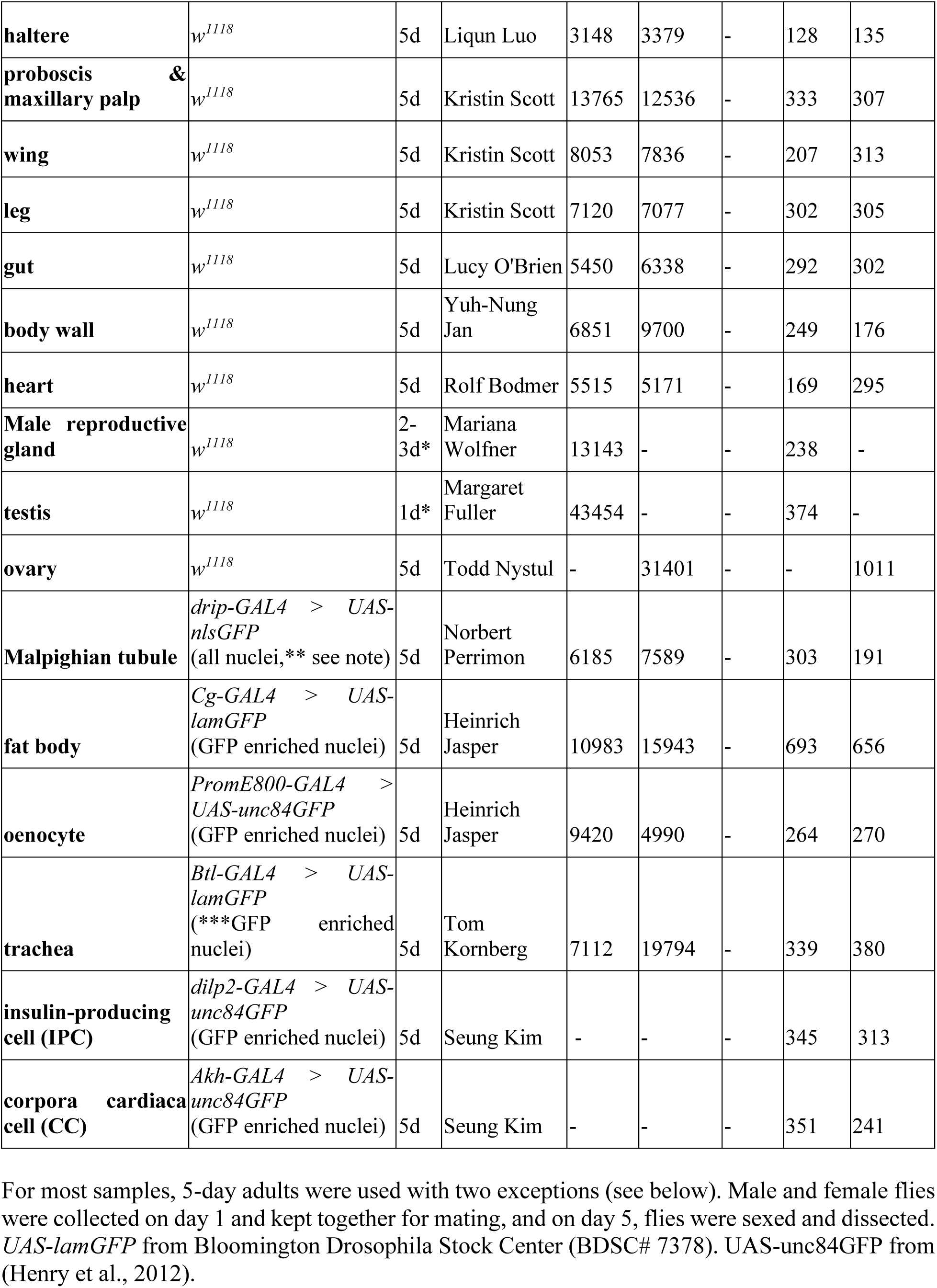
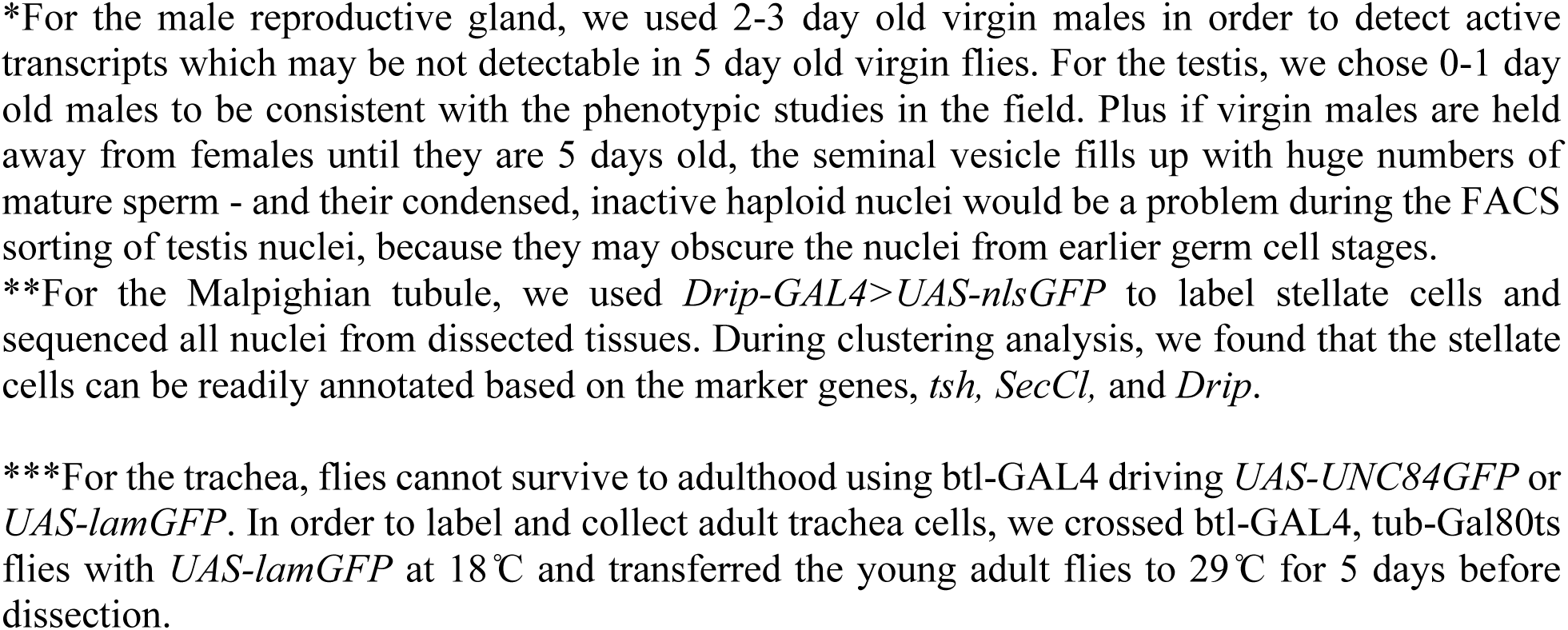

### Single-nucleus RNA-seq

#### Fly dissection and single-nucleus suspension

Fly tissues were dissected by different dissection labs, flash-frozen using liquid nitrogen, stored at – 80 °C, and shipped to Stanford University for processing in the Luo and Quake labs. When using nuclear GFP to label tissues, we compared UAS-nlsGFP, UAS-UNC84GFP and UAS-lamGFP, and found that there was no GFP signal from UAS-nlsGFP after nucleus isolation, while the other two gave good fluorescent signal of the nuclear labeling. Single-nucleus suspensions were prepared as detailed below, largely adapted from our recently published protocol (McLaughlin et al., 2021).

1. Prepare *w^1118^* flies or GAL4 driving UAS-nuclear-GFP flies.
2. Dissect tissue in cold Schneider’s medium, and use P20 pipette (coat the tip with fly body fat) or forceps to transfer them into 100 µl Schneider’s medium in a nuclease-free 1.5ml EP tube on ice. Label the tube clearly using permanent marker. Note: for tissues that float in the medium (e.g., adult antennae), before dissection, prepare three clean dishes: 1st with 100% ethanol, 2nd and 3rd with Schneider’s medium. Rinse the fly in the 1st dish with 100% ethanol for 5 seconds, then rinse the fly in the 2nd dish, and dissect in the 3rd dish.
3. After dissection, spin down samples in 100 µl Schneider’s medium using a bench top spinner.
4. Fresh: The sample can be processed for nuclei extraction immediately following dissection. Frozen: Alternatively, the sample can be flash-frozen for long-term storage. Seal the 1.5 ml EP tube with parafilm and put into liquid nitrogen for >30s. Immediately store the sample at -80°C freezer for long-term storage (several months).
5. Prepare fresh homogenization butter (see details below) and keep on ice.
6. Thaw samples from -80°C on ice if using frozen samples. Spin down samples in 100 µl Schneider’s medium using the bench top spinner, discard medium as much as possible, and add 100 µl Homogenization butter.
7. Optional: if sample pieces are too big, e.g. whole body or whole head, use a pestle motor (Kimble 6HAZ6) to grind the sample for 30s–60s on ice.
8. Add 900 µl homogenization buffer, and transfer 1000 µl homogenized sample into the 1ml dounce (Wheaton 357538). Dounce sets should be autoclaved at 200°C >5h or overnight.
9. Release nuclei by 20 strokes of loose dounce pestle and 40 of tight dounce pestle. Keep on ice. Avoid bubbles.
10. Filter 1000 µl sample through 5 ml cell strainer (35 µm), and then filter sample using 40 µm Flowmi (BelArt, H13680-0040) into 1.5 ml EP tube.
11. Centrifuge for 10 min at 1000g at 4°C. Discard the supernatant. Do not disturb the pellet.
12. Re-suspend the nuclei using the desired amount (we normally use 500-1000 µl) of 1xPBS/0.5%BSA with RNase inhibitor (9.5 ml 1x PBS, 0.5 ml 10% BSA, 50 µl RNasin Plus). Pipet more than 20 times to completely re-suspend the nuclei. Filter sample using 40 µm Flowmi into a new 5 ml FACS tube and keep the tube on ice. Now the single-nucleus suspensions are ready for FACS.

According to our experience, the nuclei are stickier than whole cells. For users making single-nucleus suspension for the first time, we suggest taking 10 µl of the single-nucleus suspension, stain with Hoechst (Invitrogen 33342), and check on a cell counter slide to confirm if they are mostly individual nuclei. If nuclei are not sufficiently dissociated, adjust above steps (e.g., increase the number of strokes of the tight pestle when releasing nuclei).

|  | Amount | Storage | Item (add in this order) | Final concentration |
| --- | --- | --- | --- | --- |
| 1 | 10 ml | RT | H2O (nuclease free) |  |
| 2 | 0.856 g | RT | Sucrose (nuclease free) | 250 mM |
| 3 | 100 µl | 4°C | 1M Tris PH 8.0 | 10 mM |
| 4 | 250 µl | 4°C | 1M KCl | 25 mM |
| 5 | 50 µl | 4°C | 1M MgCl<sub>2</sub> | 5 mM |
| 6 | 100 µl | 4°C | 10% Triton-x 100 | 0.1% |
| 7 | 50 µl | -20°C | RNasin Plus (Promega, N2615) | 0.5% |
| 8 | 200 µl | -20°C aliquots | 50x protease inhibitor (Promega, G6521) | 1x |
| 9 | 50 µl | -20°C | 20mM DTT | 0.1 mM |

#### FACS

We used the SONY SH800 FACS sorter for collecting nuclei. Nuclei were stained by Hoechst-33342 (1:1000; >5min). For wildtype tissues, Hoechst+ nuclei were collected; for collecting GFP+ nuclei, we first gate on Hoechst+ events and then choose GFP+ population.

Since polyploidy is common for many fly tissues, we observed different populations of nuclei according to DNA content (Hoechst signal). For example in the testis, we have observed 6 different populations with different Hoechst signal intensity, and confirmed that the Hoechst signal intensity correlates well with the nuclei size. For the fly gut, we observed 5 different populations with different Hoechst signals. In order to include all cell populations with different nuclear sizes, we have included all nuclear populations during FACS except for testis.

In the testis, haploid spermatids present at 64 for each germ line stem cell. To avoid overrepresentation of small haploid spermatids in the testis sample, we used the following strategy to collect testis nuclei. We have sequenced the testis for three 10x runs. For the first run, we collected nuclei with all different sizes, and for the other two runs, we collected nuclei without the population of the smallest size (haploid spermatids).

During collection, individual nuclei were collected either to 384-well plates for smart-seq2 or to one tube for 10x Genomics. For 10x Genomics, nuclei were collected into a 15ml tube with 500ul 1x PBS with 0.5% BSA as the receiving buffer (RNase inhibitor added). For each 10x Genomics run, 100k–400k nuclei were collected. Nuclei were spinned for 10min at 1000g at 4 °C, and then resuspended using 40ul or desired amount of 1x PBS with 0.5% BSA (RNase inhibitor added). 2ul nucleus suspension was used for counting the nuclei with hemocytometers to calculate the concentration. When loading to the 10x controller, we always target at 10k nuclei for each channel. We observed that loading 1.5 folds more nuclei as recommended by the protocol allowed us to recover about 10k cells after sequencing. For example, if the concentration is 1500 nuclei per ul for one sample, we treat it as 1000 nuclei per ul when loading to the 10x controller.

#### Library preparation and sequencing

Smart-seq2 sequencing libraries were prepared following the protocol we previously described (McLaughlin et al., 2021). Sequencing was performed using the Novaseq 6000 Sequencing system (Illumina) with 100 paired-end reads and 2×12 bp index reads.

10x Genomics sequencing libraries were prepared following the standard protocol from 10x Genomics 3’ v3.1 kit with following settings. All PCR reactions were performed using the Biorad C1000 Touch Thermal cycler with 96-Deep Well Reaction Module. 13 cycles were used for cDNA amplification and 16 cycles were used for sample index PCR. As per 10x protocol, 1:10 dilutions of amplified cDNA and final libraries were evaluated on a bioanalyzer. Each library was diluted to 4 nM, and equal volumes of 18 libraries were pooled for each NovaSeq S4 sequencing run. Pools were sequenced using 100 cycle run kits and the Single Index configuration. Read 1, Index 1 (i7), and Read 2 are 28 bp, 8 bp and 91 bp respectively. A PhiX control library was spiked in at 0.2 to 1% concentration. Libraries were sequenced on the NovaSeq 6000 Sequencing System (Illumina).

### Sequencing read alignment

Prior to read alignment, the raw FASTQ files were processed with the index-hopping-filter software from 10x Genomics (version 1.0.1) in order to remove index-hopped reads. More information about this software is available at https://support.10xgenomics.com/docs/index-hopping-filter.

A Cell Ranger (version 3.1.0) index was built from a pre-mRNA GTF which was derived from the Flybase version r6.31 GTF. A complete recipe on how to build this custom pre-mRNA GTF is available here: https://github.com/FlyCellAtlas/genome_references/tree/master/flybase/r6.31

For 10x, the filtered reads (index-hopped reads removed) were processed all the way up to the gene expression matrix using Cell Ranger (version 3.1.0). For Smart-seq2, reads were aligned to the *Drosophila* melanogaster genome (r6.31), the same as for 10x read alignment, using STAR (2.5.4). Gene counts were produced using HTseq (0.11.2) with default settings except ‘-m intersection-strict’. Gene counts were generated using the same GTF file as for 10x, covering both exonic and intronic regions. Low-quality nuclei having fewer than 10,000 uniquely mapped reads were removed.

### Cell filtering and clustering: *Relaxed* version

To verify the accuracy and robustness of the data processing steps, we examined marker gene expression and compared multiple methods and parameter settings for the various preprocessing steps, including index hopping filtering, genome annotation and counting intronic reads, doublet detection (scrublet), read decontamination (DecontX), batch effect removal (harmony), dimensionality reduction (automated PC determination), and clustering (Leiden). At each step QC plots are generated (Suppl. Data 1) and the final loom files can be visualised in our SCope and ASAP analysis and visualization platforms, or be downloaded for custom analysis.

Two versions of the processed data were generated: a *Relaxed* and a *Stringent* version. Here we focus on describing the *Relaxed* version of the data. To know more about the difference between the two versions please read the next section.

The Scrublet software was chosen for doublet removal. Doublet scores were calculated from the raw expression matrix generated by Cell Ranger and using Scrublet (version 0.2.1, Docker image: vibsinglecellnf/scrublet:0.1.4). The strategy taken here to remove the doublets from each sample relies on the multiplet rate one can expect from running a Single Cell 3’ 10x Genomics experiment. This number depends on the number of recovered cells. In order to estimate this rate as a function of the number of recovered cells, a linear regression was performed on the multiplet rate table (see Chromium Single Cell 3’ Reagent Kits v2 User Guide • Rev F) in order to determine the slope and the bias terms. Those numbers were 0.008 and 0.0527 respectively. Given this model, for each sample the top N cells, ranked by the doublet score, were considered as doublet hence removed. The data was further processed using the Python package Scanpy (version 1.4.4.post1, through Docker image: vibsinglecellnf/scanpy:0.5.0).

Two additional filters, cellwise and genewise, were applied. The cell filter is based on hard thresholds applied on some of the quality metrics (QC). All cells expressing less than 200 genes were filtered out. Moreover, cells exceeding a 15% mitochondrial content were removed. Regarding the gene filter, all genes not expressed in at least 3 cells were filtered out.

For the different analysis runs with VSN-Pipelines, the samples were concatenated using the anndata.concatenate (join=outer). Consequently, the combined matrix was normalized (scanpy.pp.normalize_per_cell, with counts_per_cell_after=10000) and log transformed (scanpy.pp.log1p). Highly variable genes were selected using sc.pp.highly_variable_genes (min_mean=0.0125, max_mean=3, min_disp=0.5). The data was further scaled so that each gene had unit variance and values exceeding a standard deviance of 10 were clipped. In order to determine the number of principal components (PC) to select, a cross-validation approach was performed using the pcacv module (available in the VSN-Pipelines). The scaled matrix was projected to a principal component analysis (PCA) space using the scanpy.tl.pca function (svd_solver=arpack). Batch correction was applied using the Harmony software (Docker image: vibsinglecellnf/pcacv:0.2.0) with default parameters and using sample as batch variable. The neighborhood graph was calculated from the corrected PCA space with scanpy.pp.neighbors and default parameters except for the number of PCs (see aforementioned). For visualization purposes, two non-linear dimensionality reduction methods were used: t-SNE and UMAP. The t-SNE embeddings were generated using scanpy.tl.tsne while the UMAP embeddings with sc.tl.umap. Default parameters were used for both methods except for the number of PCs (see aforementioned). Clustering was performed using the Leiden algorithm via scanpy.tl.leiden (default parameters) except for the resolution parameter where a range of values were selected. A default clustering was selected using a custom method available in the directs module from VSN-Pipelines (vibsinglecellnf/directs:0.1.0). The default clustering is selected as follows: for a range of min_cluster_size and min_samples, a density-based clustering is performed using HDBSCAN on the t-SNE embedding; an adjusted rand score is computed between this clustering and the previously generated Leiden clusterings; a clustering is assigned to each pair of parameters which maximizes the score; the final selected clustering is the one that maximizes the most the score over all pairs. Cluster markers for each of the generated clusterings were computed from the log_2_ normalized expression matrix by means of scanpy.tl.rank_genes_groups (method=wilcoxon, n_genes=0). To ensure reproducibility of the 10x Genomics data processing, all the analyses from raw counts to final processed files (.loom and .h5ad) were performed using the VSN-Pipelines (https://githhub.com/vib-singlecell-nf/vsn-pipelines).

### Cell filtering, decontamination and clustering: *Stringent* version

In the previous section, we described how the *Relaxed* version of the data was generated. The main reason to generate a *Stringent* version was that we identified a significant number of cells, we called “black hole” cells, which are expressing multiple big cell type markers e.g.: *grh* (epithelial cell), *Mhc* (muscle cell), *onecut* (neuron). These cells likely originated from droplets that were contaminated by ambient RNA.

DecontX was chosen in order to correct for this bias. Practically, the raw counts generated from the Cell Ranger pipeline were corrected using this algorithm, available in the celda R package (version 1.4.5, Docker image: vibsinglecellnf/celda:1.4.5). The corrected counts were then rounded using the R base::round function and newly generated empty cells were removed.

Additionally, we applied a more stringent filter on the cells. This cell filter is based on the median absolute deviation (MAD) from the median of the following quality control (QC) metrics across all cells: n_counts i.e.: number of counts per cell, n_genes i.e. number of genes per cell. A value is considered an outlier if it is greater than 3 MADs away (both directions) from the median of these two metrics. This filter strategy is applied in the log_2_ space of these QC metrics. Moreover, minima of 200 genes and 500 counts per cell are required for a cell to be considered in the downstream analysis. All cells exceeding a 5% mitochondrial content were removed.

Finally, after the highly variable gene selection, the number of UMIs per cell and the percentage of mitochondrial genes were regressed out using scanpy.pp.regress_out. All other steps described previously remained the same.

For most analyses, we have focused on the *Stringent* dataset. The UMAP for testis presented in Figure 2C is plotted using Relaxed dataset because the important hub cell cluster was filtered out by the algorithms, likely due to the expression of many “somatic” transcripts including *Upd1* in late spermatocytes.

### SCope features and applications

SCope platform crowd annotation system was used to gather all annotations that were added during the Jamborees by all tissue experts. All tissue analysis results across all clustering resolutions were used as a basement to annotate the cells of the atlas. The system allows for tracking the author of the annotations as well as their confidence through a like/dislike feature. The latter feature was crucial for building a consensus annotation. Annotated Loom files can be directly downloaded from SCope.

### ASAP features and applications

The ASAP platform (David et al., 2020) was used to perform more detailed analyses on the datasets. In particular, ASAP was used to perform sub-clustering and additional differential expression / marker gene discovery. Since the platform also allows annotating (e.g., a color gradient) cells according to the expression of a particular gene set (rather than a single gene), ASAP was also used to study the activity state of KEGG pathways. Finally, because ASAP allows users to share a project (or its copy) privately with a group, all FCA-related projects have been made public so that researchers can share/clone them freely, and annotated Loom files can then be directly downloaded in ASAP.

### Brain-Head integration (Fig. 3E)

Single cell RNA-seq dataset from Davie et al. was downloaded from GEO (GSE107451) and processed using VSN. Next, the data was integrated with the single nucleus data from the FCA using Seurat. Data was normalized using SCT normalization (Hafemeister and Satija, 2019) and batch correction was performed as described (Stuart et al., 2019). 150 components were selected for clustering and UMAP/tSNE visualization. Annotations were added using three computational approaches. First, we transferred annotations based on co-clustering of annotated cells with new cells: if a cluster contained at least 25% of cells with the same annotation, this annotation was transferred to all cells in the cluster. Next, we used a classifier from Özel et al. (Özel et al., 2021) to annotate optic lobe cell types. Finally, we trained an SVM classifier on the Davie et al. data, using scikit-learn in Python, following 10 fold cross-validation, optimizing C, kernel and gamma parameters. All computed annotations were then manually curated in jamborees.

### Trajectory inference of testis subsets (Fig. 6E–G)

We used slingshot to infer a possible branching trajectory in subsets of the testis cells. Specifically, 1) for the spermatogonia-spermatocyte trajectory we used clusters annotated as spermatogonia or spermatocytes, 2) for spermatids we used clusters annotated as early/late spermatids, and 3) for early cyst cells, we used cyst stem cells, early cyst cells and the two spermatocyte cyst cell branches. As input for slingshot, we used Seurat’s FindClusters function with resolution 0.4 to find clusters, and the first 20 PCA components as dimensionality reduction. We also provided the start cell of the trajectory as “cyst stem cell”, “spermatogonium”, and “early elongation stage spermatid”. To determine differentially expressed genes, we used the “calculate_overall_feature_importance” function from dyno (https://github.com/dynverse/dyno) and filtered genes based on a feature importance of at least 0.1 and a log2-fold change along any point in the trajectory of at least 0.5. To map the trajectory onto the UMAP embedding, we used the project_trajectory function, also implemented in dyno.

### Integration of 10x Genomics and Smart-Seq2 data

We integrated 10x Genomics and Smart-Seq2 data using Harmony (Korsunsky et al., 2019). To facilitate and improve integration, we first selected most relevant genes by performing differential gene expression on annotated 10x Genomics data (t-test; Benjamini-Hochberg corrected p-value < 0.1). To integrate individual tissues, we used genes differentially expressed between cell types of a given tissue. To integrate entire cell atlases, we used genes differentially expressed between tissues. After selecting differentially expressed genes, we batch-corrected datasets using Harmony to remove the influence of the sequencing technology. We included an additional step of batch-correction for those tissues in which gender specific clusters were present. In total, we performed additional batch-correction based on the gender for 12/15 tissues.

We next systematically validated integration in the following way. If the marker genes were known, we visually inspected whether marker genes are expressed at the same tSNE location for 10x Genomics and Smart-Seq2 data after integration. In case marker genes were not known, we found differentially expressed genes based on annotations of 10x Genomics data and selected 3-5 genes that are as cluster specific as possible. Finally, for each tissue and cell type we came up with the list of genes and validated whether these genes are expressed at the same location in the tSNE space for 10x Genomics and Smart-Seq2 data. Besides Harmony, we considered three other integration approaches [BBKNN (Polański et al., 2020), Scanorama (Hie et al., 2019) and DESC (Li et al., 2020)] and finally decided to use Harmony based on our validation procedure.

### Annotating Smart-Seq2 data

After integrating 10x Genomics and Smart-Seq2 data, we next aimed at transferring cell type annotations from 10x Genomics to unannotated Smart-Seq2 data. To develop an approach and quantitatively compare performance of different classification methods, we used Smart-Seq2 cells from olfactory receptor neurons (ORN) (McLaughlin et al., 2021) annotated using MARS (Brbić et al., 2020) and manually validated based on the marker-gene expressions. We integrated this dataset with ORN antenna 10x Genomics dataset that was annotated on the same granularity level. The classification accuracy was high (0.88) with a linear logistic regression model and did not improve by using non-linear models. Therefore, we decided to use logistic regression as the base classifier due to its simplicity and interpretability. Finally, for each tissue we used a 10x Genomics dataset as the train set and trained a logistic regression classifier to distinguish different cell types of annotated 10x Genomics data. We then applied the classifier on the Smart-Seq2 dataset to obtain cell type annotations. To confirm that Smart-Seq2 annotations are indeed correct, we checked expressions of known marker genes and validated if they agree with the predicted Smart-Seq2 annotations.

### Comparison of 10x Genomics and Smart-Seq2 data

To compare the number of detected genes between 10x Genomics and Smart-Seq2 data (Fig. 1E), we considered a gene detected if a single read maps to it. In particular, for 10x Genomics data we used UMI greater or equal to 1 as a threshold, while for Smart-Seq2 data we used CPM greater or equal to 1 as a threshold. Given these thresholds, we counted the number of genes detected in at least 1% of cells. To obtain examples of genes that are detected using Smart-Seq2, but not using 10x Genomics (fig. S17G) we obtained a list of genes that are expressed in less than 20 cells in 10x Genomics data, and two to four times more cells in Smart-Seq2 data depending on a tissue.

### Transcription factors and cell type specificity analysis (Fig. 5A-D)

Cell’s expression profiles were averaged in log_2_CPM space and subsequently Z-normalised per gene. We then calculated the tau value for each TF (Flybase r6.36 TF list) using the tspex Python package (version 0.6.2) and 500 TFs were selected and plotted. Regarding the analysis showing the TF specificity heatmap (Fig 5C), we used the log_2_ normalized gene by cell expression matrix. The cells’ expressions were averaged by the broad annotations. We then calculated the tau value for each TF (Flybase r6.36 TF list) using the tspex Python package (version 0.6.2). The values shown in the heatmap are the feature-scaled values using the zscore function available in the package. Only genes passing the thresholds of tau greater than 0.85, log_2_ normalized expression greater than or equal to 0.5 and log_2_ normalized scaled expression greater than or equal to 1 were retained. The plotting was performed using the ComplexHeatmap R package.

The network shown in Fig 5B is based on 5 different .sif files (network files). The first one is the network based on TFs and broad annotations where links passing the aforementioned thresholds were kept. The second one is the network based on TFs and narrow annotations where 0.85, 2 and 5 thresholds were applied respectively. The other networks represent the narrow-to-narrow, broad-to-narrow and broad-to-broad annotation associations. Since the annotations were mainly driven by the EBI OLS system, most of them are associated with a curated FBbt term. We leverage this graph-based ontology structure in order to compute a semantic similarity between annotations using the ontologySimilarity R package (version 2.5). For broad-to-narrow and broad-to-broad annotation associations, the ones with a semantic similarity below 0.4 were removed except for a few of the broad-to-narrow associations that resulted in a loss of broad terms (fat cell to muscle cell, cardial cell to multidendritic neuron and neuron to multidendritic neuron). For narrow-to-narrow connections, after the expression-based filter, only terms that had a TF assigned were kept and moreover we selected for each term the two most connected terms.

Those 5 processed networks were used as input in the Cytoscape software (version 3.8.0) to build the visual network depicted in Fig 5B. The width of the edges represents the log normalized scaled expression (z-score) while the tau values are represented by the colour intensity of the gene nodes.

### Sex-differences analysis

Tissue average expression profiles were calculated in log_2_CPM space. Next, all genes with log_2_CPM>1 were selected as being highly expressed in the tissue. For every gene the evolutionary age was determined using the GenTree database (http://gentree.ioz.ac.cn/). Finally, sets of genes with their evolutionary age were then plotted in an upset plot using Python (UpSetPlot version 0.4.4).

### Sex-differences analysis (Fig. 6A-6D)

For the sex bias analysis, we CPM normalised the gene expression matrix. Next, we filtered out the cells from sex specific samples, i.e., testis, ovary and male reproductive glands and also the cells which were marked to be of ‘mix’ed sex or marked ‘artefact’ or ‘unannotated’ in the annotation. Since ‘body’ samples also contain sex specific tissues, we further removed the cells annotated as germ cells or cells assigned to other sex specific clusters.

To be as stringent as possible, we further removed cells that were either (i) co-clustered in the t-SNE with these sex-specific cell types or organs, or (ii) might be improperly annotated as evidenced by co-expression of mutually exclusive cell-type specific markers. These cells were identified using the SCope web interface and “lasso” tool and removed. The list of cells removed by this manual procedure is included as in **Supplementary Table S2**.

At the end of the filtering, 270,486 cells from 176 annotated clusters remained and were used for our analysis. **Supplementary Table S3** gives the details of these cells and annotations. These were grouped by annotation and for each gene in each annotated cluster, we computed i) it’s sex bias B (B = log2((male_avg+1e-9) / (female_avg+1e-9))) where male_avg and female_avg denote the average expression (computed from the normalized expression matrix) of the male and female cells, respectively, in the cluster and p-value for the bias (multiple tests corrected) using Wilcoxon test (scanpy function sc.tl.rank_genes_groups() with default parameters) of the difference in male and female means, and ii) average dsx expression (normalized).

For each annotated cluster and gene, we denoted the gene to be male-biased in this cluster, if sex-biased B was > 1 (i.e., 2-fold change in favor of male) with FDR < 0.05. Similarly, female-biased if sex-biased B was < -1 (i.e., 2-fold change in favor of female) with FDR < 0.05. A gene was considered sex biased if it was either male or female biased. Using this definition, we obtained the list of 9179 genes which are sex-biased in at least one annotated cluster. Next, for each cluster we computed what percentage of these sex biased genes were male (respectively female) biased in the given cluster. We define these fractions as male-bias (respectively female-bias) of the cluster. This information is kept in the data file **Supplementary Table S4**.

Data used for the SNE visualizations on panel B is kept in the data file **Supplementary Table S3**. This table also includes the dsx expression for each cell, extracted from the normalized expression matrix. The *dsx* expression level displayed on panel B uses log scale: (log2(dsx+1)). The cells with zero *dsx* expression are shown in gray and remaining using the color scale shown in the legend. For the bottom two subpanels each projected cell is colored according to the sex bias (as defined above) of the cluster it belongs to. For comparison with the top panels, we show the female-bias for female cells only on the left and male-bias for the male cells only on the right. For all four subpanels, if the displayed value is outside the scale, we use the closest extreme color (using sign < or >).

For the cut-off for *dsx* presence in panel D, we used 0.1 which is equal to maximum of average dsx expression (rounded to single decimal digit) of all germ cells which are known to not express dsx but show trace expression in FCA data (we note that these clusters are otherwise removed from this analysis and only used to decide the threshold for other clusters in this analysis).

### Common cell type analysis – Hemocyte

With the cross-tissue analysis, we extracted 8,391 Hml^+^ cells from most body parts, including the fat body, heart, body wall, oenocytes, legs, the Malpighian tubule, tissues in the head, and reproductive organs. Harmony was used to remove batch effects with different parameters to control against overcorrection. We tested lambda (ridge regression penalty parameter) and theta (diversity clustering penalty parameter) in a grid with each parameter ranging from 0 to 3. In the end, the T1L2 combination was found to preserve cell types while not overtly separating cells in batches (based on visual exploration). To explore hemocytes in adults, we annotated cell clusters according to the expression of previously published markers in larval hemocytes (Tattikota et al., 2020; Cattenoz et al., 2020; Cho et al., 2020; Girard et al., 2021) and identified twenty-one clusters of Hml^+^ cells (Figure 4D). Crystal cells are readily segregated by high PPO1, PPO2, and lozenge expressions, while plasmatocytes are largely combined as a population most akin to embryonic and larval hemocytes. Plasmatocytes are categorized into five clusters based on their gene expressions and we named the clusters with representative marker genes: Pxn^High^, Nplp2/Tep4^High^, Cecropin^High^, LysX/trol/Pvf2^High^, and nAChRalpha3^High^. LysX/trol/Pvf2^High^ plasmatocytes exhibit lower Hml compared to other plasmatocytes whereas Pxn^High^ shows the highest Hml with phagocytosis markers including crq, Sr-CI, and NimC1. Nplp2/Tep4^High^ plasmatocytes show a prohemocyte marker, Tep4, and an intermediate prohemocyte marker, Nplp2, along with phagocytosis markers. Cecropin^High^ plasmatocytes display immune- or stress responsive genes such as upd3, Mmp1, Mmp2, and puc. Further, we observed Antp and collier expressing Hml^+^ cells reminiscent of the posterior signaling center in the lymph gland (Mandal et al., 2007; Krzemien et al., 2007). Yet, lamellocytes are not observed in adults as previously suggested (Bosch et al., 2019) (Figure 4D). In addition to Hml^+^ cells with classical hemocyte gene expressions, we noticed Hml^+^ cells originating from a single tissue, including the testis and antenna, constitute independent clusters significantly enriched with resident tissue marker genes. Overall, single-cell transcriptome profiles of adult hemocytes provide ample resources for understanding adult immunity, hematopoiesis and repertoires of tissue-resident hemocytes.

Hemocytes in adults are largely resident and the majority is found in the thorax or head while a small fraction circulates the hemolymph (Sanchez Bosch et al., 2019). Thus, cross-tissue dissection of adult hemocytes categorized represent hemocytes in adults. Although *Hml* is a well-known marker for plasmatocytes in embryonic- and larval stages, the expression of *Hml* is heterogenous during development which could hinder labeling the entire population of adult hemocytes.

### Common cell type analysis - Muscle

Muscle cell clusters were identified by their expression of common sarcomeric gene products, including *Mhc, sls, bt* and *Unc-89* (Sarov et al., 2016). With the cross-tissue analysis, we extracted 63,441 muscle cells from most body parts. Harmony was used to remove batch effects with different parameters to control against overcorrection. We tested lambda (ridge regression penalty parameter) and theta (diversity clustering penalty parameter) in a grid with each parameter ranging from 0 to 3. In the end, the T1L2 combination was found to preserve cell types while not overtly separating cells in batches (based on visual exploration). The abundant indirect flight muscle nuclei cluster was uniquely identified by expression of flight muscle-specific markers *TpnC4, Act88F* and *fln* (Schönbauer et al., 2011). Furthermore, the here identified specific expression of different troponinC gene isoforms (*TpnC4, TpnC73F, TpnC41C, TpnC47D, TpnC25D*) was used to further annotate the different muscle clusters taking into account their body part of origin (Chechenova et al., 2017).

### Metabolic clustering using ASAP

For probing the FCA data for an enrichment in Fatty acid synthesis (FAS) and Fatty acid degradation (FAD) metabolic genes, we first downloaded the latest KEGG assignments of genes of *Drosophila* melanogaster (DM) which were mapped to KEGG pathways (FAS: map00061, and FAD: map00071) on December 2020. We next used the ASAP (Automated Single-cell Analysis Pipeline) platform to probe the FCA datasets for a transcriptional enrichment of the genes of these metabolic pathways (David et al., 2020). ASAP generates a score from the difference of expression of the genes in the FAS or FAD gene sets as compared to background genes. In short, for each gene in the gene set, the function will take n random genes from the same expression quantile and add them to a background gene set. The gene set score is calculated as the difference of average expression of the genes in the module score and the genes in the background, for each cell. Scores close to zero indicate a similar expression, positive scores indicate higher expression and negative scores indicate lower expression of the genes in the gene set than the background genes. This function was adapted from the AddModuleScore function from the Seurat package (Satija et al., 2015), and was entirely recorded in Java. We have used the non-normalized parsed Fat body v2 data as input matrix, 24 bins, 100 background genes and set the seed to 42. We plotted the results using the visualization feature implemented in ASAP and colored the cells according to the score values (David et al., 2020). Finally, we used the FCA 0.4 resolution clustering information available on Scope to delineate the frontiers between the different cell clusters.

### Transcription factor pleiotropy analysis

Markers were calculated with the wilcoxon test, comparing every cell type against all other cells. Next only genes with pval adj<0.05 and average log_2_[foldchange] > 1 were selected as selective markers.

### Ovary data integration (fig. S33)

Four scRNA-seq datasets of the adult ovary were used for the data integration: current FCA data and three other published datasets (Rust et al., 2020)(Jevitt et al., 2020; Rust et al., 2020; Slaidina et al., 2020) merged and batch corrected using Seurat v4.0.1. Datasets were processed with Seurat v4.0.1 in RStudio Version 1.4.1103. Batch correction was performed as described (Stuart et al., 2019) 4. Clustering of unannotated cell types was performed using the FindClusters command in Seurat v4.0.1 with a resolution factor of 0.6. Image processing was performed with FIJI. Fly lines were ordered from BDSC: *sick-Gal4* (#76195), *Wnt4-Gal4* (#67449), *UAS-RedStinger, UAS-Flp, Ubi- (FRT.STOP.FRT) -Stinger* (#28281).

## Supplementary Text

### Testis snRNA-seq features developmental trajectories in stem cell lineages

For the following discussion, we are referring to the UMAP plot using the *Relaxed* dataset of the testis in Figure 2C. The 10x snRNA-seq data from adult testis plus seminal vesicle, like other dissected tissues, showed clusters representing several terminally differentiated cell types, including muscle and pigment cells from the testis sheath and a large number of nuclei representing cells in various epithelial components of the organ, such as the terminal epithelial cells and embedded late head cyst cells, which hold the head ends of elongated spermatid bundles at the base of the testis (Fig. 2C). In addition, sample dissection carried over small numbers of tracheal and fat body cells, hemocytes, neurons, and male reproductive tract secretory cells not considered integral parts of the testis (Fig. 2C).

Unlike many other tissues, but similar to the ovary and gut, the geography of the UMAP plot of 10x snRNA-seq data from testis was dominated by continually renewing differentiating cell lineages produced by dedicated stem cells. In testis, there are two such adult stem cell-founded lineages: somatic cyst stem cells (CySCs) produce post-mitotic differentiating daughters that play signaling and key cell-biological supporting roles essential to spermatogenesis, while germline stem cells (GSCs) produce proliferating daughters that undergo a well-orchestrated series of differentiation steps to produce haploid sperms. These two stem cell types are anchored in proximity at the apical tip of the testis by physical attachment to a small cluster of epithelial-like somatic cells termed the hub, which provides close-range niche signals important for the maintenance of the two stem cell states. The interleaved arrangement of GSCs and CySCs and their attachment to the hub ensures that immediate stem cell daughters are positioned to interact: two postmitotic early cyst cells enclose the gonialblast produced by a germ line stem cell to form a cyst, the functional unit of further co-differentiation of both the germ line and the somatic cyst cells. Despite their physical proximity and cooperation in the organ, the germ line and cyst cell lineages map to largely non-overlapping formations in the UMAP plot of 10x snRNA-seq data, consistent with their different embryological origin, cell biology, and known roles.

The dynamic sequence of cell states in the germ line and cyst cell lineages is manifest in the geographical arrangement of nuclei with similar but progressing gene expression patterns in the UMAP plot of the 10X snRNA-seq data from testis (Fig. 2C). Clustering of similar nuclei at a higher resolution (Leiden 6; see Methods) revealed as an emergent property many of the steps of differentiation in the germ line and cyst cell lineages known from the literature. This emergent geography correlates well with the output of trajectory inference on the 10x snRNA-seq data for the germ line and early stages in the cyst cell lineage (Fig. 6E-G). At the view angle presented in Figure 2C, the UMAP plot whimsically brings to mind a hammerhead shark (germ cell lineage) playing a saxophone (somatic cyst cell lineage) watched over by a mermaid (largely epithelial based structural elements such as the seminal vesicle (mermaid head and possibly torso) and the head cyst cells that become embedded in the terminal epithelium at the base of the testis (likely the “calf” of the mermaid lower leg).

The germ line lineage begins at the tail end of “the shark” with the germ line stem cells (GSCs) and proliferating spermatogonia (Fig. 2C and 6E, areas labeled as *5 and *6) at the tapered point of a long, thick tube moving from left to right along the bottom of the UMAP image. After a slight bulge, which may contain germ cells transitioning from spermatogonial proliferation to premeiotic S phase, early spermatocyte markers such as *kmg* and *Rbp4* begin to be expressed (Fig. 2C, areas sp.0-2). Subsequently, clusters of progressively maturing spermatocytes march rightward, with the most mature spermatocyte stages curving upward on the right (Fig. 2C, sp.3-sp.7a). At the top of the tube (head of the shark) lie a mass of early spermatids, with mid-to-late elongation stage spermatids in a blunt projection emerging toward the upper right (Fig. 2C, *4).

Both already known and surprising new features of the germ cell transcription program are visible in the snRNA-seq data. As expected, the spermatocyte stage features a robust increase in the number of different genes being transcribed (high UMI) (Fig. 6E), leading to a substantial increase in RNA complexity. Many of the genes strongly upregulated in spermatocytes (*kmg, Rbp4, fzo, can, sa,* and, for later spermatocytes, Y-linked fertility factors *kl-3* and *kl-5* for example (see SCOPE)) are not substantially expressed in most other cell types. X-linked genes were underexpressed in late spermatocytes, also as expected. Early spermatids, in contrast, had a very low number of nuclear transcripts (Figure 6F: low UMI). This was expected from the single nucleus RNA-seq strategy because, although spermatids contain many cytoplasmic mRNAs produced during the preceding spermatocyte stages, early spermatids have little active transcription. The low number of unique genes detected as being actively transcribed by snRNA-seq was used as a diagnostic in assigning identity as early elongating spermatids. However, due to noise, the relatively low signal also made it difficult to assign distinct identities within this broad designation. Both the geography of the testis UMAP and plots of gene expression in nuclei ordered by trajectory inference identified spermatids at the mid-to late elongation stage (Fig. 2C, *4), located in the blunt protruding region at the top of the “shark’s head”, as these nuclei turn on a coordinated burst of new transcription of a specific set of genes previously identified by the White-Cooper laboratory (Barreau et al. Development 2008). Trajectory inference analysis of the snRNA-seq data identified a number of additional such genes (152 in all) upregulated in late elongating spermatids, and showed that onset of their transcription is tightly coordinated Figure 6F).

Some surprising new features of the spermatocyte transcription program emerged from the data. snRNA-seq revealed that many mRNAs normally thought of as markers of somatic cells (*Upd1* and *eya*, for example) are expressed in late spermatocytes. Indeed, the cell-type specific markers, *grh* (epithelial cells), *Syt1*(neurons), *Mhc* (muscle), and *Hml* (hemocyte) shown in Figure 4 were all upregulated in late spermatocytes (see SCope). The cause of this seemingly promiscuous expression, and what role if any these genes may have in the biology of germ cells remain to be investigated. A second surprise emerged from the geography of the UMAP plot: the later stage spermatocytes split into three streams, all of which express diagnostic spermatocyte specific markers. Analysis of gene expression differences between the three parallel streams using ASAP revealed that the thinnest (leftmost) was composed of nuclei in which genes on chromosome arms 3L and 3R were expressed at relatively lower levels, while the intermediate (middle) stream had nuclei in which genes on chromosome arms 2L and 2R were expressed at lower levels than in the rightmost stream, which contained the majority of the maturing spermatocytes (see SCope). Again, the cause(s) and possible role in germ cell biology of such chromosome-wide regulation are not yet known.

The early stages of the somatic cyst cell lineage also emerged as a clear trajectory in the UMAP plot, with Leiden resolution 6 highlighting clusters representing cells known to have differing identities and roles. In the view angle shown in Fig. 2C, the early cyst cell lineage appears as a long thin line in the center (mouthpiece and upper part of the saxophone). At the head (Leiden 6 cluster 62) lie the cyst stem cells (CySCs), confirmed as proliferative by expression of *string*, the generally used homolog of the cell cycle activating phosphatase cdc25. Down along the linear stem lie post-mitotic (no *string*) early and later cyst cells (early CC1 and early CC2), presumably representing the cyst cells that enclose each cluster of transit amplifying spermatogonia. The cyst cell lineage then splits into two streams, which likely represent cyst cells associated with spermatocytes. Trajectory inference analysis showed that different genes are more strongly upregulated in the two streams of spermatocyte associated cyst cells (hence the split) and also confirmed the relatedness of the two streams, as each featured a ‘shadow’ of many of the genes up-regulated to a higher degree in the other (Fig. 6G).

Developmental progression of the different Leiden 6 clusters that form the germ line and cyst stem cell lineages emergent from the UMAP plot was confirmed in annotation jamborees through expression of known stage specific markers, many of which had been previously identified genetically and studied functionally by laboratories investigating the *Drosophila* testis. The FCA data identify many additional genes differentially up or down regulated at specific steps in these stem cell lineages, presenting a unique opportunity for functional tests to identify new players in the transitions from one cell state to the next.

As described above, a small cluster of epithelial-like cells termed the hub, located at the apical tip of the testis, is a key player in the organization and function of the testis as a reproductive organ. The hub cells provide close range signals required to maintain stem cell state in the associated germ line and cyst cell lineages, BMP and Upd1 respectively. Represented by only 79 nuclei (Fig. 2C, germinal proliferation center hub) in the snFCA V1 testis dataset, the hub cells express many markers common to epithelial cells, representing their cell biological arrangement and attachments to each other, and to cyst cells, likely reflecting their embryonic co-origin and ability to interconvert with cyst cells after drastic injury (Hétié et al., 2014). The hub cells differ from cyst cells and most other epithelial cells, however, in their diagnostic expression of the cytokine-like ligand Upd1. The UMAP for testis presented in Fig. 2C is plotted using *Relaxed* version of the snFCA 10X data because the important hub cell cluster was filtered out by the algorithms used to generate the *Stringent* FCA datasets, likely due to the expression of many “somatic” transcripts including Upd1 in late spermatocytes. This same expression of “somatic” transcripts also caused many late spermatocytes to be filtered out by the algorithms used to generate the *Relaxed* FCA dataset.

## Supplementary Figures and Legends

**Figure S1.**
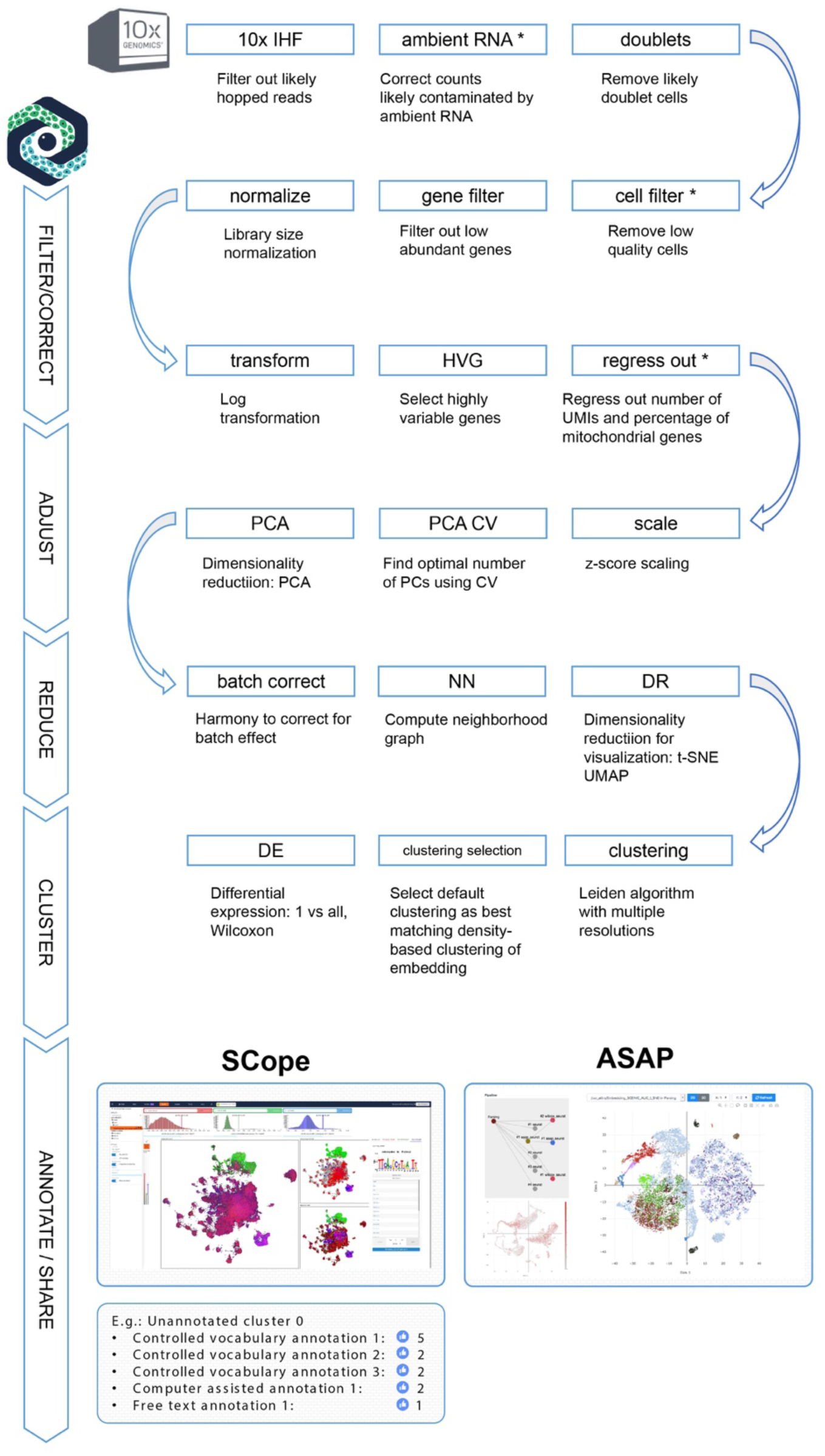
**Steps for 10x data processing, from raw sequencing data to cluster analysis.** Processed data are annotated and shared through SCope and ASAP. See Methods for detailed description.

**Figure S2.**
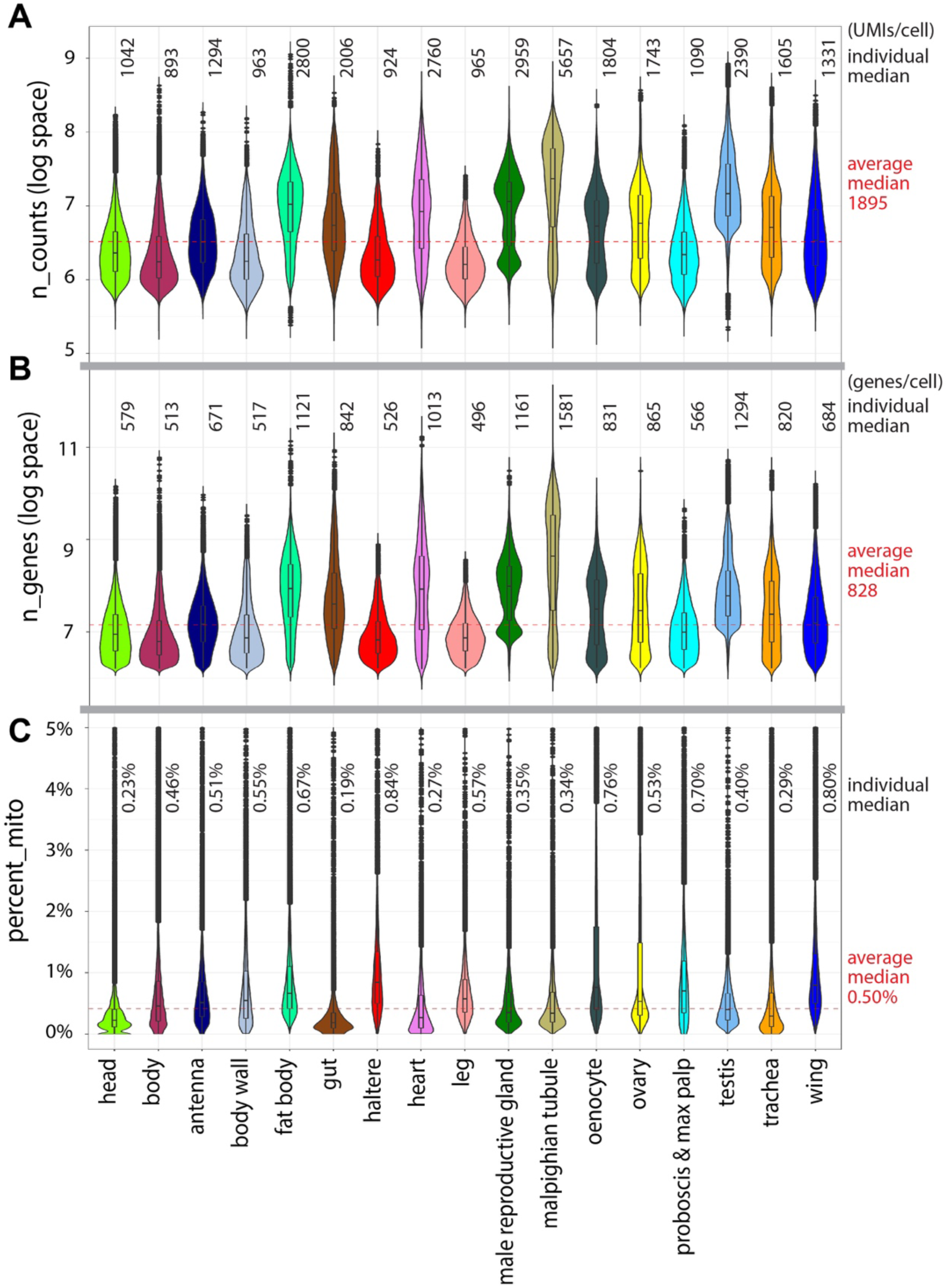
**Quality control of 10x data.** **(A)** The average median UMI count is 1895 UMIs per cell. The log space is log_e_. **(B)** The average median gene detection is 828 genes per cell. The log space is log_e_. **(C)** The average median percentage of mitochondrial genes for all samples is 0.50%. Numbers for individual tissues are indicated.

**Figure S3.**
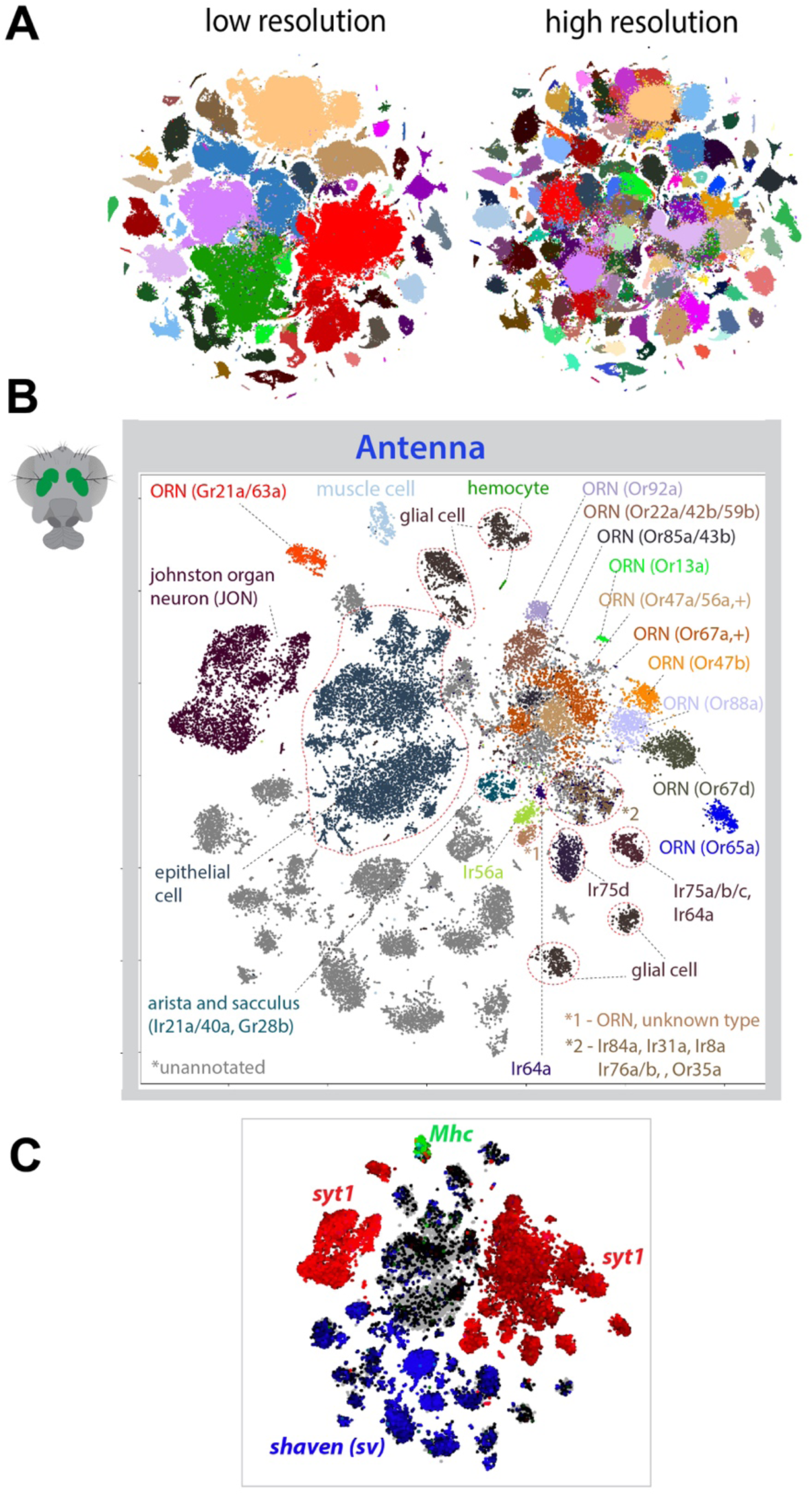
**Different clustering resolutions and cell type annotation in the antenna.** **(A)** Different clustering resolutions (using Leiden) are used for annotating cell types, because some cell types are present at low clustering resolution, and others appear only at higher resolution. **(B)** tSNE plot with annotations for the fly antenna from the *Stringent* 10x dataset. All three antennal segments were dissected for single-nucleus sequencing. ORN: olfactory receptor neuron. **(C)** The unannotated clusters in the antenna are largely *shaven+,* likely to be different types of non-neuronal and non-glial supporting cells from different segments of the antenna.

**Figure S4.**
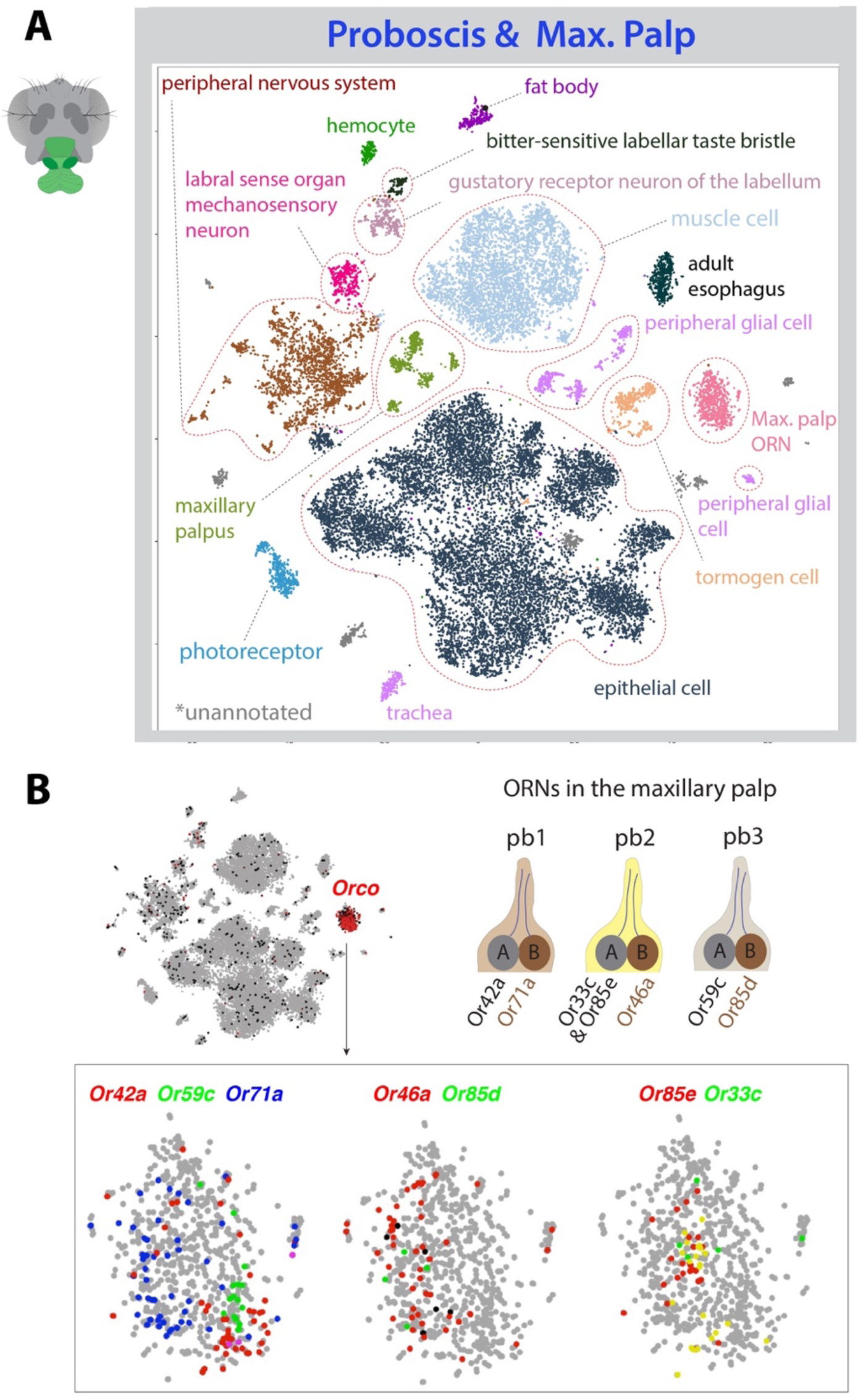
**Cell type annotation in the proboscis and maxillary palp.** **(A)** tSNE plot with annotations for the fly proboscis and maxillary palp from the *Stringent* 10x dataset. ORN: olfactory receptor neuron. **(B)** Expression of olfactory receptor co-receptor (*orco*) in one cluster annotated as maxillary palp ORNs. All 7 known olfactory receptor genes can be detected, including two receptors, Or33c and Or85e, that are co-expressed in the same ORN <u>(Bai et al., 2009)</u>. Palpal basiconic 1 (pb1), pb2, and pb3 are three different types of sensilla in the maxillary palp.

**Figure S5.**
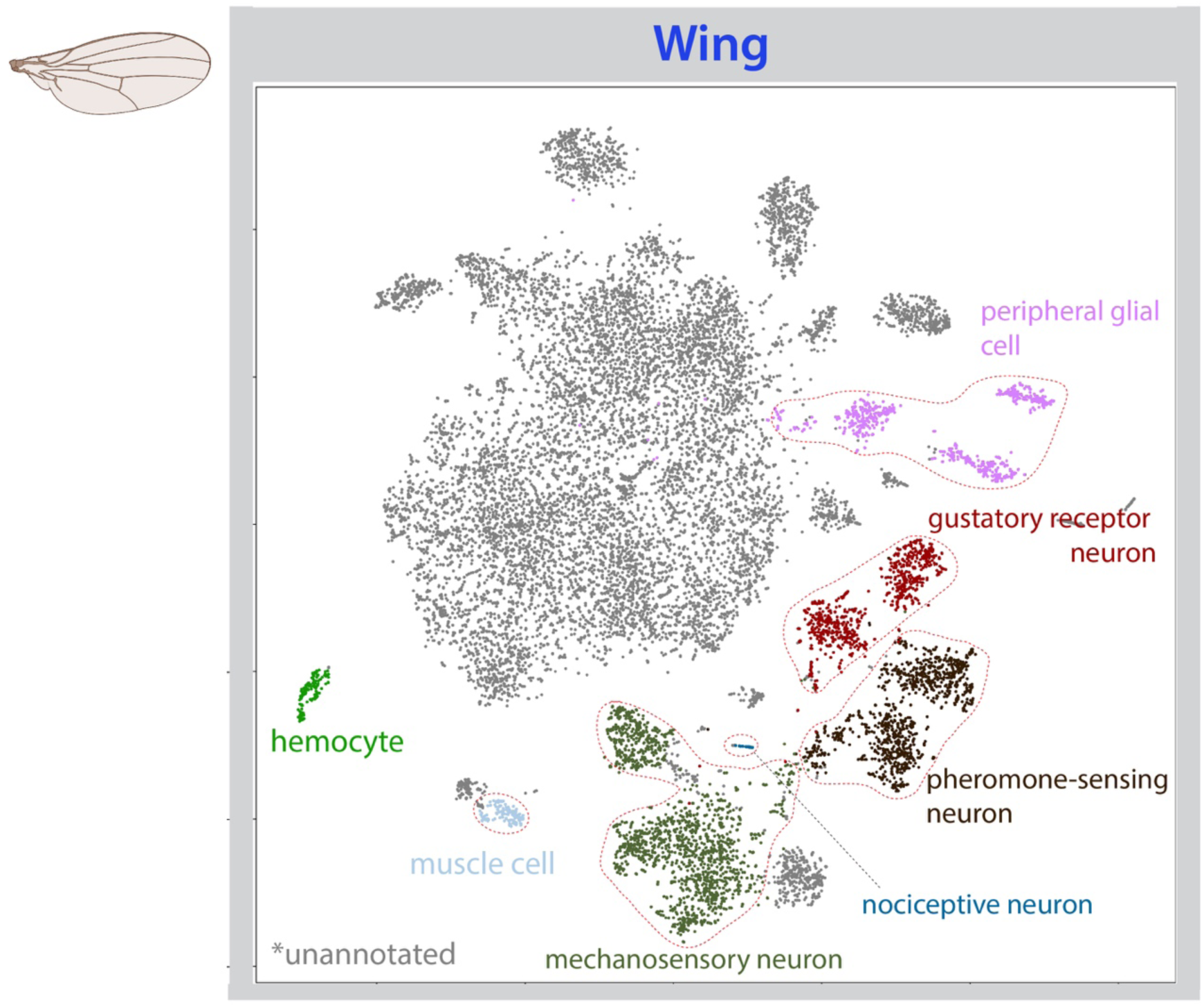
**Cell type annotation in the wing.** tSNE plot with annotations for the fly wing from the *Stringent* 10x dataset. Note a large group of cells are currently unannotated, which are likely to be epithelial cells.

**Figure S6.**
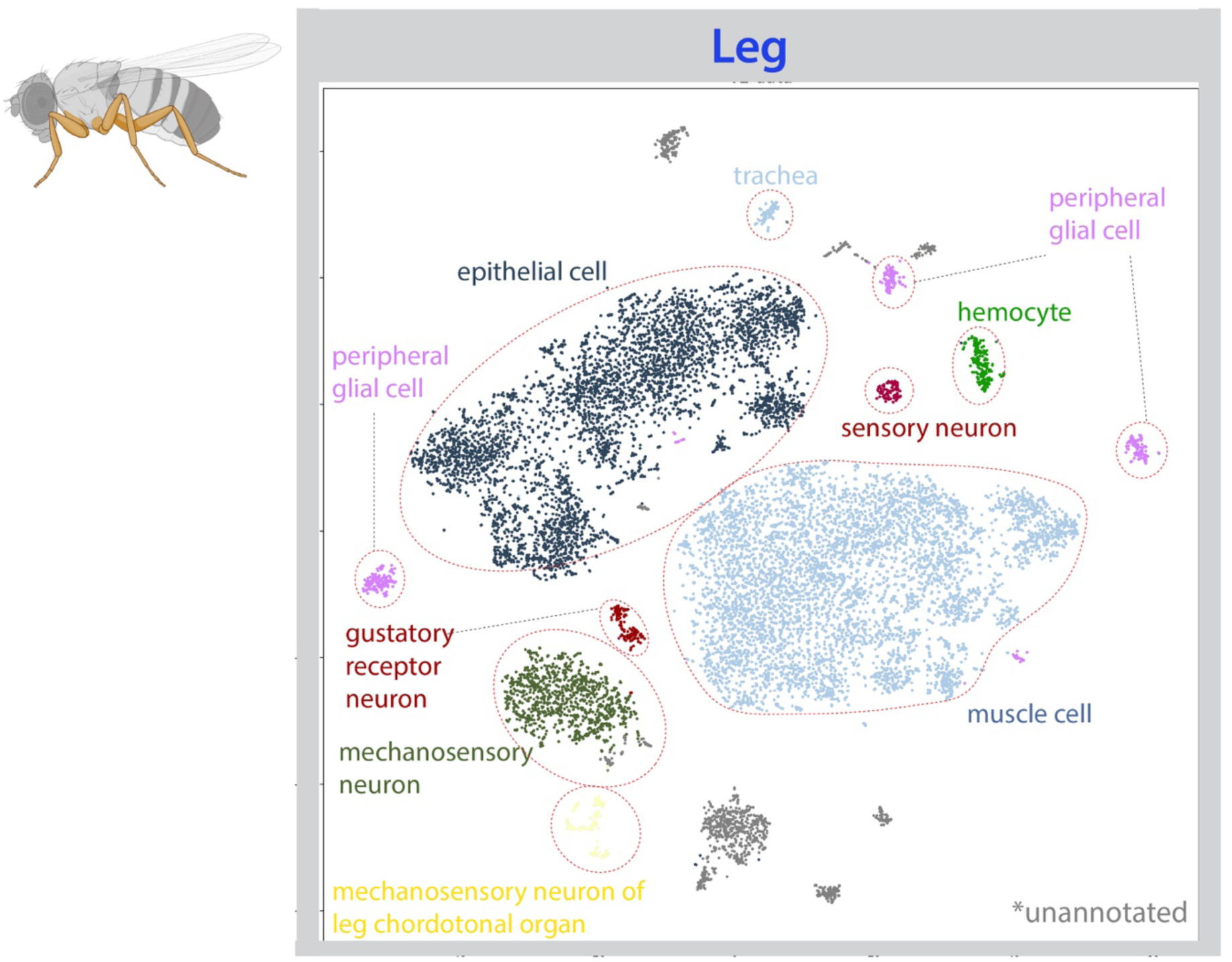
**Cell type annotation in the leg.** tSNE plot with annotations for the fly leg from the *Stringent* 10x dataset. All six fly legs were dissected and pooled for single-nucleus sequencing.

**Figure S7.**
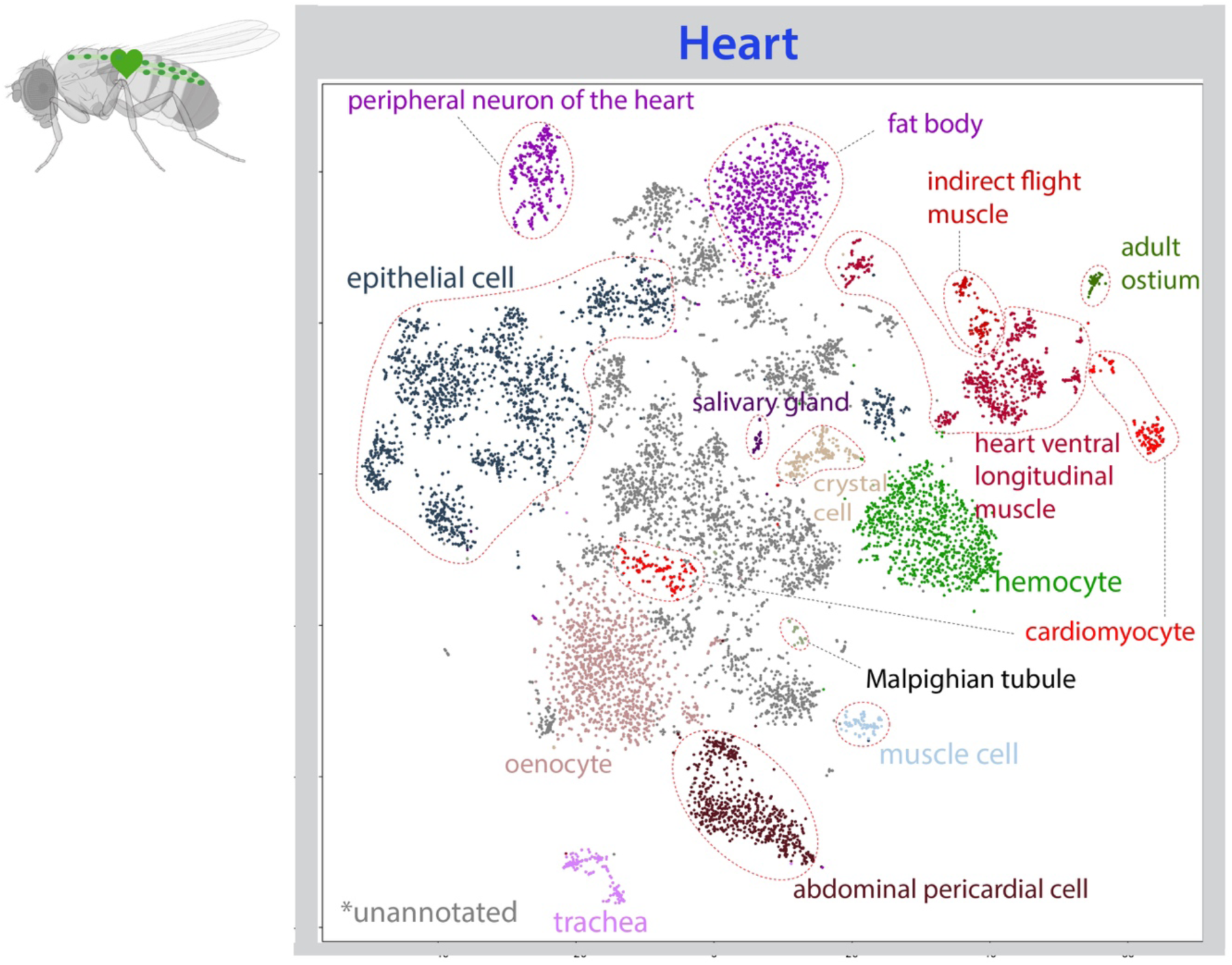
**Cell type annotation in the heart.** tSNE plot with annotations for the fly heart from the *Stringent* 10x dataset.

**Figure S8.**
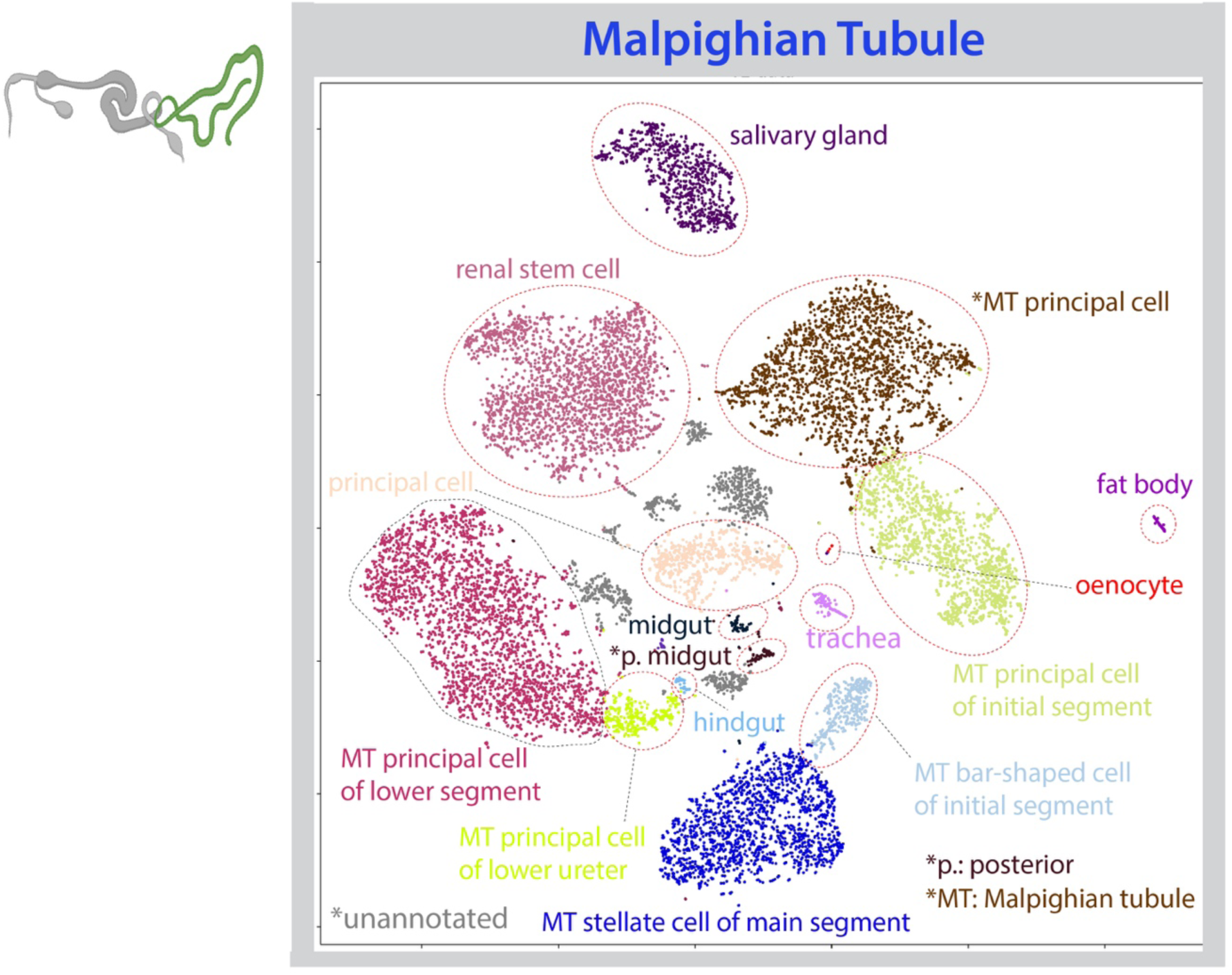
**Cell type annotation in the Malpighian tubule.** tSNE plot with annotations for the fly Malpighian tubule (MT) from the *Stringent* 10x dataset.

**Figure S9.**
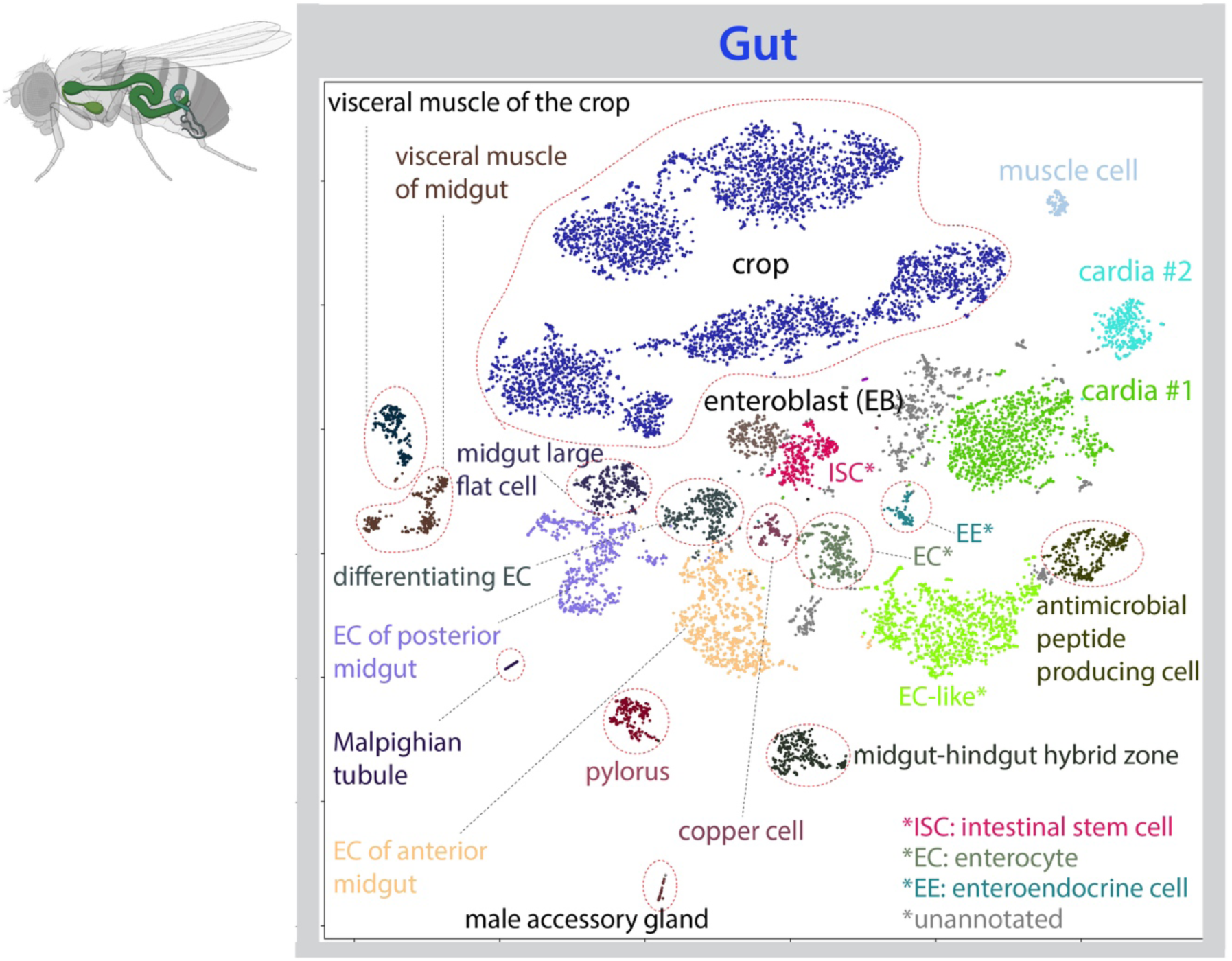
**Cell type annotation in the gut.** tSNE plot with annotations for the fly gut from the *Stringent* 10x dataset.

**Figure S10.**
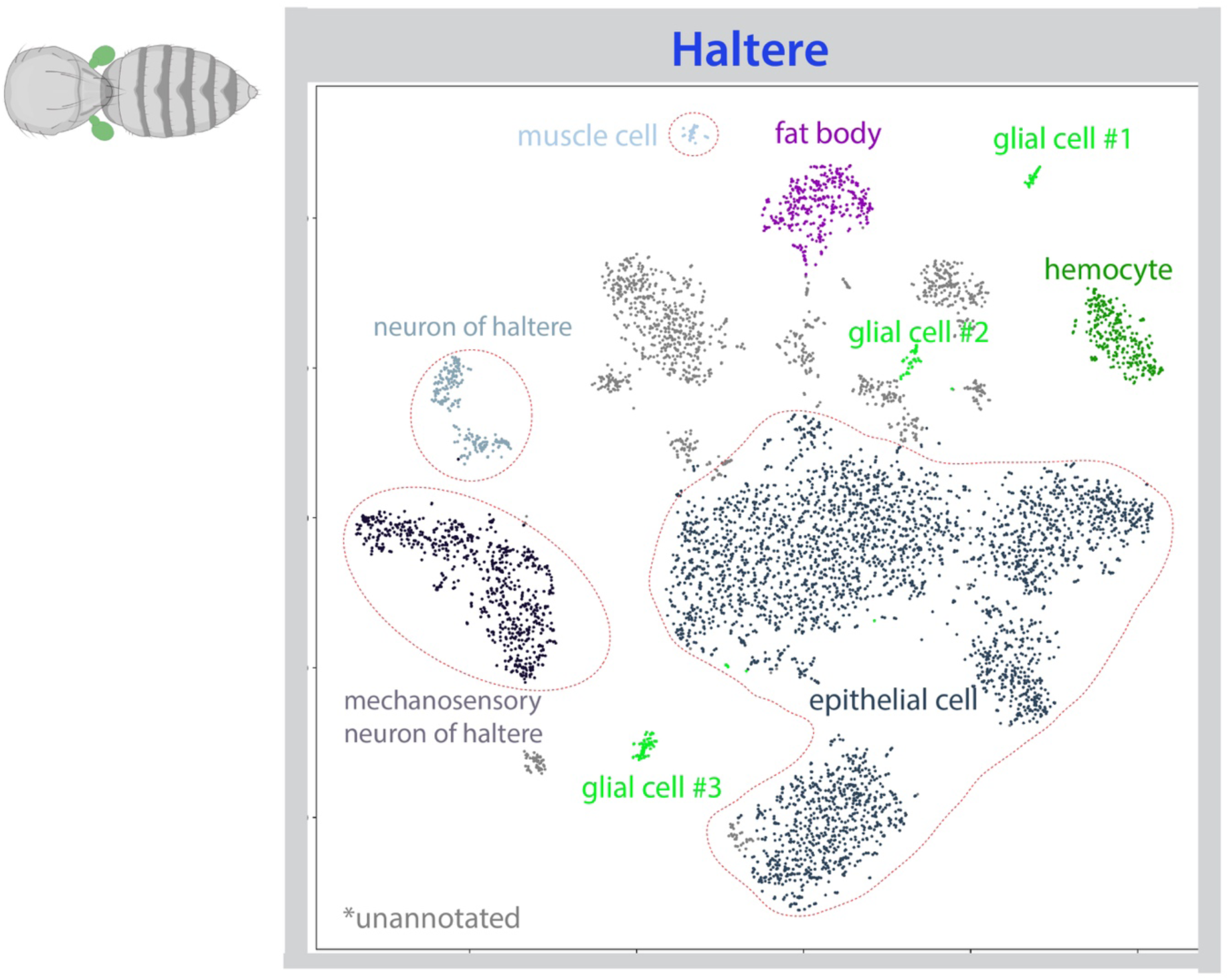
**Cell type annotation in the haltere.** tSNE plot with annotations for the fly haltere from the *Stringent* 10x dataset.

**Figure S11.**
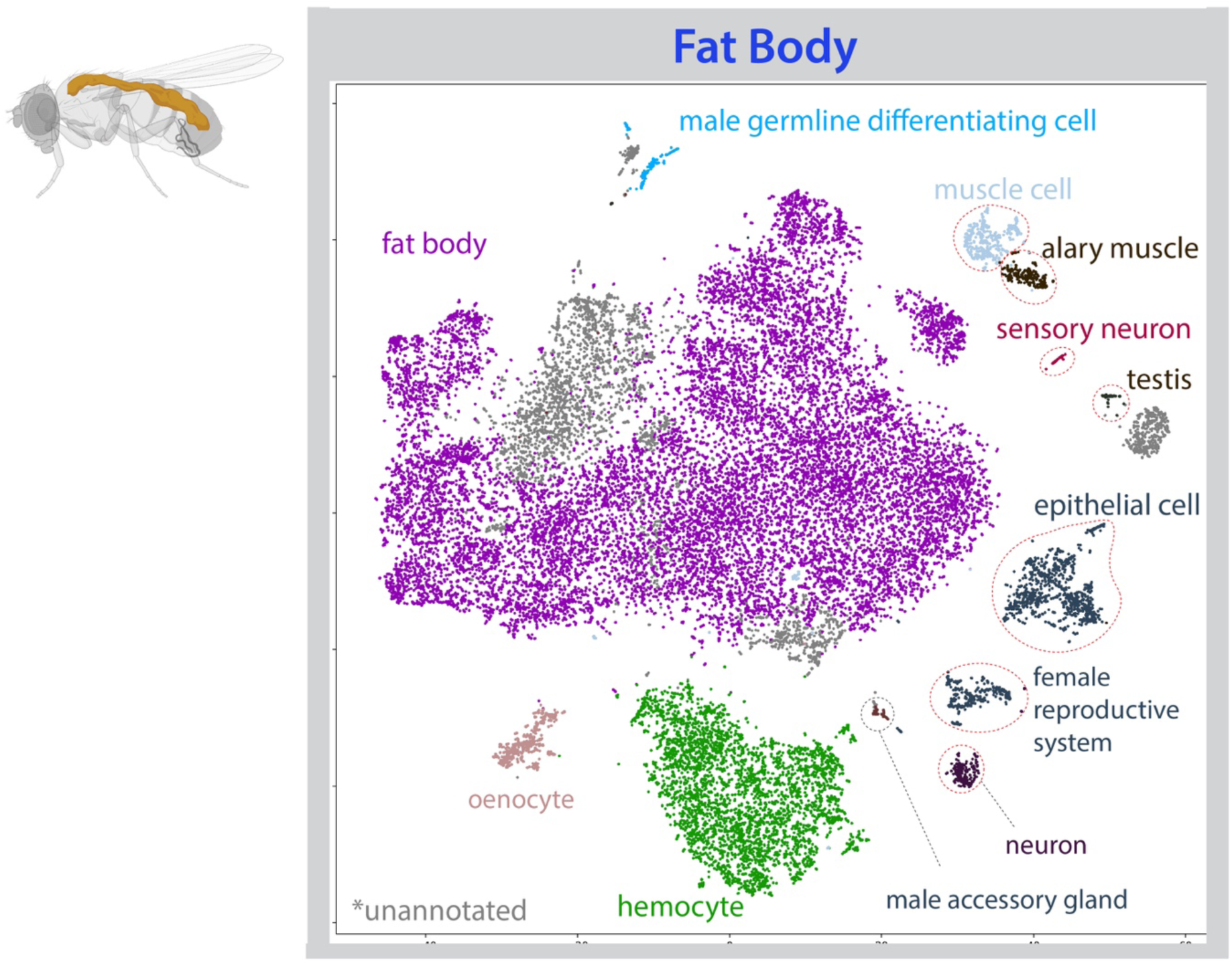
**Cell type annotation in the fat body.** tSNE plot with annotations for the fly fat body cells from the *Stringent* 10x dataset. Fat body cells are FAC-sorted based on the nuclear GFP signal; flies are *Cg-GAL4 > UAS-lamGFP*.

**Figure S12.**
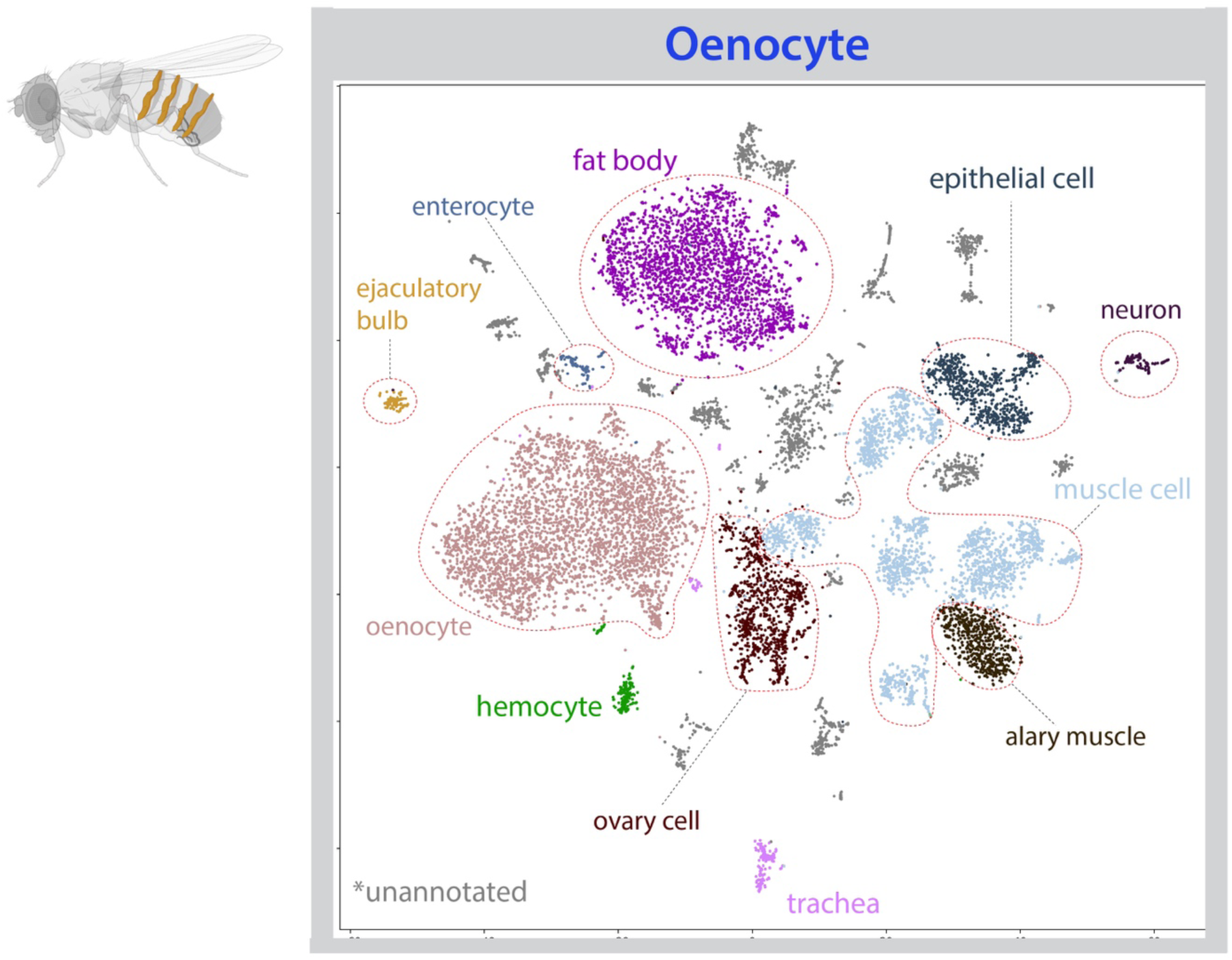
**Cell type annotation in the oenocyte.** tSNE plot with annotations for the fly oenocyte from the *Stringent* 10x dataset. Oenoctyes are FAC-sorted based on the nuclear GFP signal; flies are *PromE800-GAL4 > UAS-unc84GFP*.

**Figure S13.**
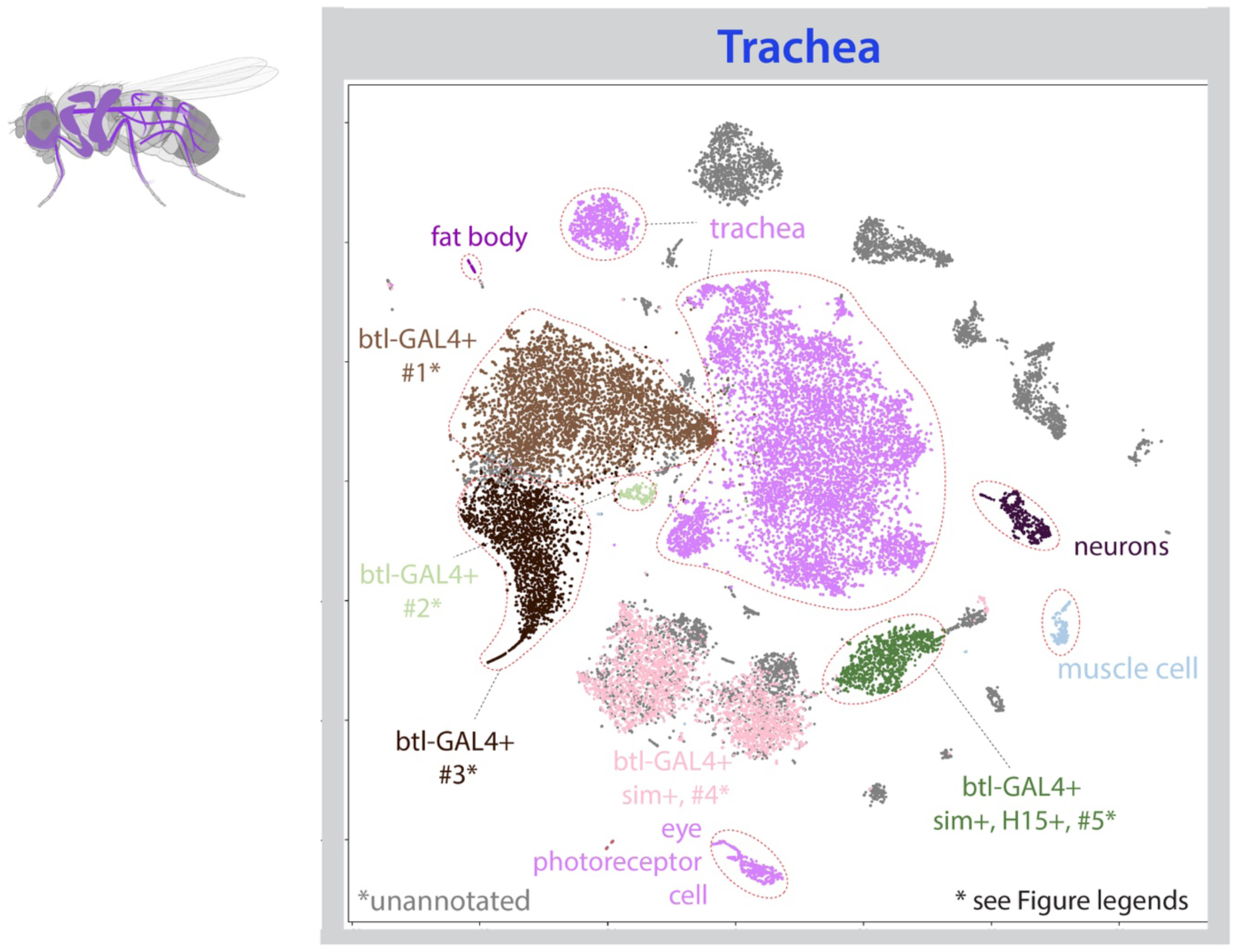
**Cell type annotation in the trachea.** tSNE plot with annotations for the fly trachea from the *Stringent* 10x dataset. Tracheal cells are FAC-sorted based on the nuclear GFP signal; flies are *btl-GAL4 > UAS-lamGFP*.

**Figure S14.**
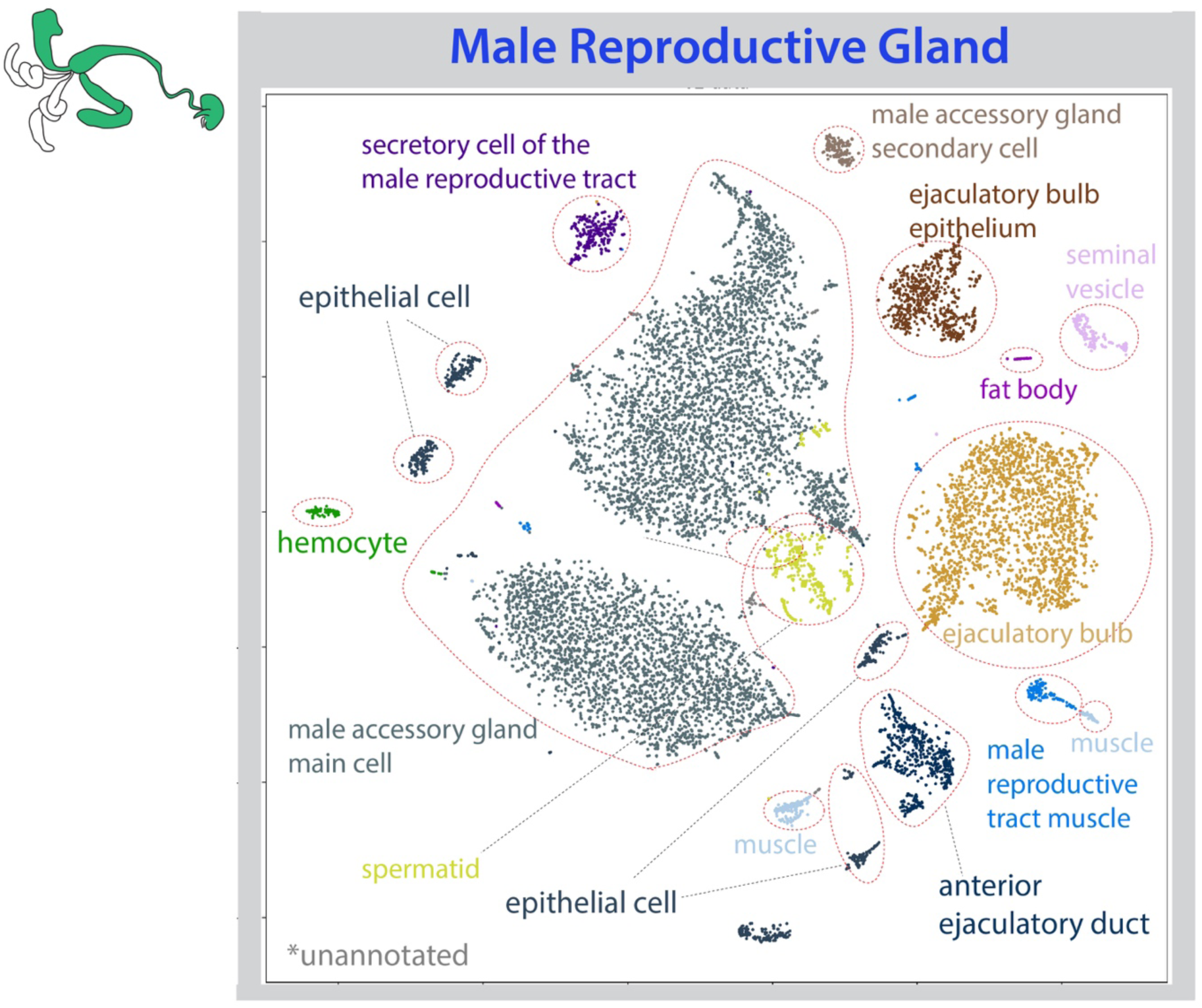
**Cell type annotation in the male reproductive gland.** tSNE plot with annotations for the fly male reproductive gland from the *Stringent* 10x dataset. The sequenced cells are from dissected male accessory glands, ejaculatory ducts and ejaculatory bulbs.

**Figure S15.**
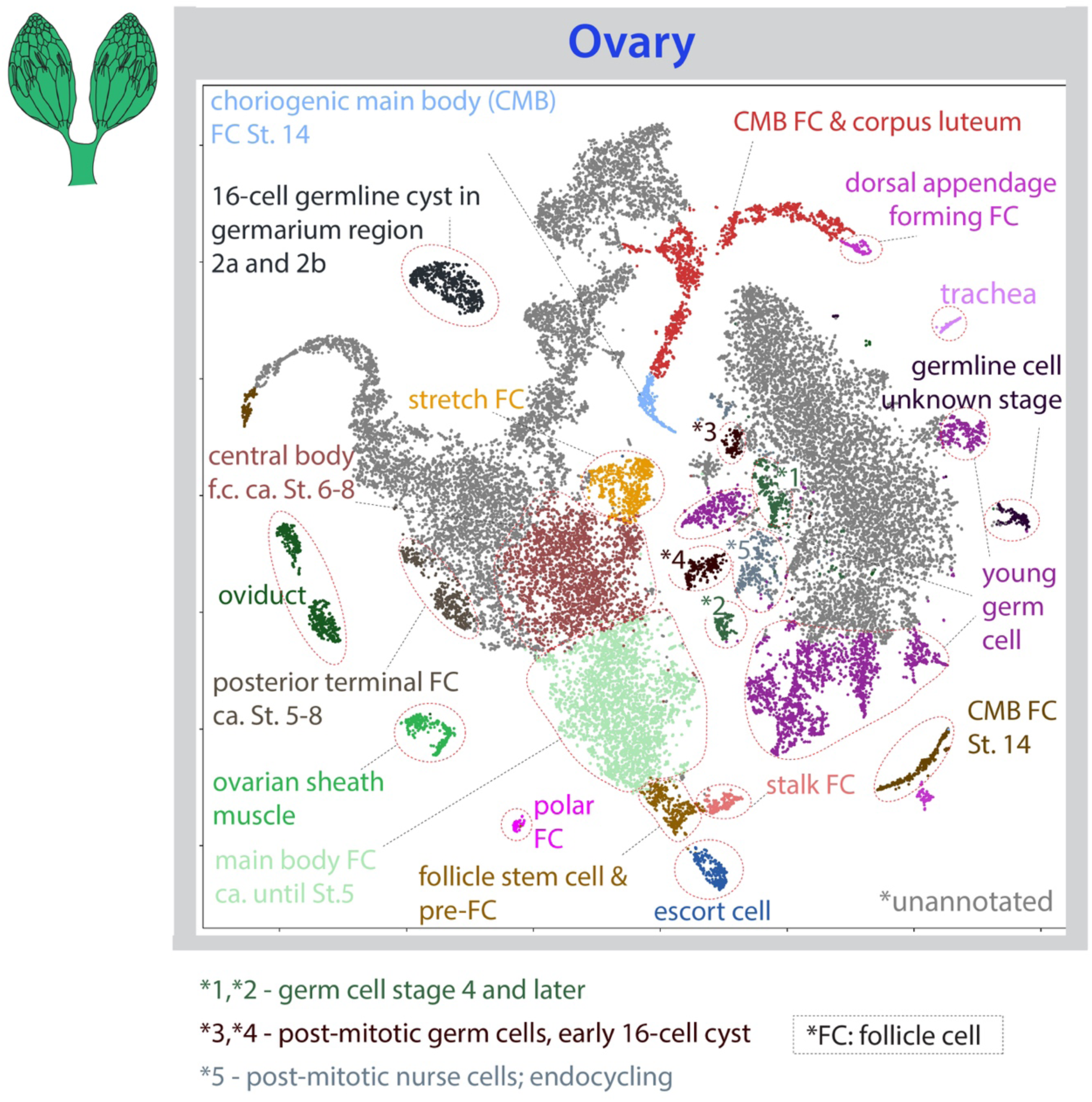
**Cell type annotation in the ovary.** tSNE plot with annotations for the fly ovary from the *Stringent* 10x dataset.

**Figure S16.**
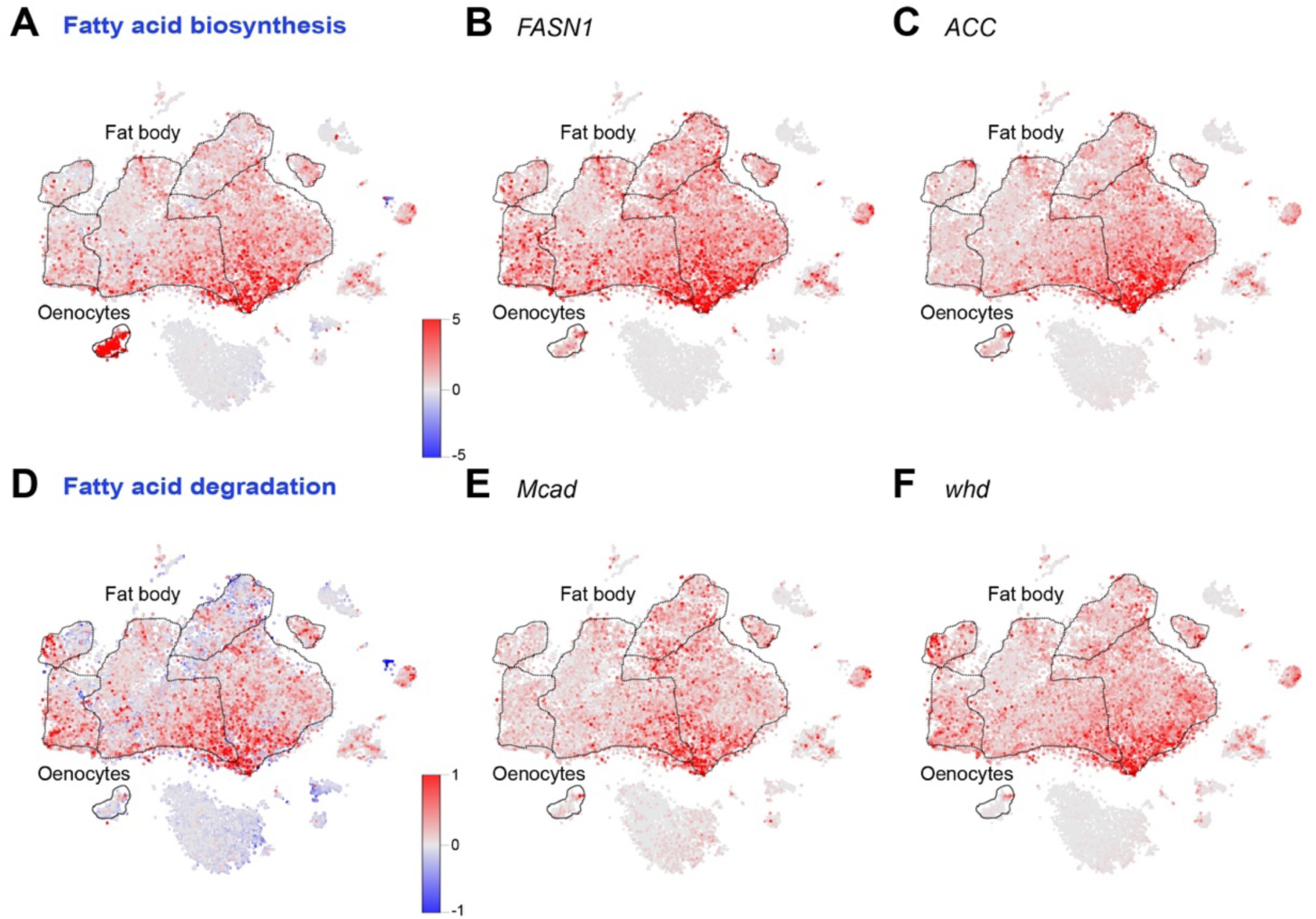
**Metabolic pathway enrichment reveals the existence of cell subpopulations suggesting tissue specific functional specialization.** **(A)** Fatty acid biosynthesis pathway enrichment analysis as performed by ModuleScore (see Methods) in ASAP reveals strong homogeneous positive enrichment in oenocytes, while the fat body shows non-homogeneous enrichment across all cells. Red colors correspond to positive enrichment while blue colors correspond to negative enrichment. **(B, C)** Similar profiles were obtained if using single genes in this pathway, *FASN1* and *ACC*. **(D)** Fatty acid degradation pathway enrichment as revealed by ModuleScore analysis in ASAP shows low enrichment in oenocytes, while in the fat body there is a positive enrichment in specific subpopulations of cells. Red colors correspond to positive enrichment while blue colors correspond to negative enrichment. **(E, F)** Similar profiles were obtained if using single genes in this pathway, *Mcad* and *whd*.

**Figure S17.**
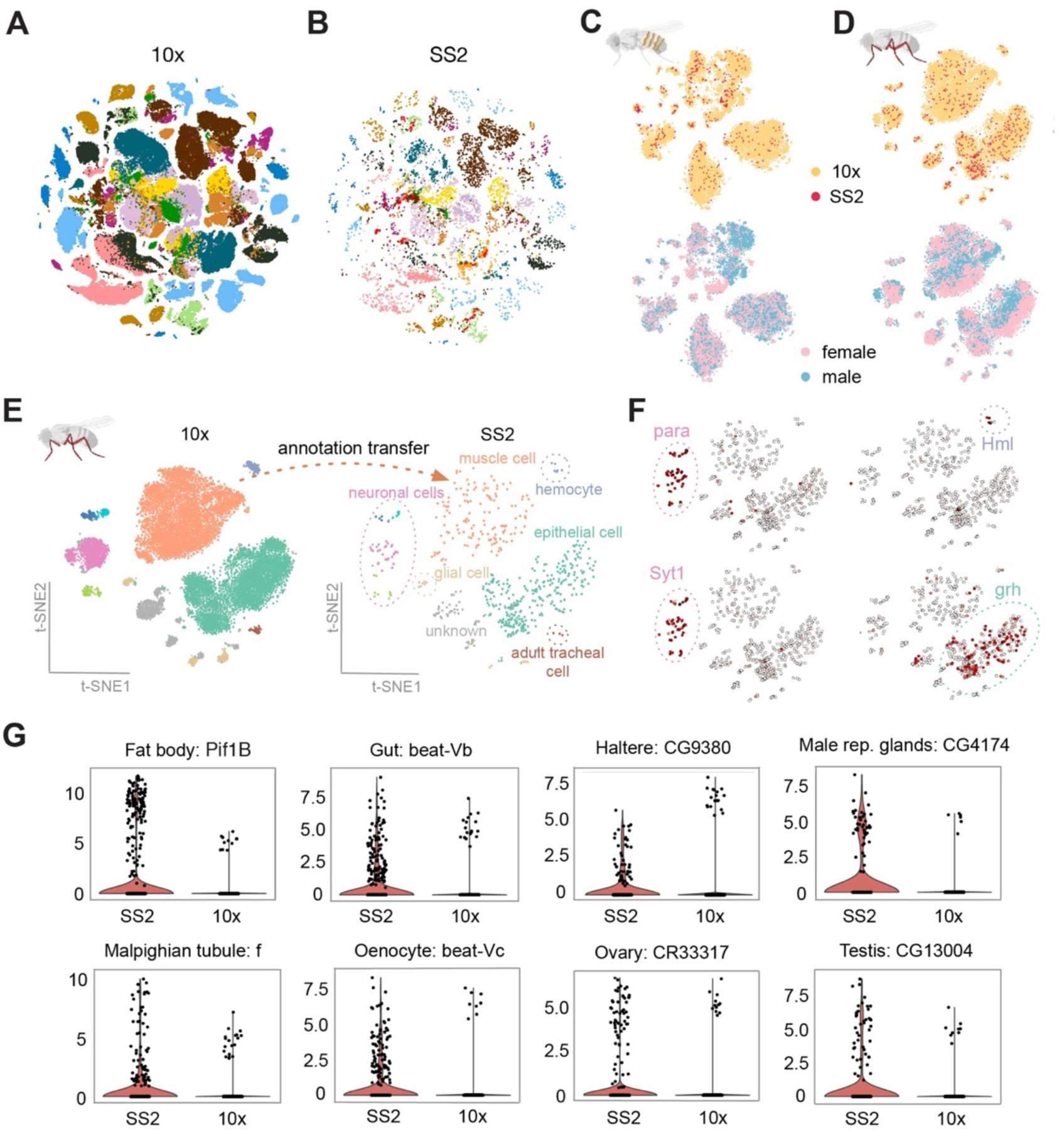
**Integration of Smart-seq2 (SS2) and 10x Genomics data.** **(A)** tSNE visualization of individually dissected tissues using 10x Genomics and integrated with Smart-seq2 data. Colors denote different tissues. **(B)** tSNE visualization of individually dissected tissues using Smart-seq2 and integrated with 10x Genomics data. Colors denote different tissues. **(C,D)** Examples of integrated data for (C) oenocyte, and (D) leg. Cells are colored by technology (top) and by gender (bottom). **(E)** Overview of computational pipeline for annotating Smart-seq2 data using leg as an example. After integrating 10x Genomics and Smart-seq2 data, we train a classifier on 10x Genomics data (left) and transfer annotations to Smart-seq2 data (right). Colors indicate different cell types. **(F)** Validation of Smart-seq2 annotations by known marker genes. Cells annotated as neuronal cells correctly express *para* and *Syt1* neuronal markers, while cells annotated as hemocyte and epitelial correctly express *Hml* and *grh* markers, respectively. **(G)** Examples of genes expressed in Smart-seq2 cells, but their expression is barely captured with 10x Genomics.

**Figure S18.**
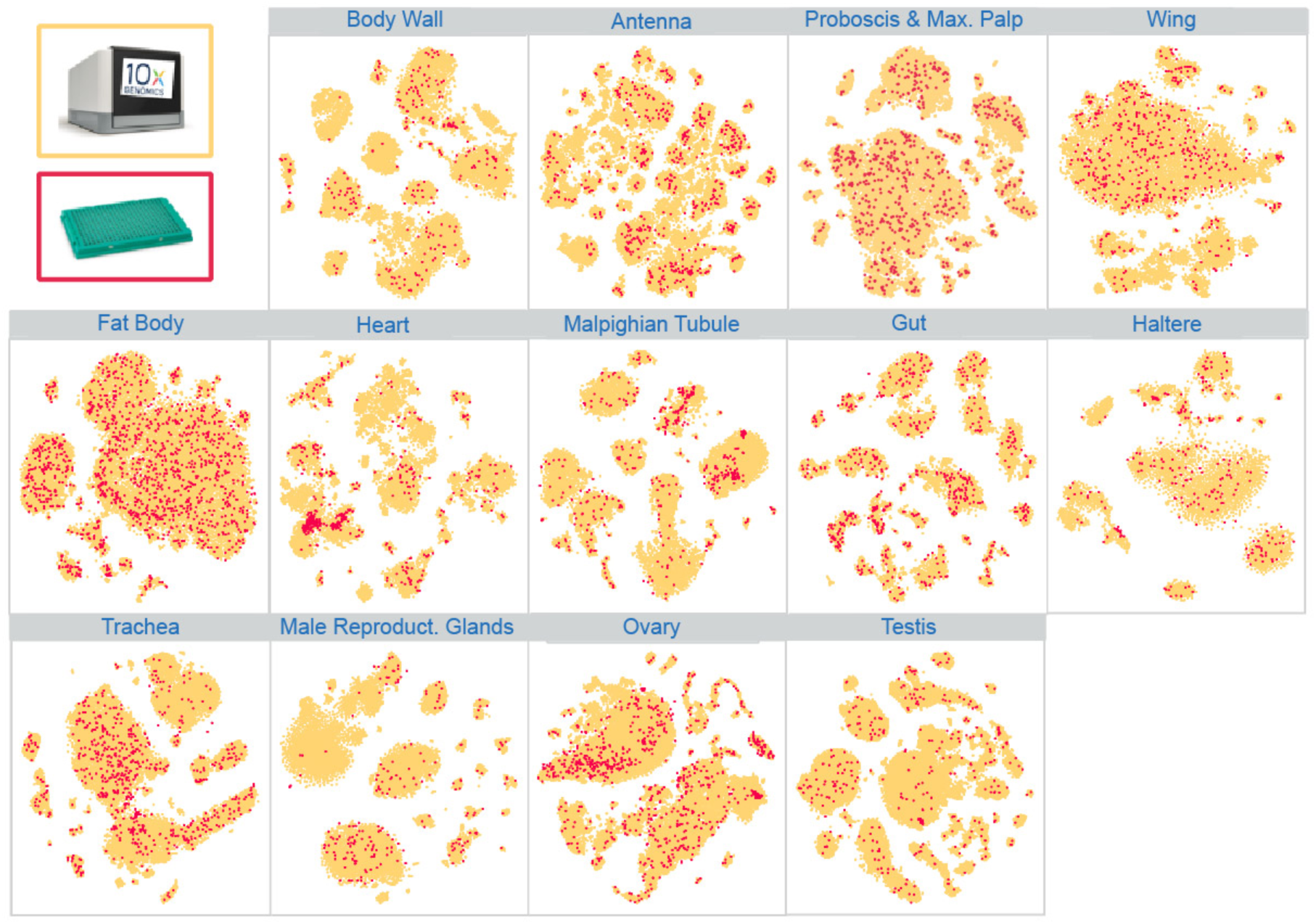
**Tissue-level integration of 10x Genomics and Smart-seq2 datasets.** tSNE visualizations of 13 individually dissected tissues. Yellow color denotes cells from 10x Genomics and red color denotes cells from Smart-seq2. Remaining tissues (oenocyte and leg) are visualized in fig. **S17 C,D.**

**Figure S19.**
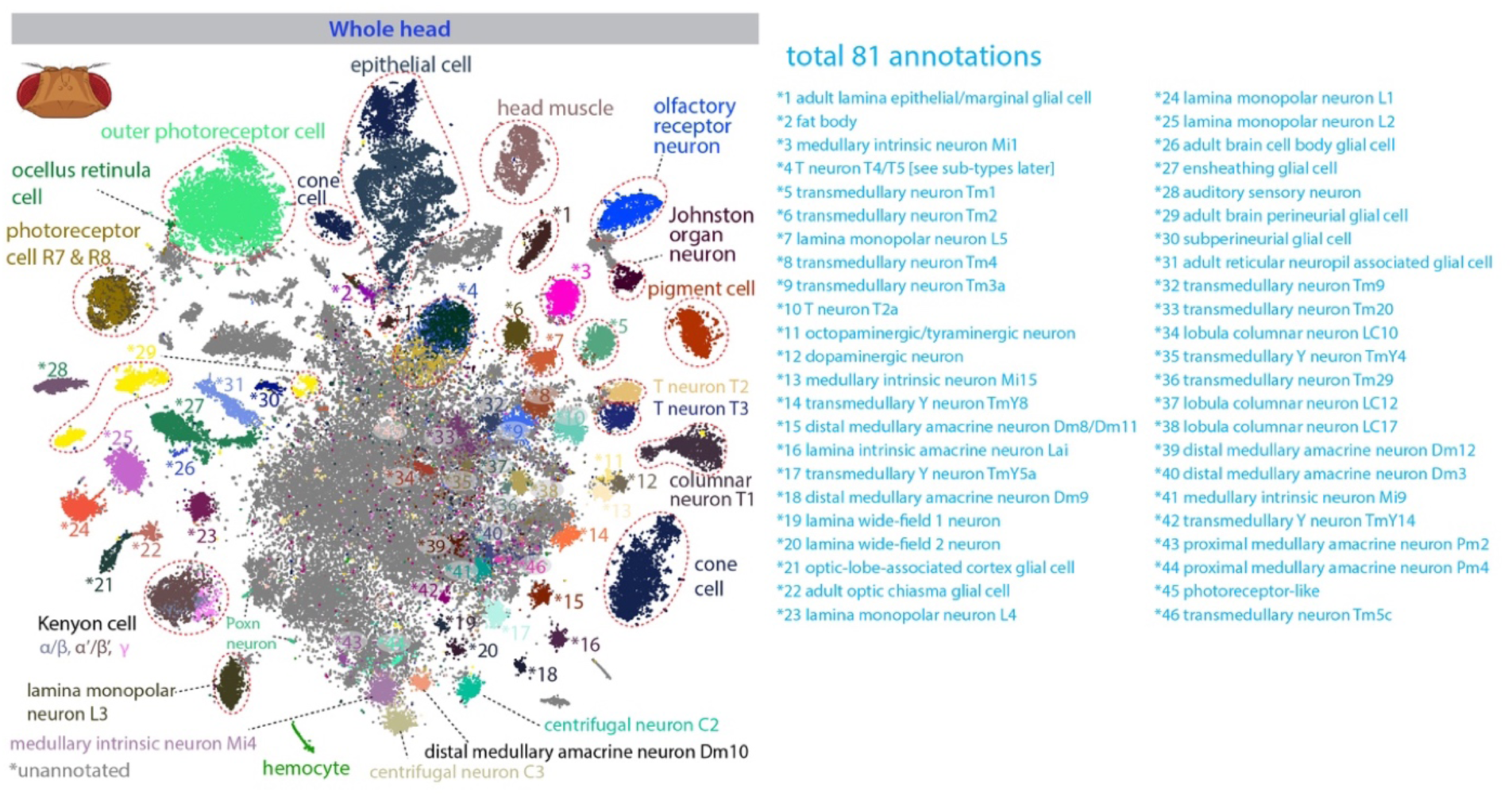
tSNE plot with annotations for the fly head from the *Stringent* 10x dataset. A large number of cells in the middle are unannotated cells, most of which are neurons from the central brain. The annotations not indicated in the plot are listed on the right.

**Figure S20.**
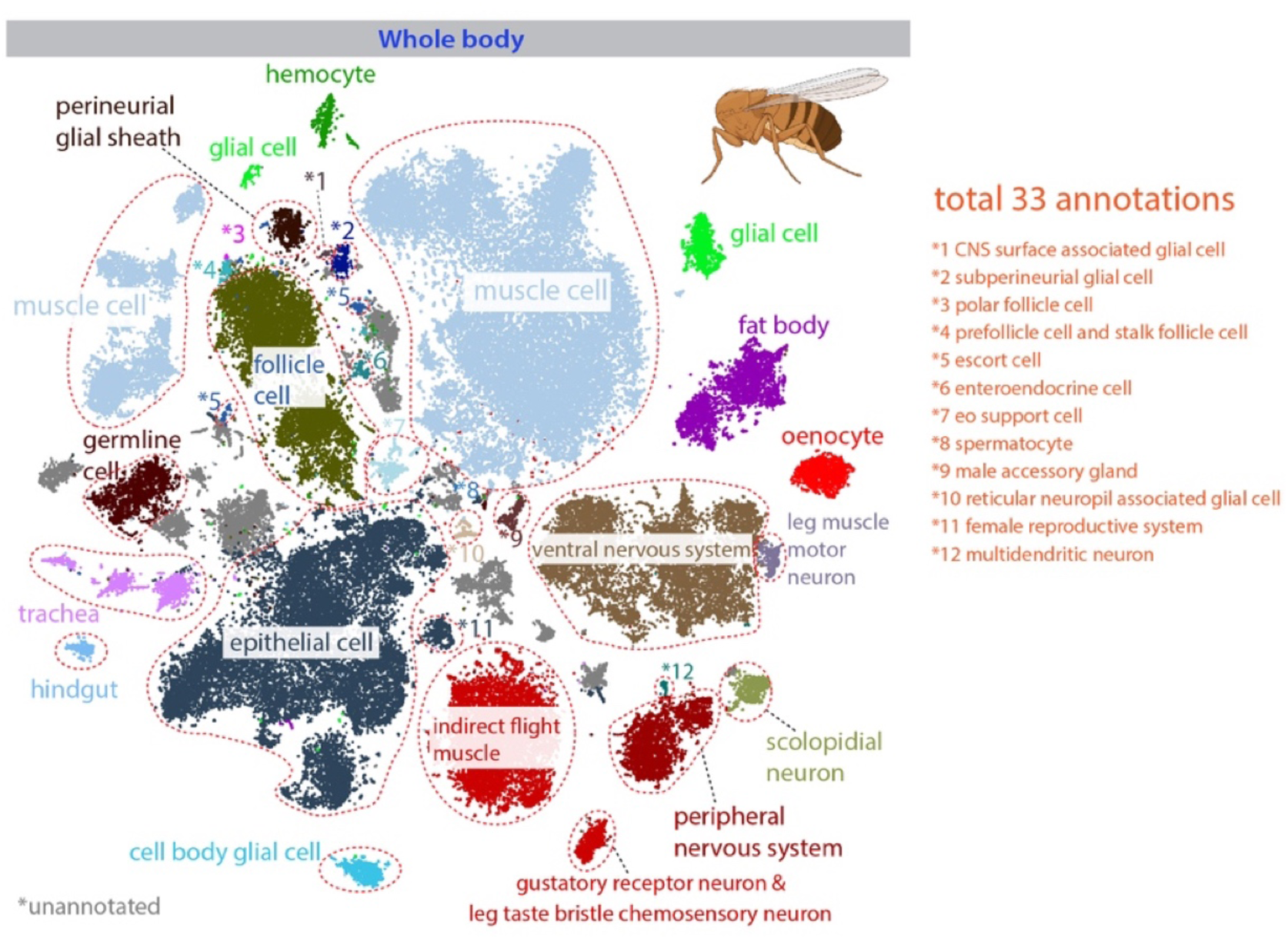
tSNE plot with annotations for the fly body from the *Stringent* 10x dataset. Note that only the top abundant cell types are annotated, and many of them can be further divided into different subtypes. The annotations not indicated in the plot are listed on the right.

**Figure S21.**
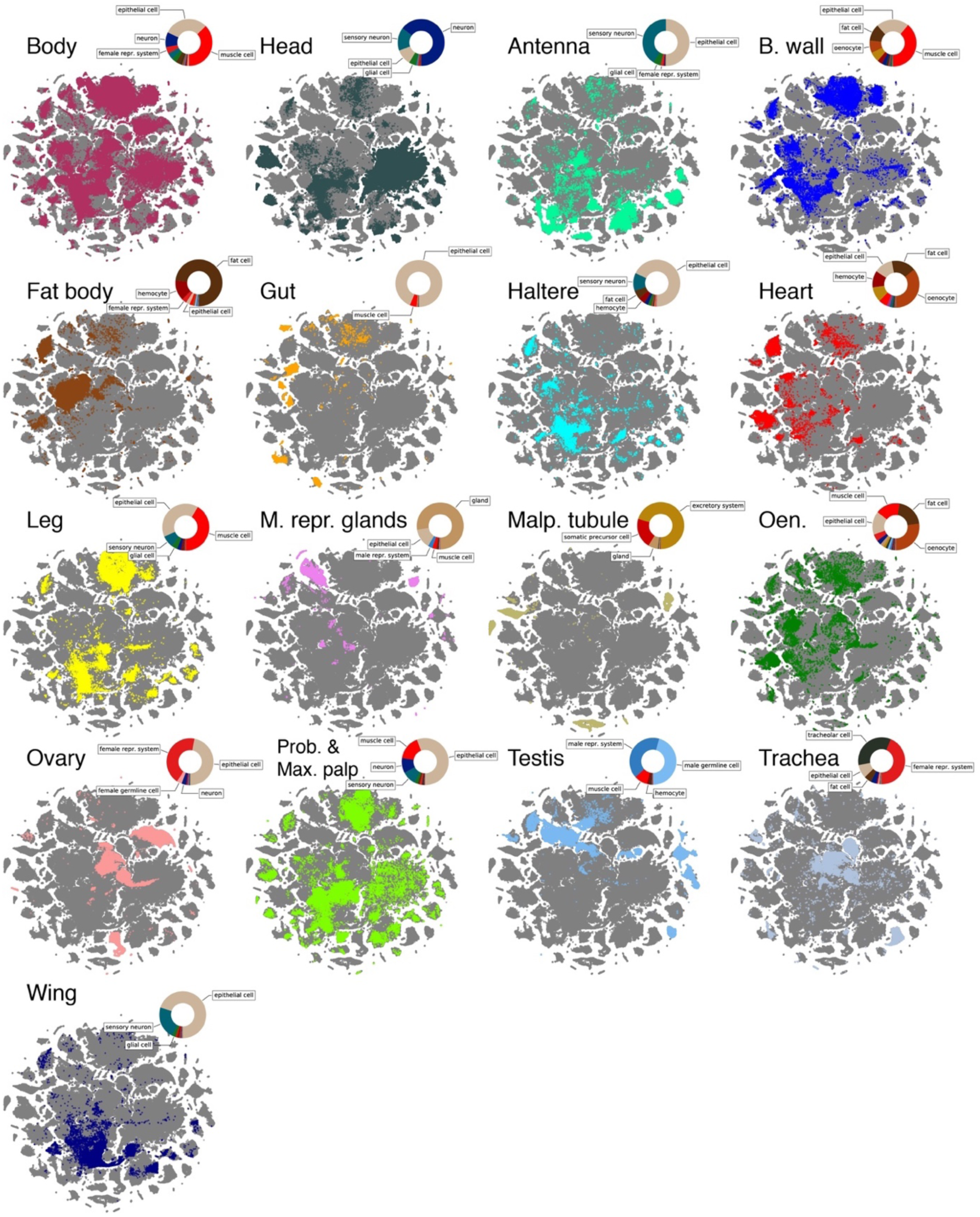
**tSNE plots of all cells from the *Stringent* 10x dataset.** Each tissue is highlighted in a different color. Pie charts show the top common cell types for each tissue, such as epithelial cells, neurons, and muscle cells, and so on. Note that some cells from two tissues overlap in one cluster, indicating these two tissues share one cell type. For example, the head and body share cell types, such as muscle, CNS neurons, and epithelial cells (see fig. S22). B. wall for body wall; M. repr. glands for male reproductive glands; Malp. tubule for Malpighian tubule; prob. max. palp for proboscis and maxillary palp.

**Figure S22.**
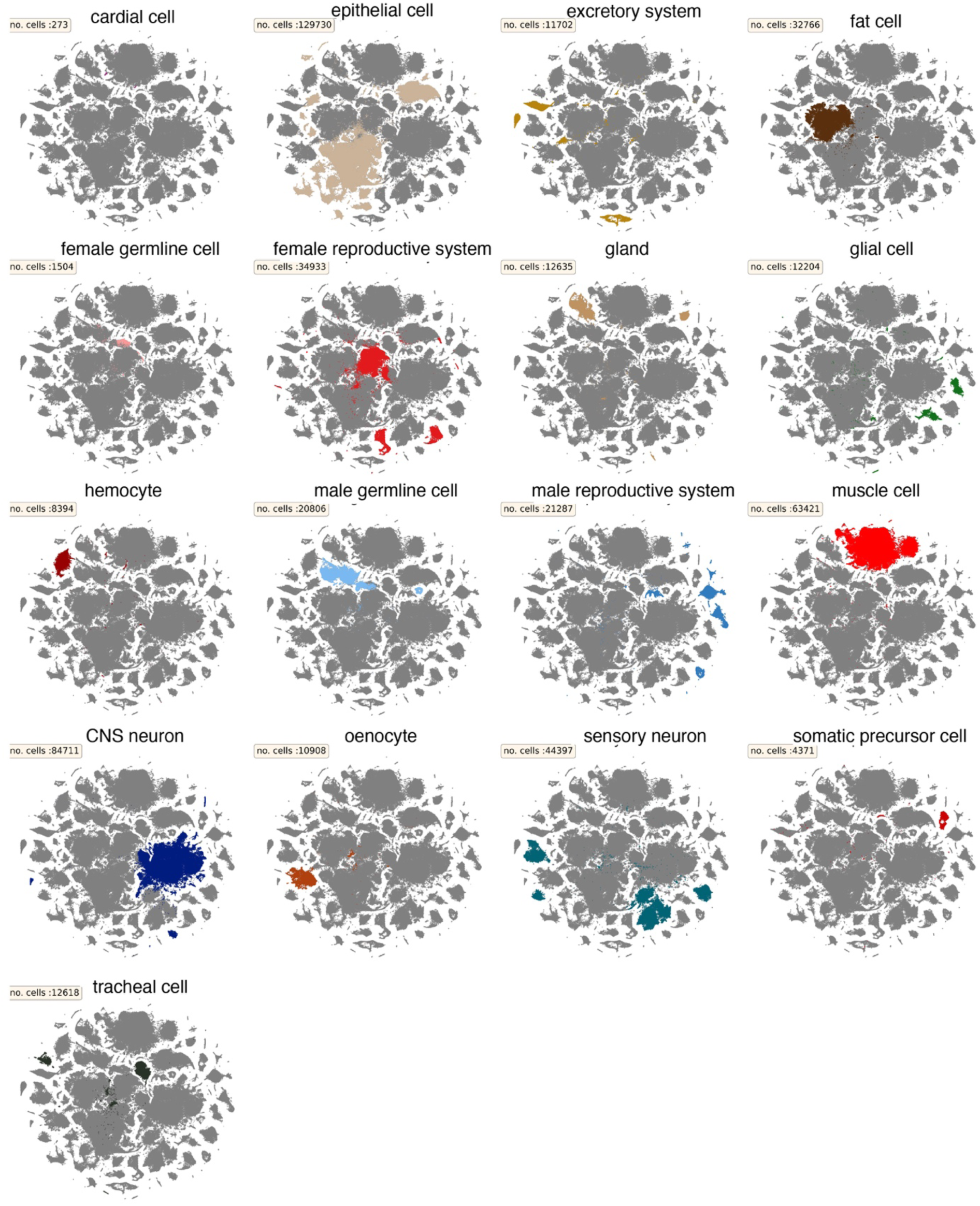
tSNE plots of all cells from the *Stringent* 10x dataset. Cell types from broad categories are highlighted and cell numbers are indicated.

**Figure S23.**
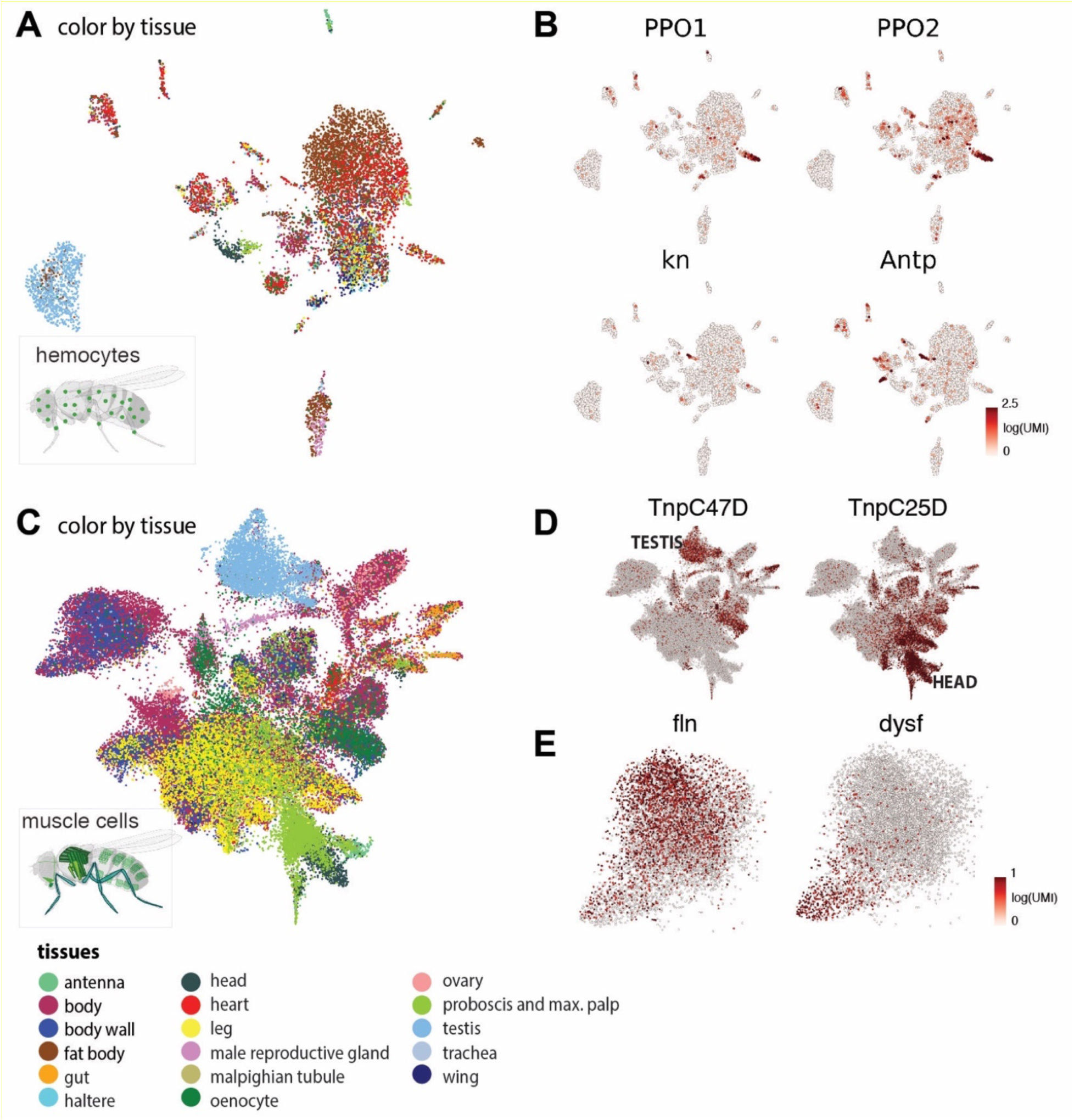
**Cross-tissue analysis of hemocytes and muscle cells.** **(A)** tSNE plot of all annotated hemocytes colored by tissue types. **(B)** Expression of *PPO1* and *PPO2* labeling crystal cells. Expression of *Antp* and *kn labeling* the presumptive lymph gland posterior signaling center. **(C)** tSNE plot of all annotated muscle cells colored by tissue types. **(D)** Expression of *TnpC47D* and and *TnpC25D* in all annotated muscle cells. **(E)** Expression of *fln* and *dysf* showing gradients in the indirect flight muscle cluster.

**Figure S24.**
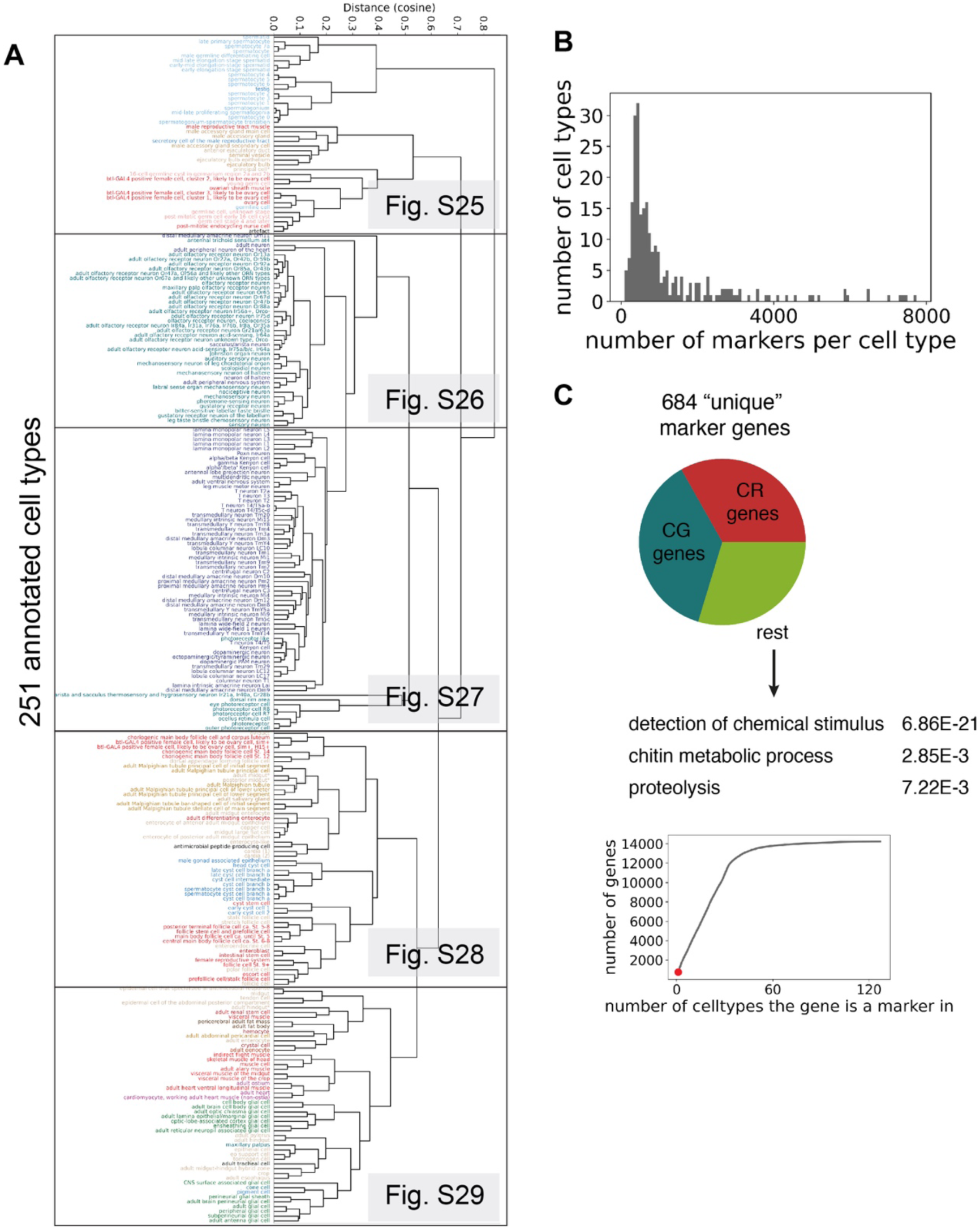
**Organism-wide gene expression comparison** **(A)** Dendrogram showing the cosine similarity of the transcriptomes of the different annotated cell types. Cell types are colored based on broad classes. Enlarged details shown in fig. S25-S29. **(B)** Histogram showing the number of markers calculated per cell type (avg. log_2_fc>1, pval adj<0.05). **(C)** Pieplot showing the 684 marker genes detected in only one cell type. Majority of unique marker genes are unknown CG and CR numbers, while the known marker genes are mostly linked to receptors. (pval adj shown as calculated by FlyMine). Insert shows the uniqueness of marker genes: 684 genes in 1 cell type, while almost all genes (∼14k) can be found as markers in multiple cell types.

**Figure S25.**
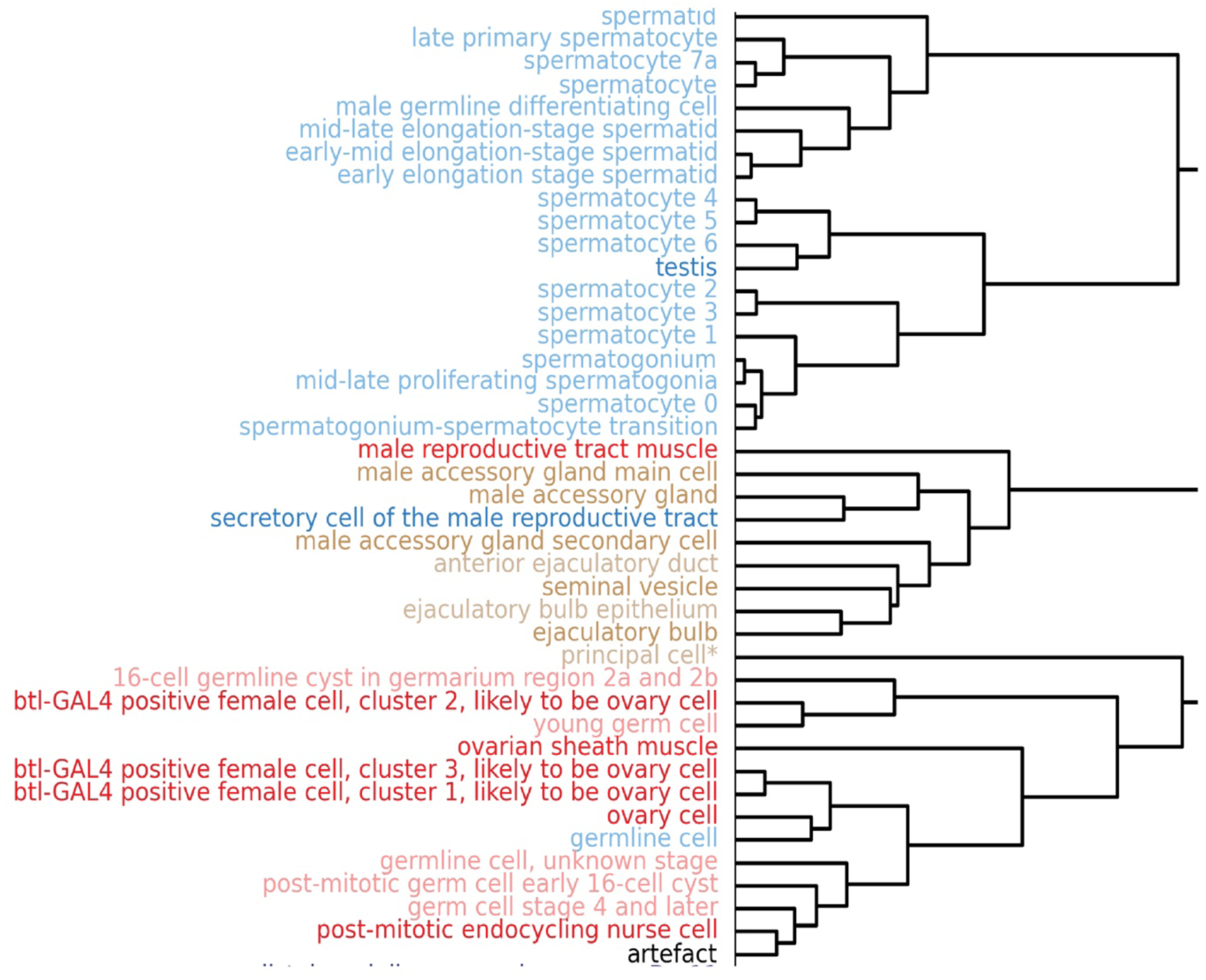
Dendrogram showing the cosine similarity of the transcriptomes of the different annotated cell types. Cell types are colored based on broad classes. Part 1 from fig. S24A.

**Figure S26.**
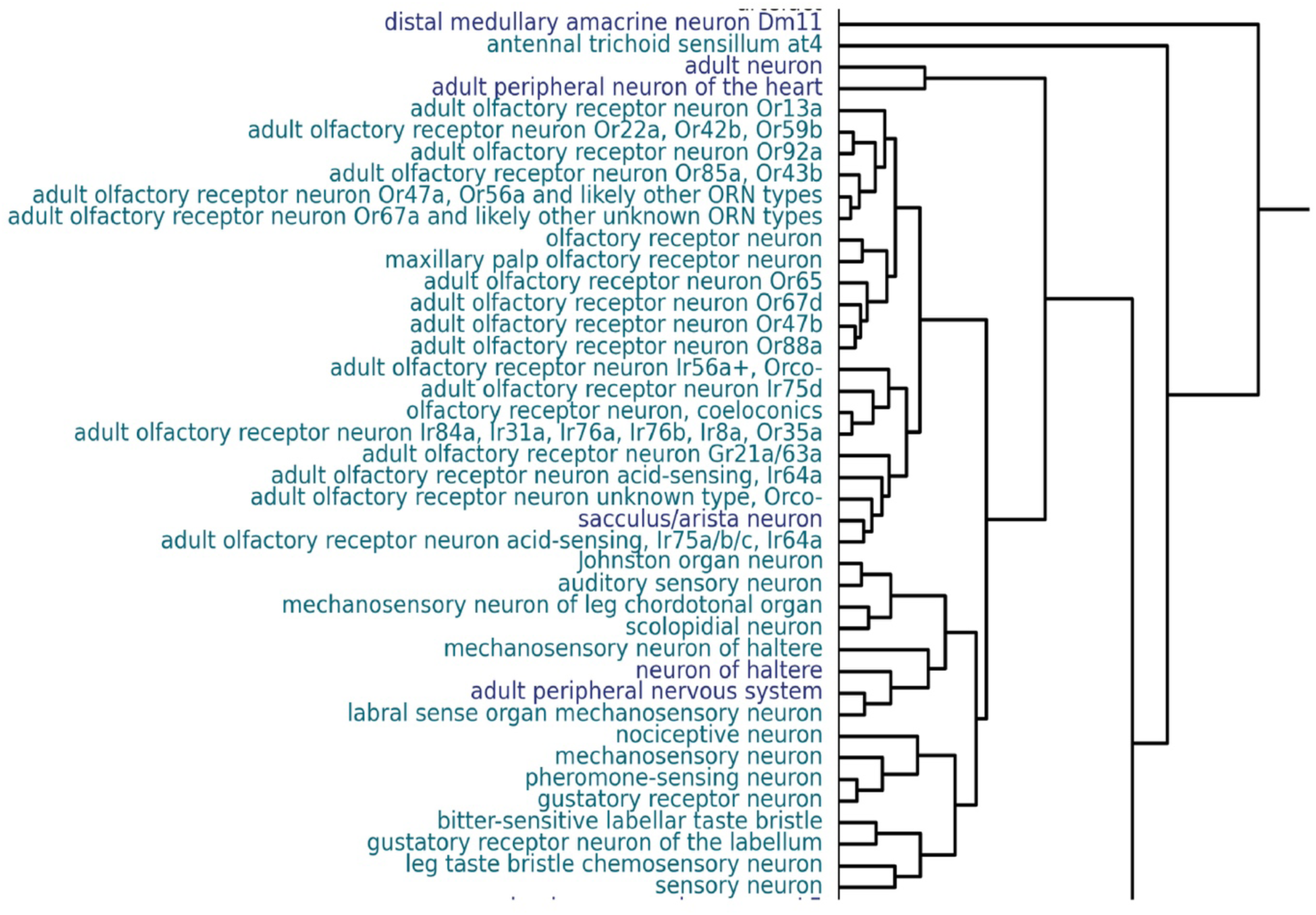
Dendrogram showing the cosine similarity of the transcriptomes of the different annotated cell types. Cell types are colored based on broad classes. Part 2 from fig. S24A.

**Figure S27.**
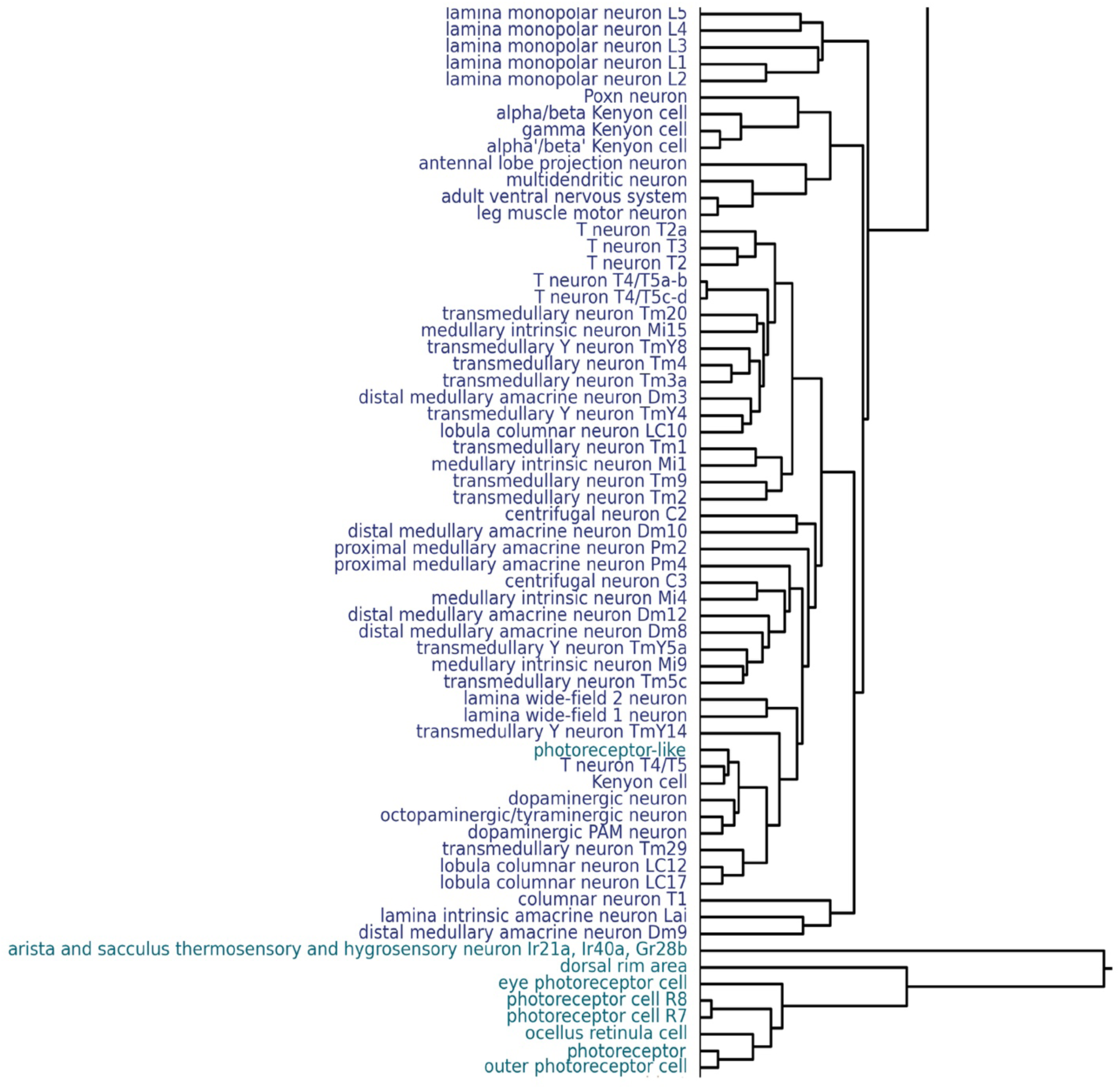
Dendrogram showing the cosine similarity of the transcriptomes of the different annotated cell types. Cell types are colored based on broad classes. Part 3 from fig. S24A.

**Figure S28.**
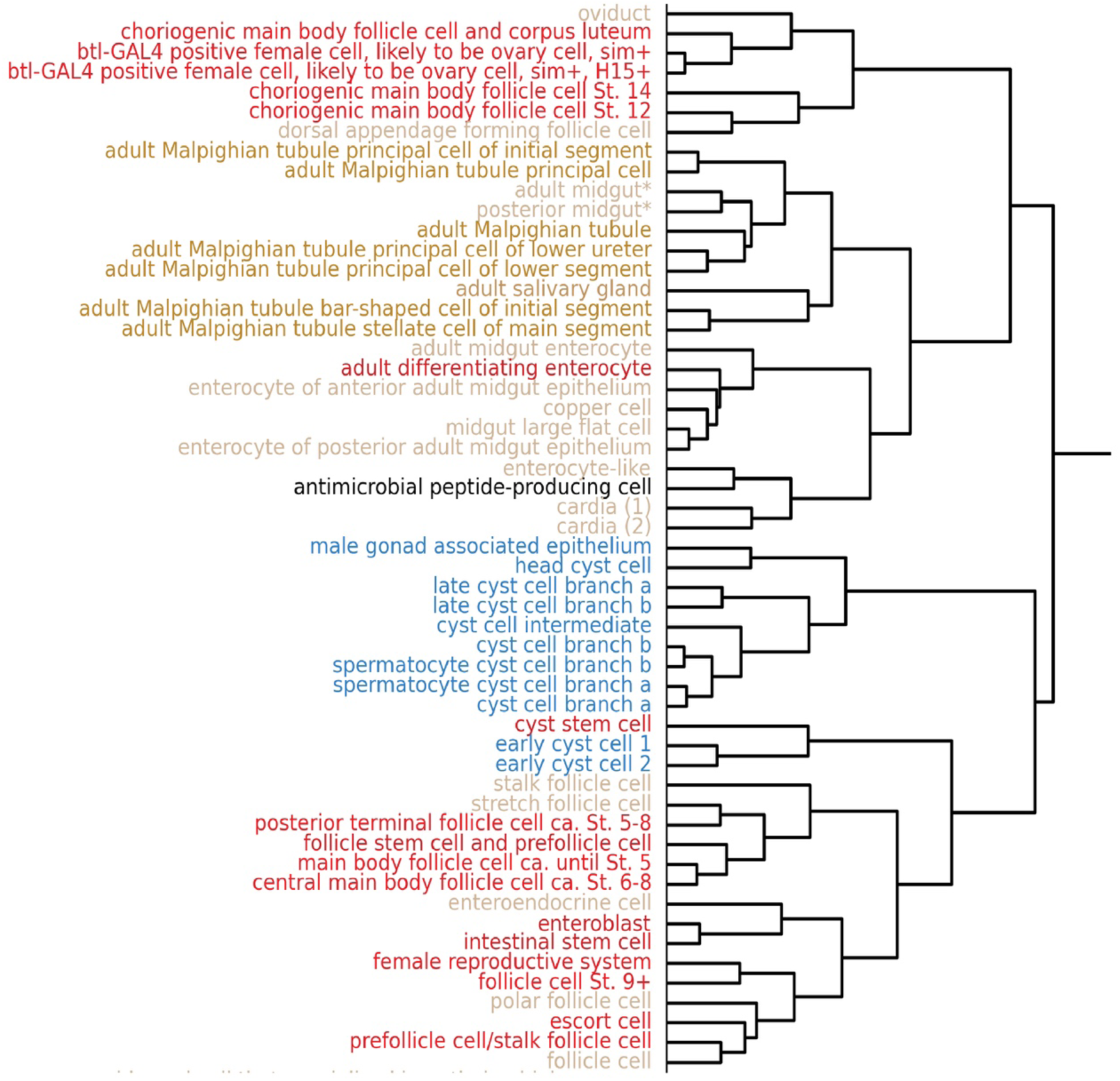
Dendrogram showing the cosine similarity of the transcriptomes of the different annotated cell types. Cell types are colored based on broad classes. Part 4 from fig. S24A.

**Figure S29.**
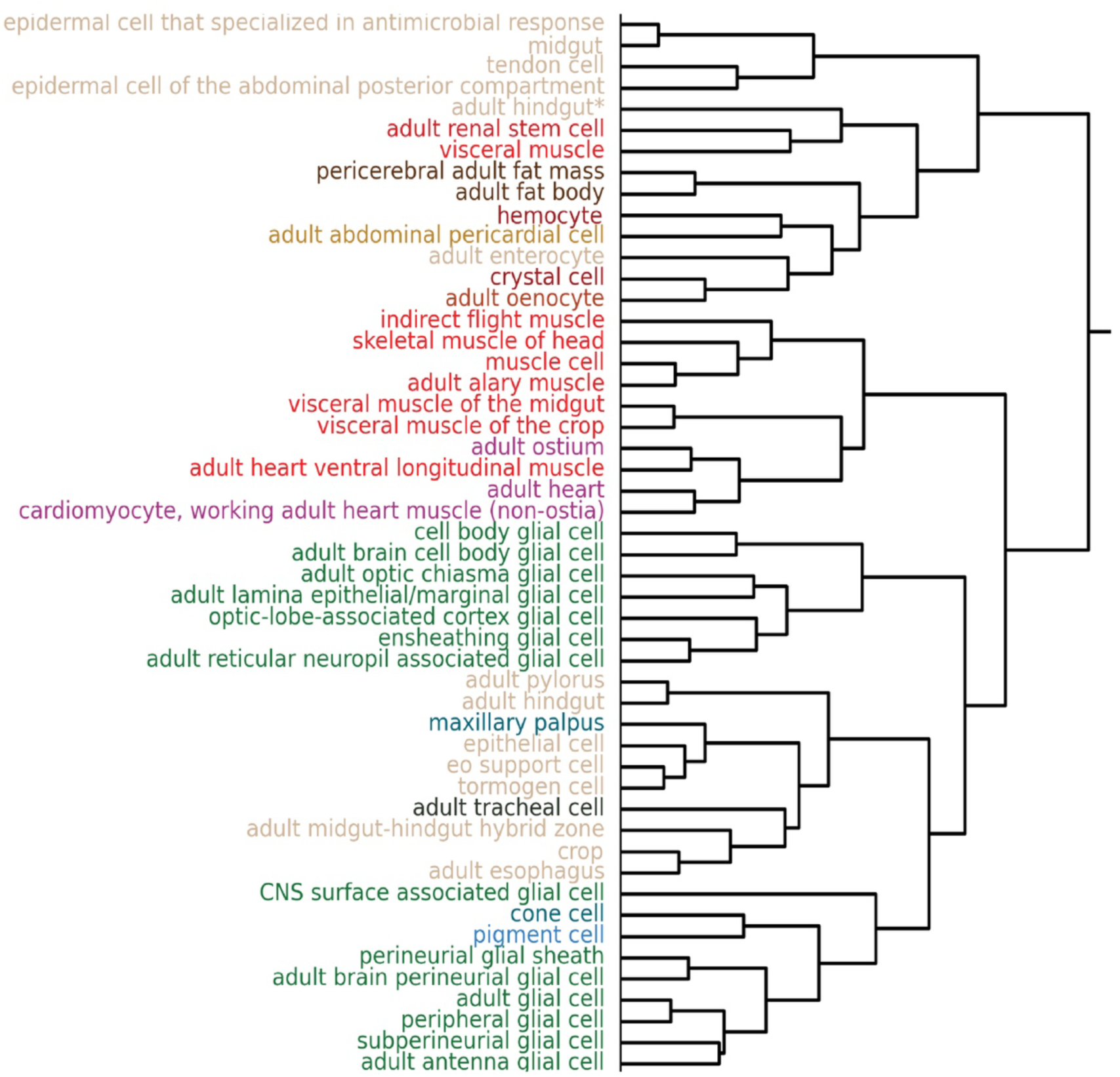
Dendrogram showing the cosine similarity of the transcriptomes of the different annotated cell types. Cell types are colored based on broad classes. Part 5 from fig. S24A.

**Figure S30.**
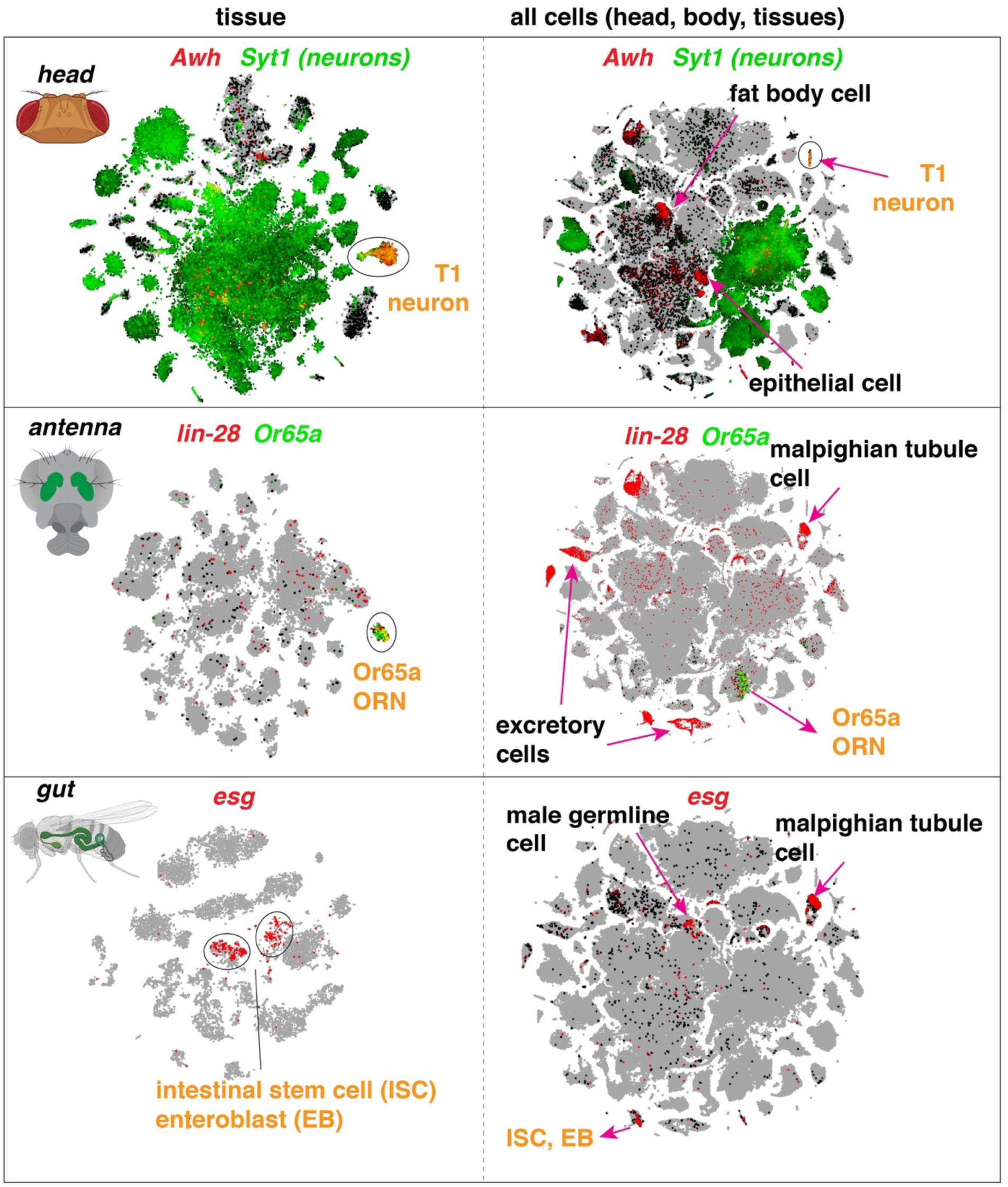
**Cell-type specific markers in individual tissues and in all cells.** Some cell type-specific markers identified in a specific tissue may have broader expression outside that tissue. *Awh* is specifically expressed in T1 neurons within the head, but also shows expression in fat body cells and epithelial cells. *lin-28* is specifically expressed in *Or65a* olfactory receptor neurons (ORNs) within the antenna, but also shows expression in malpighian tubule cells. *esg* is specifically expressed in intestinal stem cells (ISCs) and enteroblasts (EBs) within the gut, but also shows expression in Malpighian tubule stem cells and in the testis.

**Figure S31.**
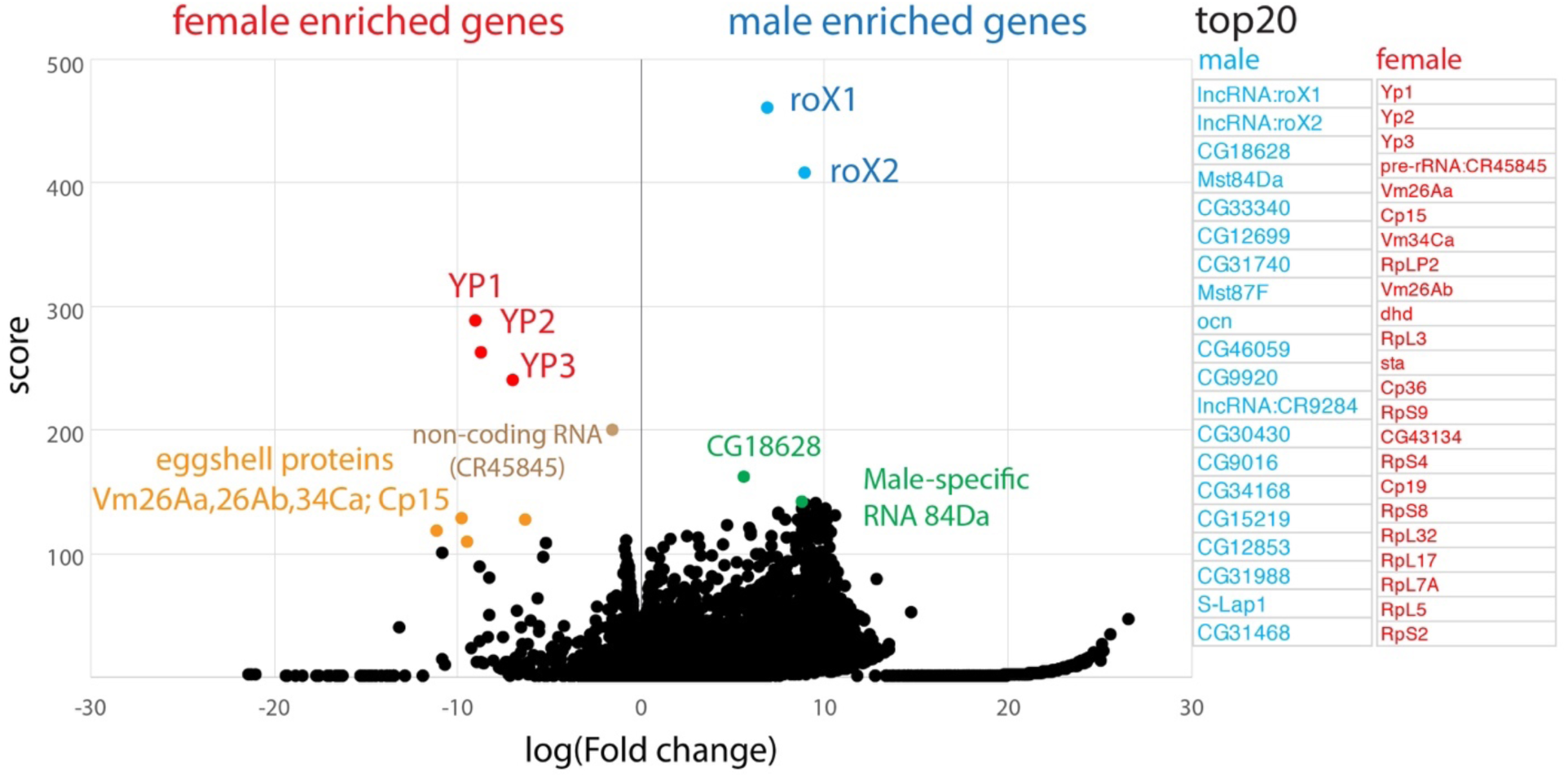
**Volcano plot showing male and female enriched genes.** Differential expression was performed between all male against all female cells in the dataset, using the Wilcoxon test in Scanpy. Score is the underlying z-value used to calculate the p-value. Fold change is in Log_2_ scale. Male and female enriched genes with top scores (20) are shown on the right. Known male specific makers (*roX1, roX2*) and female specific genes (*Yp1, Yp2, Yp3*) have the highest scores as previously seen <u>(Clough et al., 2014; Lucchesi and Kuroda, 2015)</u>, validating the quality of the data. A large number of CG genes (poorly studied or uncharacterized genes) are on the male enriched list <u>(Andrews et al., 2000; Graveley et al., 2011)</u>, suggesting their potential sex-related functions.

**Figure S32.**
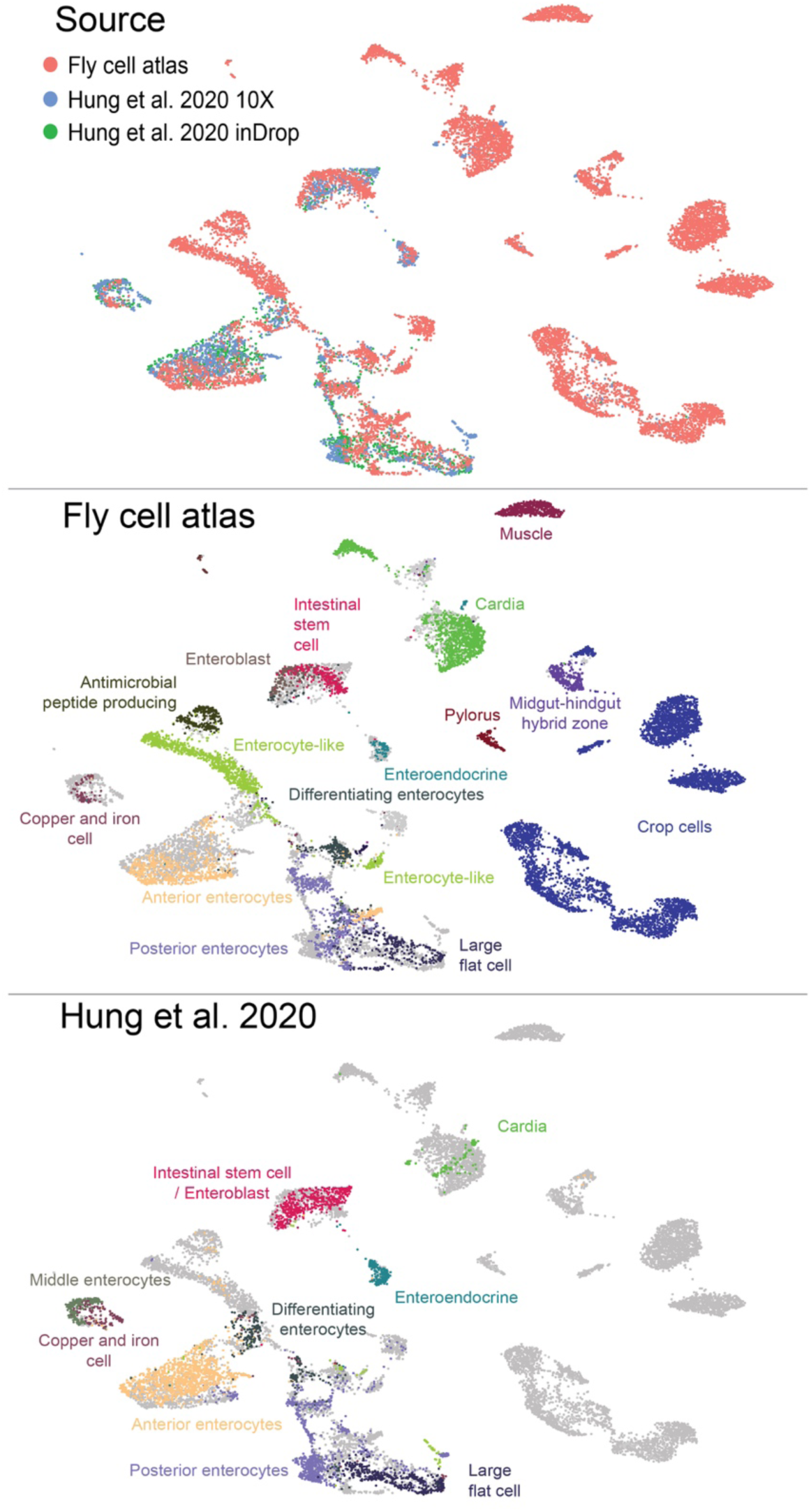
**Integration of FCA snRNA-seq data and published scRNA-seq data of the gut.** The published data are from two scRNA-seq platforms, 10x and inDrop <u>(Hung et al., 2020)</u>. Data integration was performed using Harmony <u>(Korsunsky et al., 2019)</u> using the first 30 PCA dimensions. From this analysis, we were able to identify all previously known cell types in the gut. In addition, interestingly, we were able to characterize more cell types, including visceral muscle cells and 5 subtypes of crop cells.

**Figure S33.**
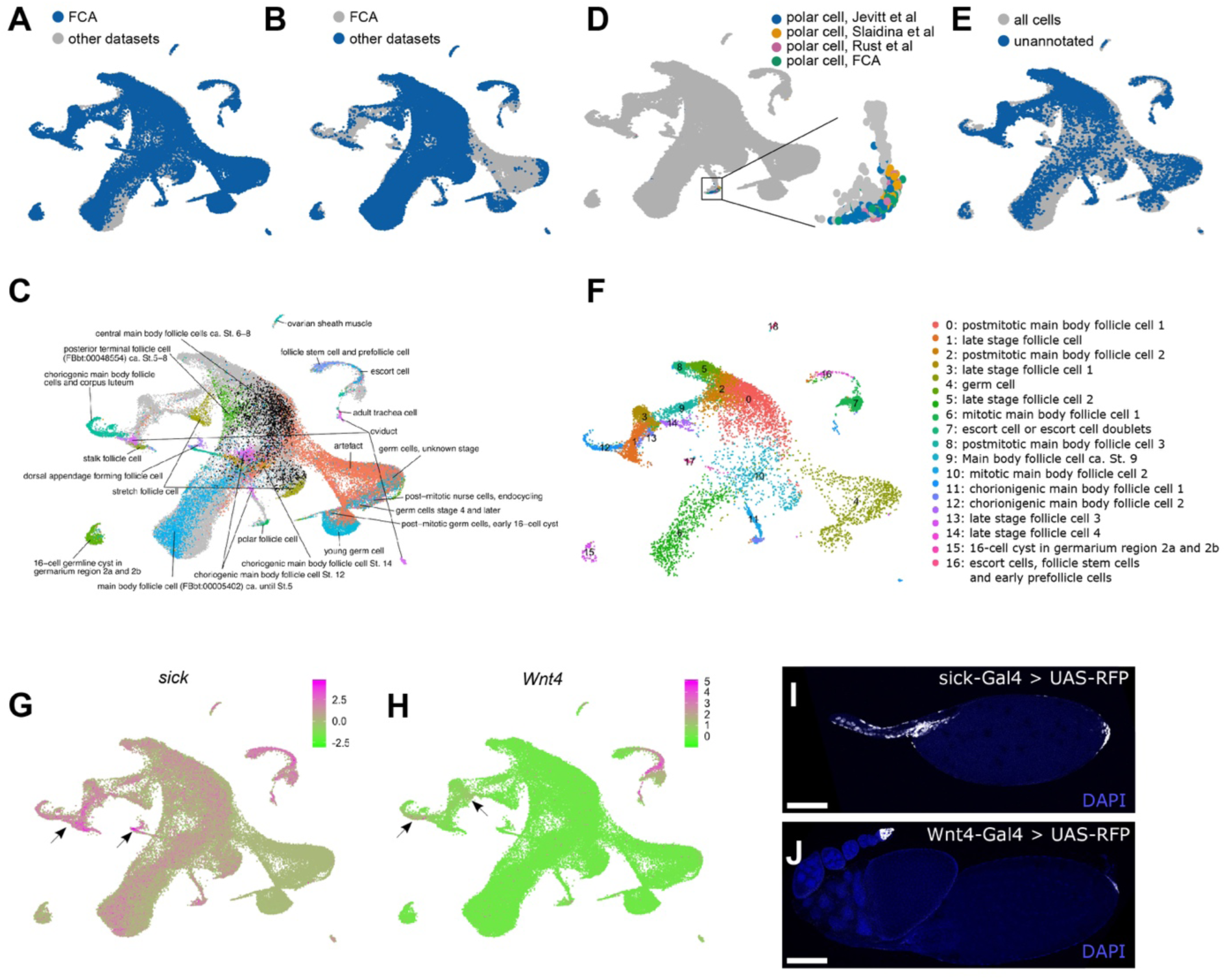
**Integration of FCA snRNA-seq data and published scRNA-seq data of the ovary** **(A)** FCA cells are highlighted in blue, and other cells are colored in gray. **(B)** Cells from the other three datasets are shown in orange, and FCA cells are displayed in gray. **(C)** Annotated FCA clusters as noted. Unannotated cells and cells from other datasets are in gray. **(D)** Polar cells identified in all datasets are highlighted and a magnified region of the UMAP plot containing polar cell clusters. **(E)** Unannotated FCA cells are labeled blue, all other cells are shown in gray. **(F)** Unannotated cells clustered independently. Presumptive cluster identities were determined by expression of marker genes as well as co-clustering with previously determined cell types. **(G, H)** Expression of *sickie (sick)* and *Wnt4* genes labeling late stage terminal follicle cells indicated by arrows. **(I, J)** Confocal images of *sick-GAL4* driving UAS-RFP showing expression in all late stage terminal follicle cells and of *Wnt4-GAL4* driving UAS-RFP showing expression in low levels in posterior terminal follicle cells and in high levels in escort cells. Confocal images are maximum intensity projections. All plots are from UMAP. Three published adult ovarian scRNA-seq datasets are from <u>(Jevitt et al., 2020; Rust et al., 2020; Slaidina et al., 2020)</u>. Datasets were integrated and batch corrected using Seurat v4.0.1. Scale bars in G and H depict average expression levels in log_2_(((UMI + 1)/total UMI)×10^4). Scale bar in I and J, 100 µm.

## Other Supplementary Materials for this manuscript include the following

Table S1: cell type-specific transcription factors for Fig. 5

Table S2: manually removed cells for sex-difference analysis in Fig. 6

Table S3: cell level data for sex-difference analysis in Fig. 6

Table S4: cluster level data for sex-difference analysis in Fig. 6

Tables S1-4 are provided as Excel files.

## FCA consortium emails and affiliations

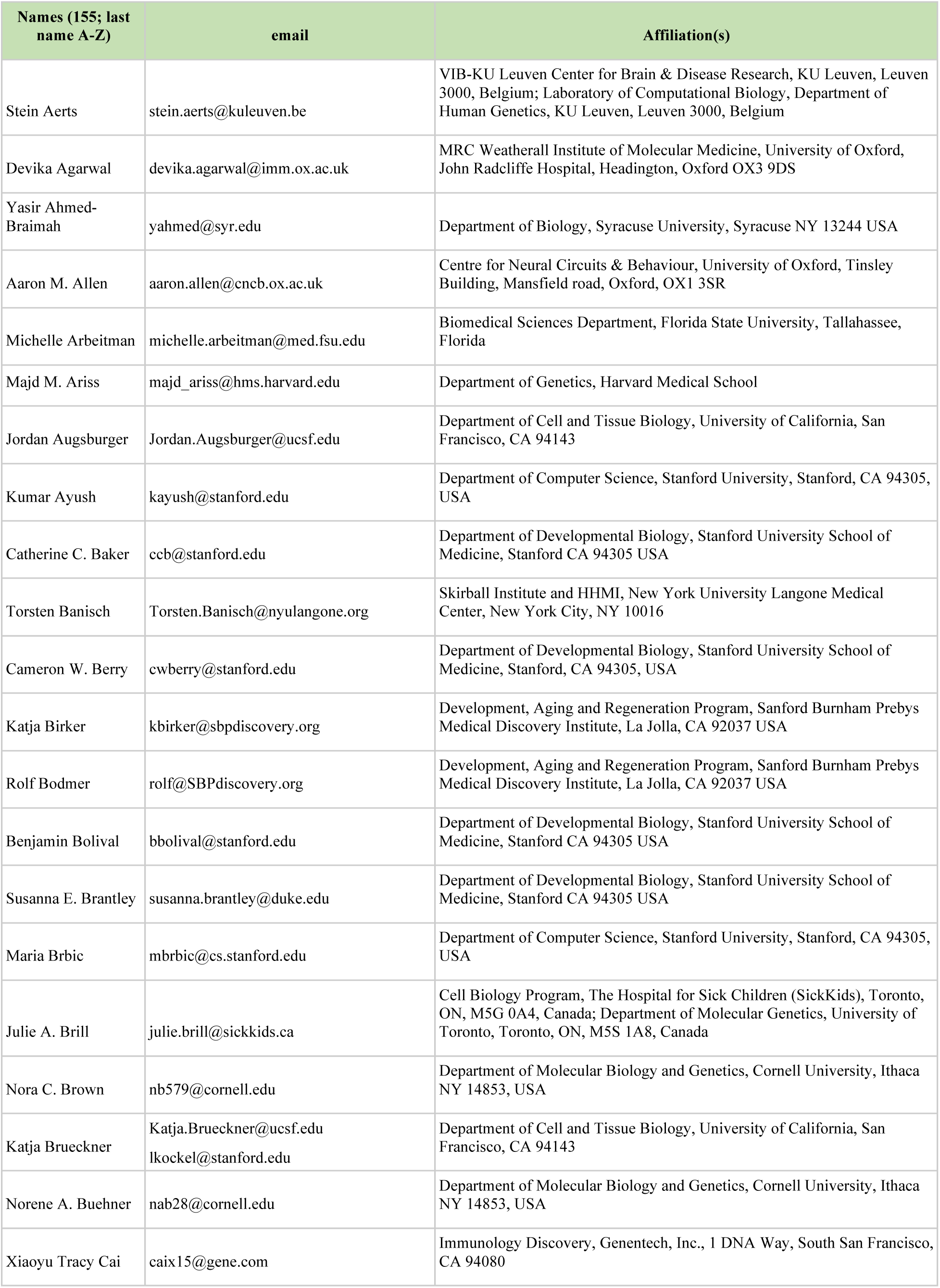
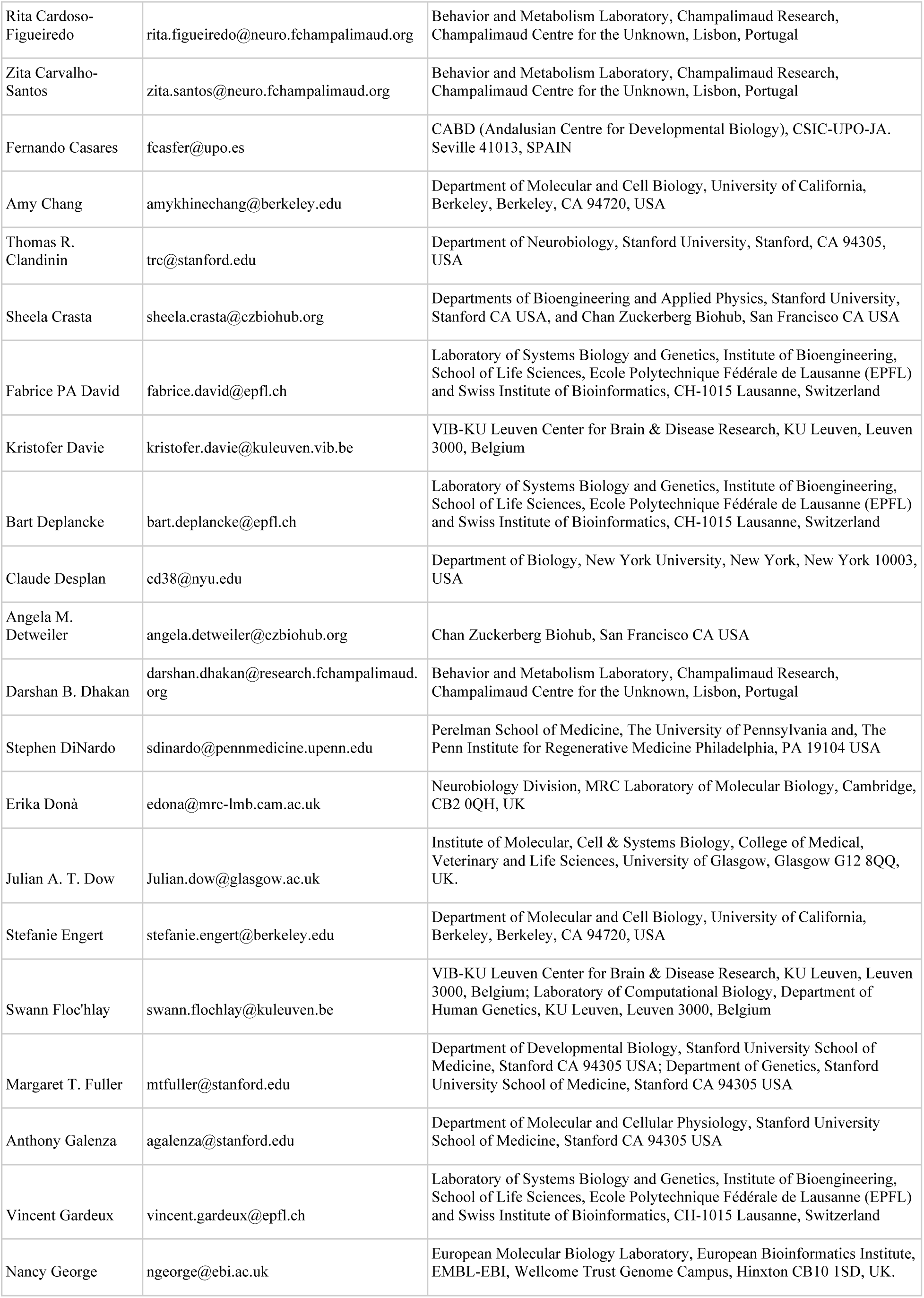
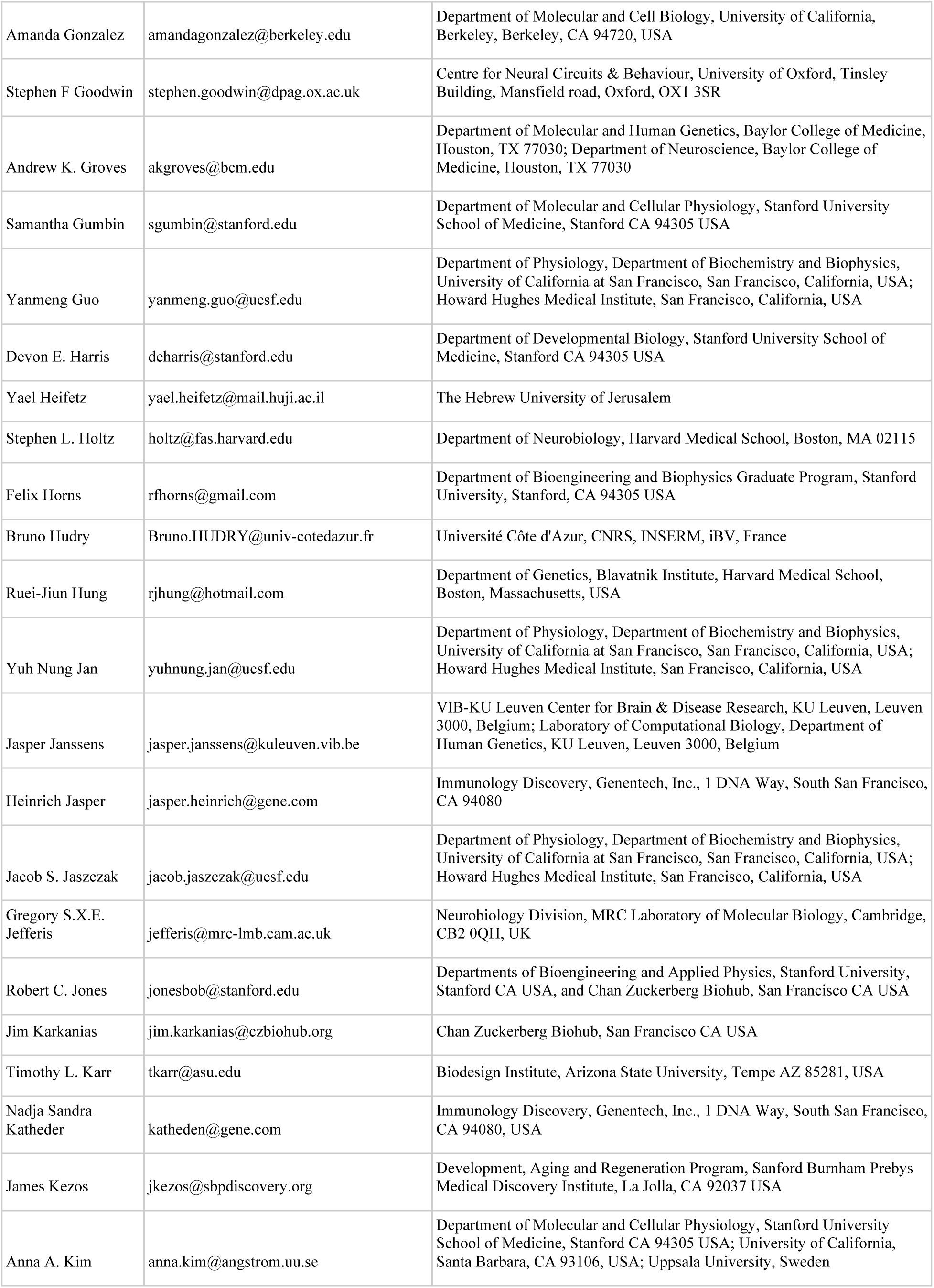
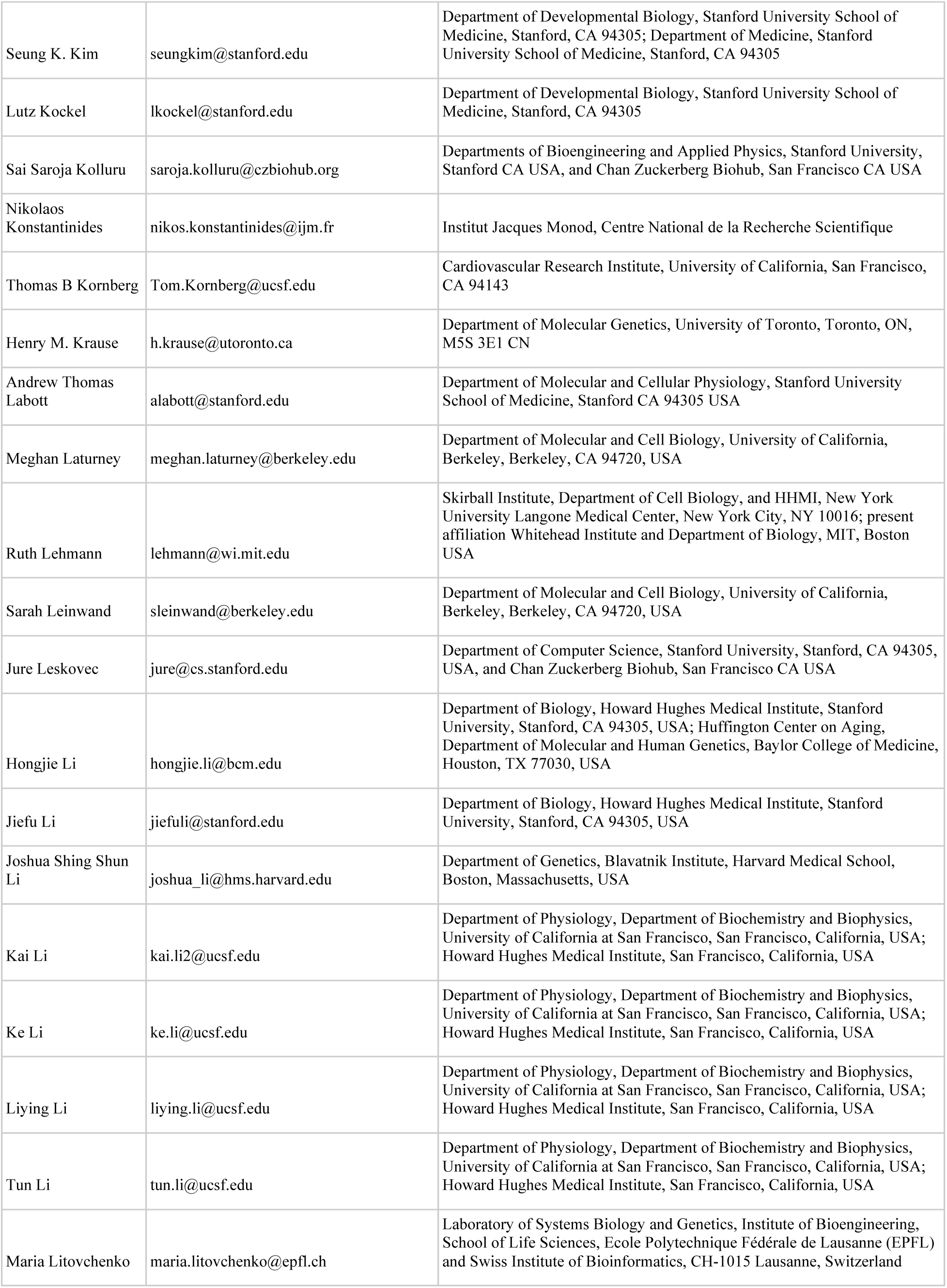
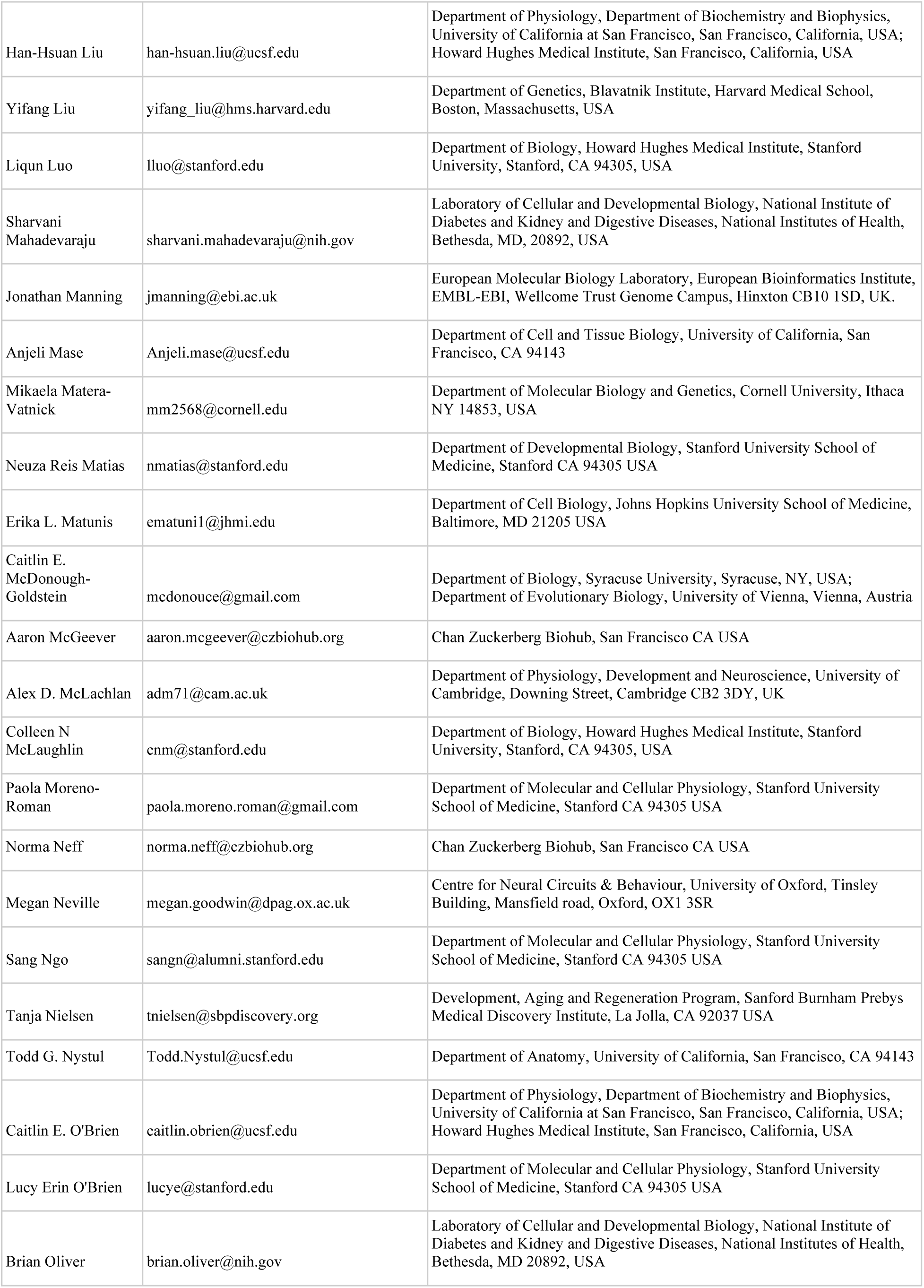
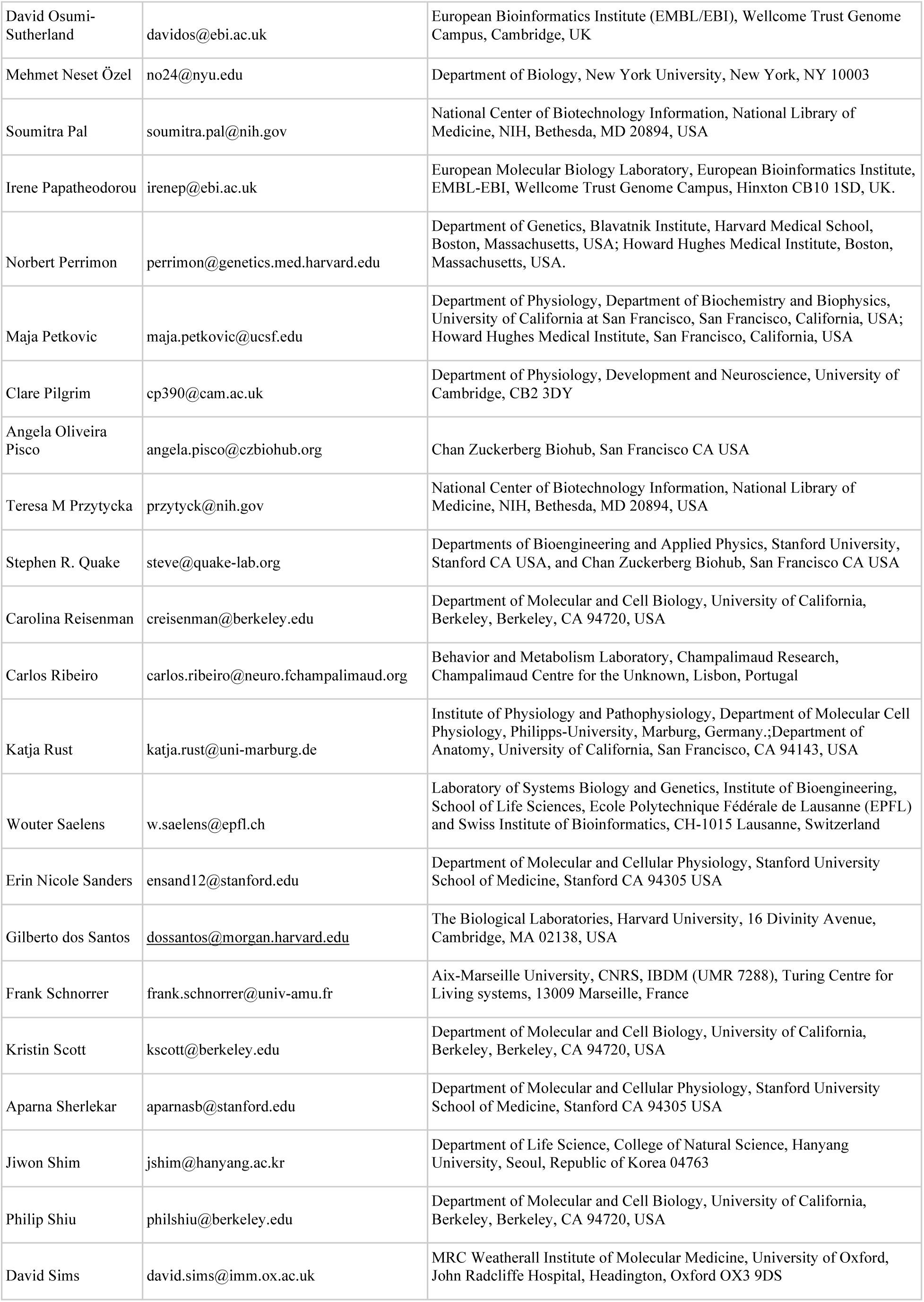
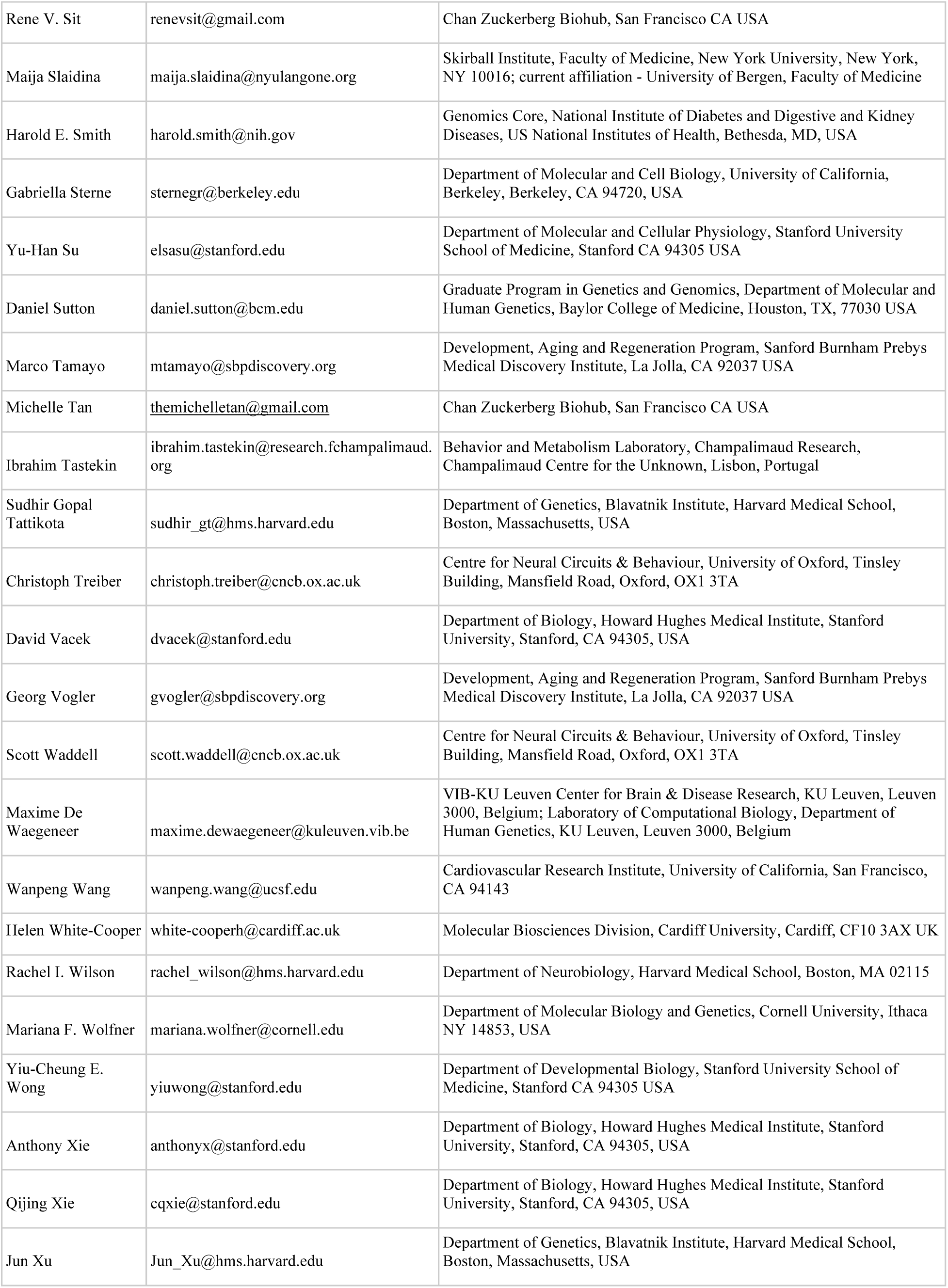
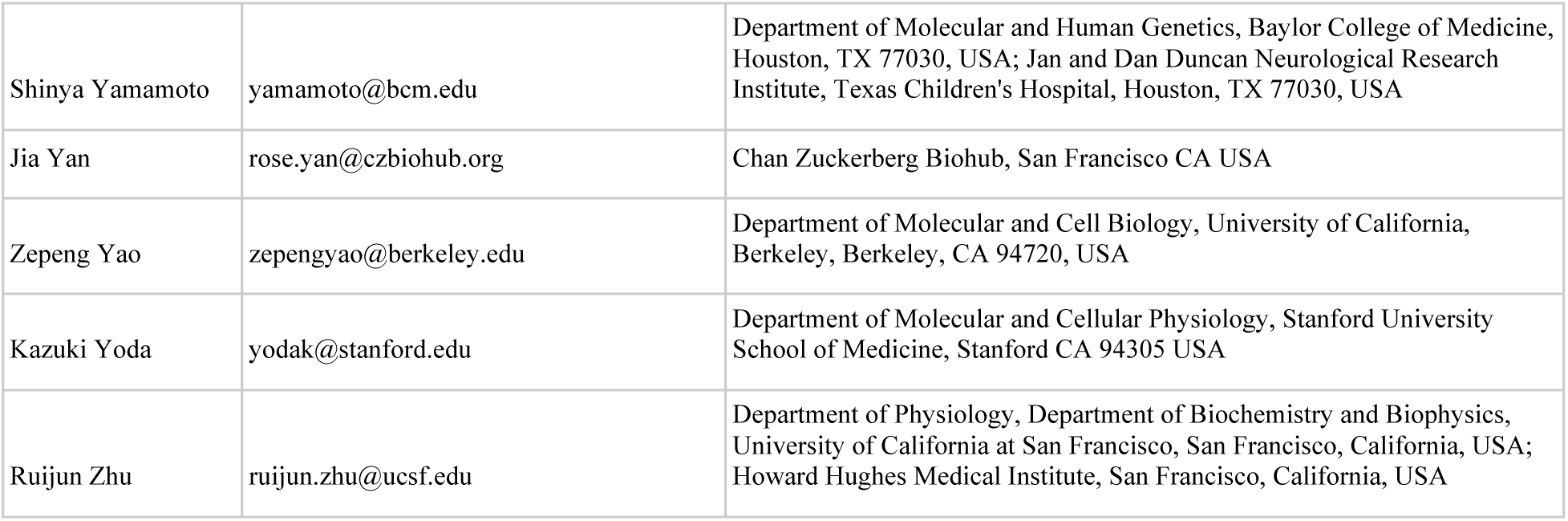

